# The catheterized bladder environment induces dysregulation of macrophage polarization exacerbating bacterial UTI

**DOI:** 10.1101/2024.07.16.603761

**Authors:** Armando M. Marrufo, Jonathan J. Molina, Chris Gager, Marissa J. Andersen, Alyssa A. LaBella, Elizabeth R. Lucas, Ellsa Wongso, Tamanna Urmi, Kassandra Arias-Parbul, Railyn Webster, Peter V. Stuckey, Kurt N. Kohler, Deborah Donahue, Victoria A. Ploplis, Matthew J. Flick, Francis J. Castellino, Felipe H. Santiago-Tirado, Ana L. Flores-Mireles

## Abstract

Urinary catheterization causes bladder damage, predisposing hosts to catheter-associated urinary tract infections (CAUTIs). CAUTI pathogenesis is mediated by bladder damage-induced inflammation, resulting in accumulation and deposition of the blood-clotting protein fibrinogen (Fg) and its matrix form fibrin, which are exploited by uropathogens as biofilm platforms to establish infection. Catheter-induced inflammation also results in robust immune cell recruitment, including macrophages (Mϕs). A fundamental knowledge gap is understanding the mechanisms by which the catheterized-bladder environment suppresses the Mϕ antimicrobial response, allowing uropathogen persistence. Here, we found that Fg and fibrin differentially modulate M1 and M2 Mϕ polarization, respectively. We unveiled that fibrin accumulation in catheterized mice induced an anti-inflammatory M2-like Mϕ phenotype, correlating with pathogen persistence. Even GM-CSF treatment of wildtype mice to promote M1 polarization was not sufficient to reduce bacterial burden and dissemination, indicating that the catheterized-bladder environment provides mixed signals, dysregulating Mϕ polarization, hindering its antimicrobial response against uropathogens.

## INTRODUCTION

<u>C</u>atheter-<u>a</u>ssociated <u>u</u>rinary tract infections (CAUTIs) are the most prevalent nosocomial infection worldwide (*1*) and often result in septicemia with 30% patient mortality (*2-8*). Patients become predisposed to developing a CAUTI by placement of urinary catheters, which are commonly used to safely remove urine from the bladder (*3*). CAUTI prevention, management, and treatment are challenging due to multidrug resistance among the CAUTI pathogens (*9-11*), resulting in increased hospital length of stay and high cost, ranging from $115 million to $1.82 billion annually (*12, 13*). Understanding the pathophysiology of CAUTIs is essential to identify new molecular targets and therapeutic strategies to prevent or treat infections, thereby reducing hospital stays, financial burden, and improving patient outcomes.

Urinary catheterization causes mechanical damage of the bladder, inducing a robust bladder inflammatory response in both mice and humans (*4, 11, 14-18*). Catheter-induced bladder inflammation and infection prompt recruitment of immune cells including monocyte-derived macrophages (Mϕs) (*5, 18-20*). However, it is unclear why despite robust immune response, CAUTI pathogens are able to evade the host antimicrobial response, colonize and persist in the catheterized bladder, and systemically disseminate (*20, 21*). Mϕs are innate immune cells that have roles in phagocytosis, cytokine production, and tissue healing and remodeling. Based on the environmental cues, such as cytokines, chemokines, and pathogen/damage-associated molecular patterns (PAMPs/DAMPs), Mϕs polarize to adopt different functional programs and are commonly classified into two states, M1 and M2 (*22-24*). However, Mϕs polarization is more complex, and these two states represent extremes ends on the polarization spectrum (*22-25*). M1 Mϕs promote inflammatory responses, microbicidal activity, and are characterized by the presence of inducible nitric oxide synthase (iNOS) for production of reactive nitrogen species (*22, 23*). In contrast, M2 Mϕs orchestrate anti-inflammatory responses and upregulate arginase-1 (Arg1) production for tissue repair and remodeling (*22, 26*).

A recent *ex vivo* study using bone-marrow derived Mϕs (BMDMϕs) demonstrated that fibrinogen (Fg) and fibrin polarize Mϕs to M1 (proinflammatory) and M2 (anti-inflammatory), respectively (*27*). Fg is a soluble protein that plays a critical role in coagulation and healing processes. Once in the damaged tissue, Fg polymerizes into insoluble fibrin clots to control bleeding (*28*). Interestingly, catheter-induced inflammation in humans and mice promotes accumulation of Fg and fibrin in the catheterized bladder and their accumulation significantly correlates with dwell time (*29-31*). However, roles of Fg and fibrin in Mϕ polarization in the catheterized bladder remain unknown.

To address this mechanism, we used two bacterial uropathogens, *Escherichia coli* and *Enterococcus faecalis,* to assess the bactericidal activity of Mϕs during CAUTI in mice carrying mutations to modulate Fg/fibrin levels or function. We found that in wildtype mice the catheterized bladder environment promotes a fibrin-dependent M2 polarization. In contrast, mice expressing mutant fibrinogen incapable of being converted to fibrin had higher numbers of proinflammatory M1 macrophages in the catheterized bladder. Notably, the fibrin-driven induction of the anti-inflammatory M2-Mϕ response outcompeted M1-Mϕ in the catheterized bladder following administration of M1-inducing cytokines. Collectively, this study reveals that Fg and fibrin exert distinct effects on Mϕ polarization and antimicrobial response during urinary catheterization. This further provides valuable information to develop therapies targeting fibrin-macrophage interactions, with the goal of changing the outcome of infection in favor of the catheterized patient.

## RESULTS

### Urinary catheterization modulates Mϕ polarization

Catheterization induces a robust innate immune response, especially neutrophils and Mϕs (*18, 19*). Yet, despite the strong immune cell recruitment, uropathogens including *Escherichia coli* and *Enterococcus faecalis*, are still able to colonize and persist in the catheterized bladder (*18-21, 31-35*). Therefore, we proposed to understand the effect of urinary catheterization in Mϕ recruitment by using uncomplicated UTI (uUTI, absence of catheter) and CAUTI mouse models in the absence or presence of infection by two of the most prevalent uropathogens, *E. coli* and *E. faecalis (4, 11)*. Furthermore, we assessed Mϕ localization and interaction with the pathogen in the bladder. Female C57BL/6 mice were catheterized or remained uncatheterized. Mice were then infected with ∼10^7^ CFUs of uropathogenic *E. coli* UTI89 or *E. faecalis* OG1RF. After 24 hours post infection (hpi), bladders were harvested for Mϕ quantification and localization by flow cytometry and immunofluorescence (IF) imaging, respectively. Uninfected and non-catheterized (naïve) bladders were used as controls.

To quantify Mϕ abundance, we gated CD45+ immune cells and identified Mϕ populations (**Fig. 1, A and B; fig. S1**, gating strategy**; and table S1**). Naïve mouse bladders showed low abundance of Mϕs (**Fig. 1B**), residing in the lamina propria (LP) (**Fig. 1C and fig. S2,** for single channels). Urinary catheterization without infection led to a significant increase of the Mϕ population when compared to naïve bladders (**Fig. 1B**). Importantly, we found that Mϕs in LP were directly interacting with Fg, which is recruited to the bladder due to the catheter-induced damage (**Fig. 1C and fig. S3,** for single channels). Differential Mϕ recruitment was observed during *E. coli* uUTI, exhibiting a significant increase over naïve bladders (**Fig. 1B)**. However, Mϕ population during *E. faecalis* uUTI was not significantly different from naïve bladder (**Fig. 1B)**.

**Figure 1.**
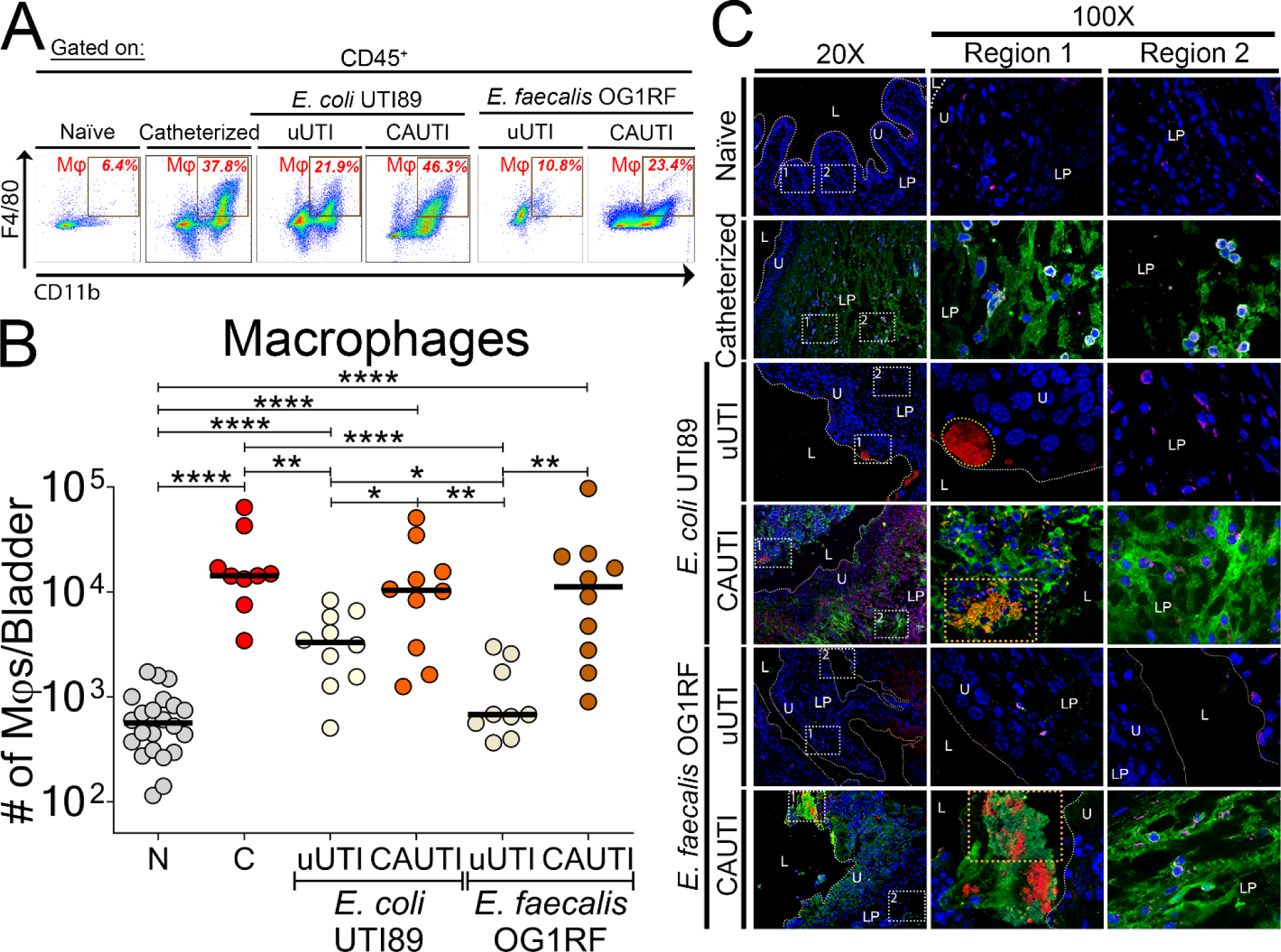
Urinary catheterization induces robust Fg/fibrin accumulation and Mϕ response. **(A)** Representative dot plots of Mϕs (Live single CD45^+^CD11b^+^F4/80^+^; **fig. S1, gating strategy**). **(B)** Quantification of number of Mϕs in the bladder. Each dot represents one mouse; horizontal lines are medians. The Mann-Whitney U test was used for determining statistical significance; *, *P* ≤ 0.05; **, *P* ≤ 0.005; ***, *P* ≤ 0.0005; ****, *P* ≤ 0.0001. **(C)** C57BL/6 female mice bladders were either naïve (N), catheterized without infection (C), or infected in absence (uUTI) or presence (CAUTI) of a catheter with either *E. coli* UTI89 or *E. faecalis* OG1RF for 24 hours. At 24 hours, bladders were stained with DAPI for cell nuclei (blue) and antibodies to detect Fg/fibrin (green), *E. coli* or *E. faecalis* (red), and Mϕs (anti-F4/80; magenta). **fig. S2-S7, individual channels**. For all representative images, white boxes at 20x represent zoomed-in areas of 100x magnification and the white broken line separates the lumen (L) from the urothelium surface (U) and the lamina propria (LP). Orange dotted rectangle depicts uropathogen-Fg biofilms.

In uUTI with either pathogen, Fg was not detected in the bladder (**Fig. 1C, fig. S4 and S6,** for single channels). During *E. coli* uUTI, the pathogen formed intracellular bacterial communities (IBCs) within the urothelium, which stimulated Mϕ recruitment (**Fig. 1C and fig. S4,** for single channels). Unlike *E. coli* uUTI, *E. faecalis* was unable to establish infection (**Fig. 1C, fig. S6 and S7,** for single channels), possibly due to the lack of Fg/fibrin accumulation during uUTIs, which has been shown to be required for biofilms formation and infection (*21, 29, 30*).

We found that significant Mϕ recruitment was observed for both pathogens during CAUTI when compared with their corresponding uUTI (**Fig. 1B**). Additionally, both uropathogens used Fg/fibrin as a biofilm formation platform (**Fig. 1C, fig. S5 and S7,** for single channels). Interestingly, Mϕs also interacted with Fg/fibrin in the LP instead of directly with the pathogen in catheterized bladders (**Fig. 1C, fig. S5 and S7,** for single channels). These data showed that the catheter-induced inflammation promotes a robust Mϕ recruitment and Mϕs interact with Fg/fibrin.

### Fibrin suppresses fibrinogen-induced M1 polarization

Previous *in vitro* reports have shown that conformational changes from the soluble Fg to its polymerized form, fibrin, alters its interaction with Mϕs (*27, 36*). Interaction with Fg induced M1-Mϕs (antimicrobial) while fibrin promoted M2-Mϕs (healing response) polarization by upregulating either inducible nitric oxide synthase (iNOS) or Arginase-1 (Arg-1), respectively (*27, 36*). Since in the catheterized bladder both Fg and fibrin are present, an understanding of their individual and co-contributions to Mϕ polarization is essential. For this experiment, we tested and optimized Mϕ polarization conditions by determining iNOS and Arg1 abundance by western blot analysis. RAW 264.7 Mϕ-like cells were exposed to different concentrations of Fg, fibrin, and M1- and M2-inducing cytokines/microbial factors (for M1: IFN-γ + LPS or GM-CSF; and for M2: IL-4). Additionally, we tested a combination of Fg and fibrin since both are available in the bladder, and as control of mixed polarization signals, we tested: 1) IFN-γ + LPS + IL-4 or 2) GM-CSF + IL-4 (**fig. S8**).

From the selected concentrations and conditions, we next treated BMDMϕs isolated from C57BL/6 and RAW 264.7 with cytokines, Fg, or fibrin for 24 hours and then assessed for non-polarized M0s (iNOS^-^Arg1^-^), M1s (iNOS^+^Arg1^-^), and M2s (iNOS^-^Arg1^+^) using IF analysis (**Fig. 2, fig. S9, and table S2 to S3,** count of each phenotype for BMDMϕs and RAW 264.7). Here, we compared changes in iNOS and Arg1 levels of Fg-or fibrin-induced Mϕs, relative to uninduced (UI), GM-CSF or IFN-γ + LPS (M1 control), IL-4 (M2 control), and GM-CSF + IL-4 or IFN-γ + LPS + IL-4 (mixed polarization signals control) induced Mϕs (**Fig. 2**). Besides M0, M1, and M2 subpopulations, both BMDMϕs and RAW 264.7 exhibited a Mϕ subpopulation that was double positive for iNOS and Arg1 (M1/M2, hybrid population), which was identified in all conditions (**Fig. 2 and fig. S9**). Therefore, the hybrid Mϕ population was analyzed in all experiments in this study.

**Figure 2.**
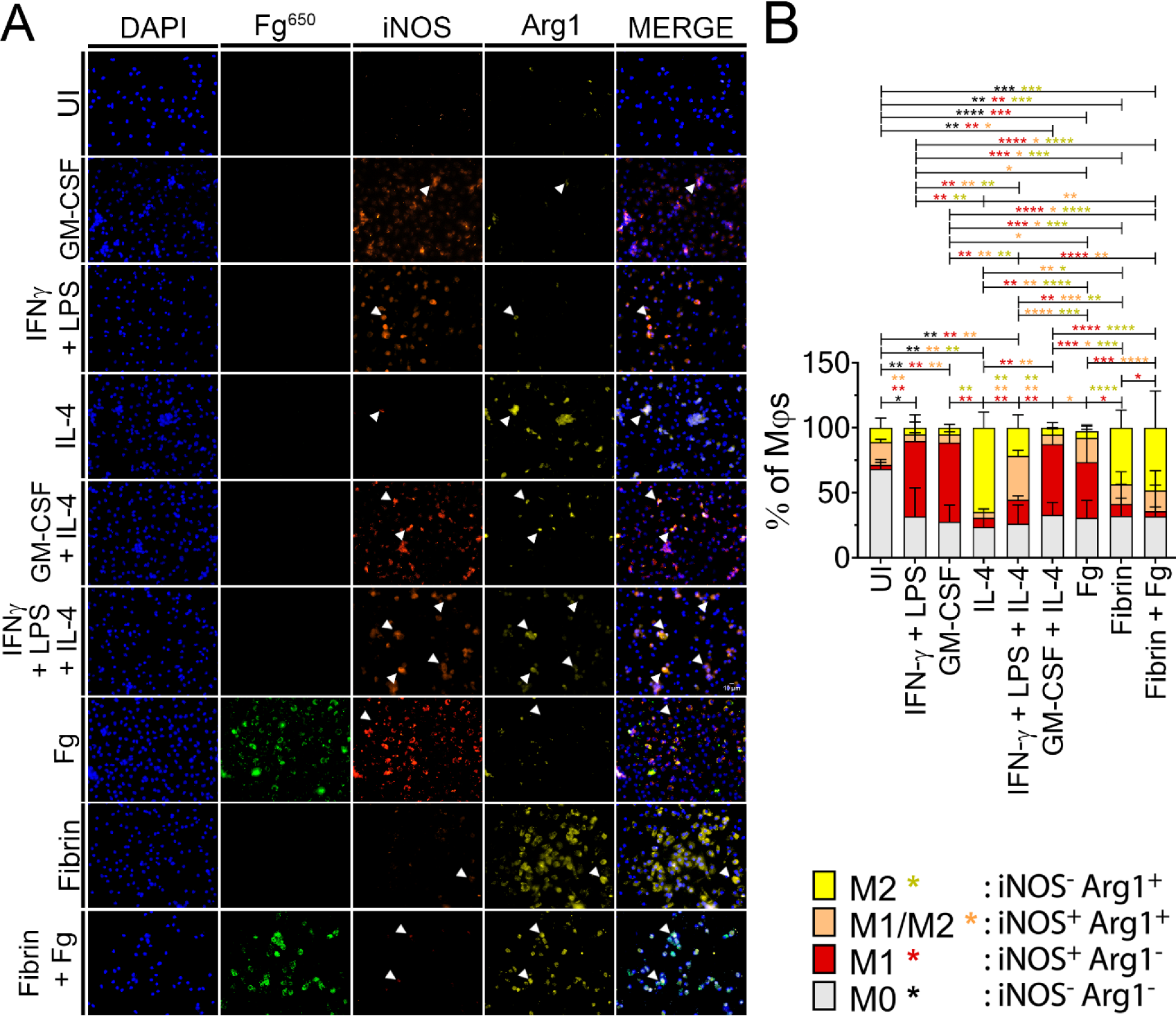
Fibrin suppresses Fg-induced M1-Mϕ activation. **(A)** BMD-Mϕs were either uninduced in DMEM cell culture media (UI) or induced with either 100 ng/ml of either GM-CSF or IL-4, both GM-CSF+IL-4, 1.5 mg/ml of either Alexa Fluor 650-conjugated Fg (Fg^650^; green) or fibrin or both (fibrin + Fg) simultaneously for 24 hours. Cells were stained with DAPI for cell nuclei and antibodies to detect iNOS (orange) and Arginase-1 (yellow) for IF analysis (representative images). Magnification is at 40x. (**B**) Percent of Mϕs that are either M0 (gray; iNOS^-^Arg1^-^), M1 (red; iNOS^+^Arg1^-^), hybrid M1/M2 (orange; iNOS^+^Arg1^+^), or M2 (yellow; iNOS^-^ Arg1^+^). Percent of Mϕs per phenotype was calculated by the number of Mϕs per phenotype divided by total number of Mϕs in each field (**table S2**, n=6-12 fields). Values represent mean ± SD. The Mann-Whitney U test was used where *P* < 0.05 was considered statistically significant; *, *P* ≤ 0.05; **, *P* ≤ 0.005; ***, *P* ≤ 0.0005; ****, *P* ≤ 0.0001.

Our BMDMϕ results showed that Fg modulates Mϕ polarization differently than fibrin (**Fig. 2A**). Fg induced significantly higher presence of M1 Mϕs (42.7%) than M2s (5.3%), M1/M2 (18.7%), or M0 (30.8%) (**Fig. 2B and table S2**). Conversely, fibrin significantly promoted M2 polarization (43.4%) over M1 (9.2%), M1/M2 (15.8%), or M0 (32.1%) (**Fig. 2B and table S2**). When Mϕs are exposed to both Fg and fibrin simultaneously, M2 (48.4%) predominates over M1 (4%), M1/M2 (15.8%) or M0 (31.8%), demonstrating that fibrin suppressed Fg-induced M1 polarization (**Fig. 2B**). This result was different from the GM-CSF + IL-4-treated Mϕs where M1 predominates while IFN-γ + LPS + IL-4 led to a majority hybrid M1/M2 population (33.7%), suggesting that GM-CSF can strongly suppress IL-4-induced M2 polarization. Similar results were found using RAW 264.7 Mϕs, except that in GM-CSF + IL-4-treated Mϕs, hybrid M1/M2 population was the most predominant population (47.0%) (**Fig. S9 and table S3**). Our data showed that Fg and fibrin exerted differential effects on Mϕ polarization, where Fg promoted M1s and fibrin promoted M2s, which may differentially affect its’ antimicrobial and wound healing response. This further demonstrated that the tissue microenvironment plays an important role in Mφ polarization and function. Additionally, BMDMϕs and RAW 264.7 Mϕs showed differential polarization profiles when GM-CSF and IL4 were used in combination, suggesting that it is critical to use a Mϕ culture model that better represents the *in vivo* models.

### Catheterized bladder environment enhances M2 Mϕ polarization

Catheterization-induced inflammation is a CAUTI hallmark characterized by reactive changes in the bladder environment (*18, 21, 29*). Previous studies have shown that acute and prolonged catheterization induce tissue damage, recruit Fg, drive fibrin polymerization, facilitate microbial colonization, promote inflammatory cytokine production, and support the recruitment of Mϕs and other immune cells (*11, 17, 21, 29, 31, 35*). Despite robust Mϕ recruitment, *E. coli* and *E. faecalis* bladder colonization and dissemination persist (**Fig. 1**, **fig. S10, and table S4**) (*17, 19, 21, 37*). The catheterized bladder environment consists of complex and diverse signals that could influence Mϕ polarization towards an antimicrobial M1 or tissue reparative M2 phenotype (*11, 18, 20, 21, 24*). To understand the effect of urinary catheterization on Mϕ phenotype, we assessed polarization by flow cytometry of Mϕs harvested from the bladders of mice with uUTI and CAUTI. Isolated single cells were immunostained for Mϕ polarization using iNOS and Arg1 (**fig. S1,** gating strategy).

Our results showed that in WT mice, urinary catheterization without infection significantly increased M2-Mϕs in number over M1-Mϕs (∼14-fold), M1/M2-Mϕs (∼5-fold), and M0-Mϕs (∼4-fold). Furthermore, M2-Mϕs predominate (∼62% of Mϕs) over M1/M2-Mϕs (∼21%), M0-Mϕs (∼9%), and M1-Mϕs (∼8%) (**Fig. 3B, fig. S10B to E, and table S5**). Differential Mϕ polarization was observed during uUTI and CAUTI with either pathogen. During uUTI with either pathogen, M1 polarization predominate over M2s with *E. coli* uUTI leading to a slightly higher M1 presence than *E. faecalis* uUTI by 2.5% (**Fig. 3B and fig. S10C**). This showed that both uropathogens promote antimicrobial M1 response in absence of a catheter. Conversely, catheter-induced inflammation during *E. coli* and *E. faecalis* infection resulted in a significant increase in number of M2-Mϕs by ∼16-fold and ∼249-fold compared to uUTIs with either pathogen, respectively (**Fig. 3B, fig. S10D, and table S5**). Although M2-Mϕs predominate in CAUTIs, we also observed a significant increase in the hybrid M1/M2-Mϕs polarization in *E. coli* and *E. faecalis* CAUTI by ∼6- and ∼4-fold, compared to uUTI with each respective pathogen (**fig. S10D)**. This indicates that the presence of each pathogen in a complex catheterized bladder environment contributes to inducing a hybrid M1/M2-Mϕs phenotype. Our results showed that Mϕs interact with Fg and fibrin during urinary catheterization and fibrin accumulates with dwell time (*31*), which suggests that fibrin accumulation may drive a more pronounced M2 response in the catheterized bladder, which could be modulated depending on the uropathogen.

**Figure 3.**
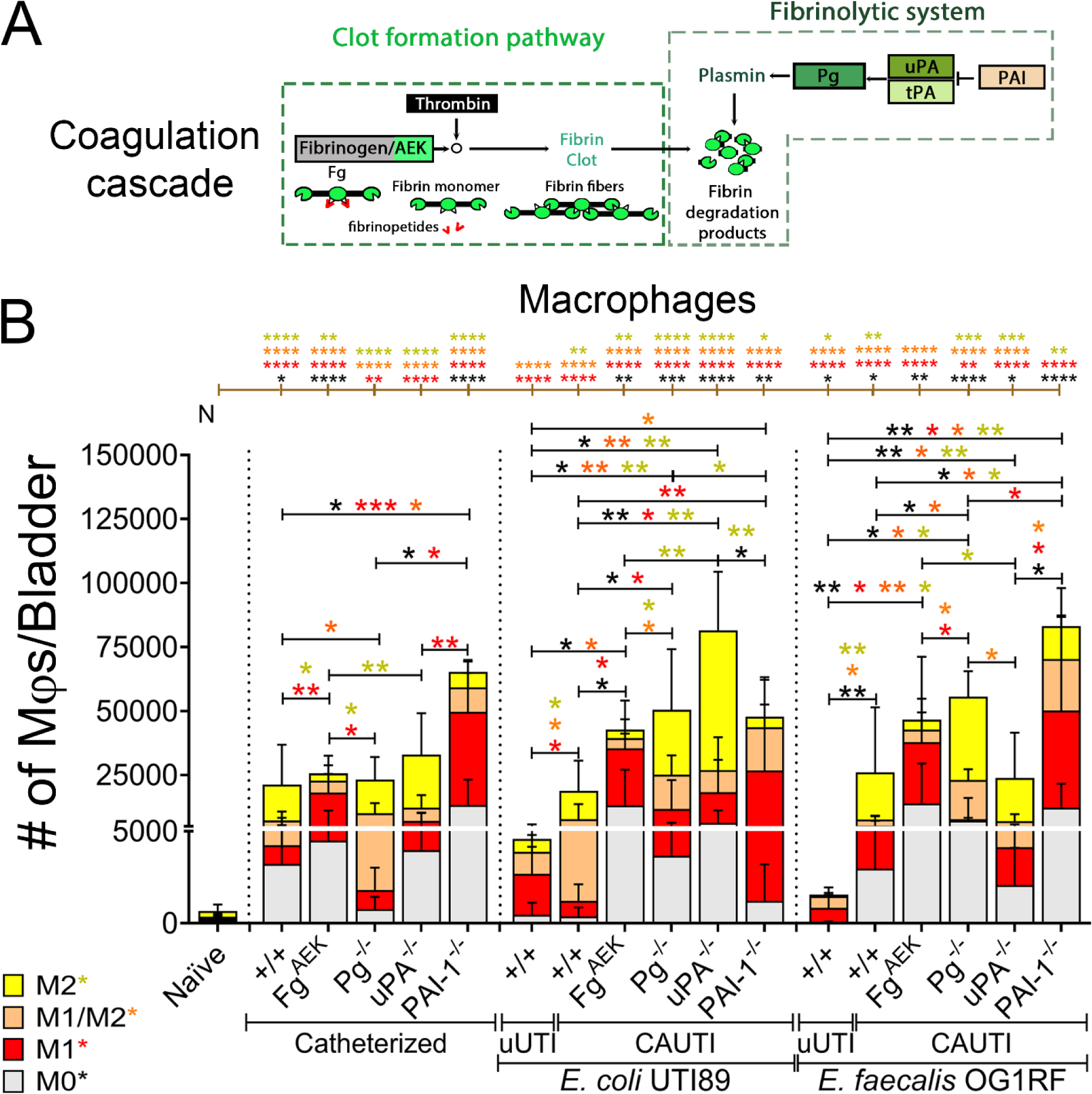
Fibrinogen and fibrin differentially polarize Mϕs during catheterization. **(A)** Coagulation cascade diagram. C57BL/6 female WT (^+/+^) and coagulation-transgenic mice bladders (**table S7**) were uninfected or infected with ∼10^7^ CFUs of *E. coli* UTI89 or *E. faecalis* OG1RF in absence (uUTI) or presence of a catheter (CAUTI) for 24 hours. Coagulation-transgenic mice used are: Fg^AEK^ (soluble Fg, no fibrin formation), Pg^-/-^ and uPA^-/-^ (elevated fibrin accumulation, no plasmin activation), and PAI-1^-/-^ (continuous fibrin clot degradation). Naïve (N) mice were neither catheterized nor infected. Bladders were then harvested and digested to isolate single cells before incubation with CD16/32 antibodies for blocking and viability dye. For flow cytometry analysis, cells were then stained with conjugated anti-mouse primary antibodies for CD45, CD11b, F4/80, iNOS, and Arginase-1. (**B**) Mϕs (CD45^+^CD11b^+^F4/80^+^) were gated from live single cell populations before further gating for M0 (Arg1^-^iNOS^-^), M1 (Arg1^-^iNOS^+^), M1/M2 (Arg1^+^iNOS^+^), and M2 (Arg1^+^iNOS^-^) populations (**fig. S1, gating strategy**). The values represent mean ± SD. Differences between groups were tested for significance using the Mann-Whitney U test. Statistical significance between each group and naïve shown as brown brackets on top of each graph. *, *P* ≤ 0.05; **, *P* ≤ 0.005; ***, *P* ≤ 0.0005; ****, *P* ≤ 0.0001. Individual data points in f**ig. S10**.

### Soluble fibrinogen in the bladder promotes M1 Mϕ polarization during catheterization

To assess the role of Fg and fibrin *in vivo*, we investigated Mϕ polarization in coagulation-transgenic mice. We recently demonstrated that soluble Fg and fibrin modulate the outcome of CAUTI (*21*). For example, catheterization of C57BL/6 mutant mice expressing a mutant form of Fg locked in the form of a monomer (*i.e.,* Fg^AEK^; **Fig. 3A**) exhibited significantly reduced *E. coli* and *E. faecalis* colonization in bladders and catheter as well as systemic dissemination (*21*). To investigate if soluble Fg exerts a distinct impact on Mϕ polarization, we again utilized Fg^AEK^ in our models of uUTI and CAUTI (*38, 39*).

Mice were catheterized in presence or absence of infection with either *E. coli* or *E. faecalis* for 24 hours. Mϕ polarization was then assessed by flow cytometry analysis (**Fig. 3B and table S5 to S6**). Compared to uninfected catheterized WT mice, uninfected Fg^AEK^ catheterized mice significantly increased M1-Mϕs in number over M2-Mϕs (4.8-fold), M1/M2s-Mϕs (3-fold), and M0-Mϕs (3-fold), which resulted in majority of Mϕs polarizing to M1s (47.7% of Mϕs) (**Fig. 3B, fig. S10B to E, and table S5**). Furthermore, this significant increase in M1-Mϕs was also observed in Fg^AEK^ mice with *E. coli* and *E. faecalis* CAUTIs. These data show that soluble Fg accumulation promoted M1-Mϕs polarization during catheterization, consistent with our prior data indicating significantly decreased bacterial burden during *E. coli* and *E. faecalis* CAUTIs in Fg^AEK^ bladders (*21*).

### Fibrin accumulation enhances M2-Mϕ polarization during catheterization

In wound healing, fibrinolysis is essential for dissolving fibrin clots to restore tissue homeostasis (*40*). Hence, defective fibrinolysis results in excess fibrin accumulation (*41-43*). Recently, we showed that plasminogen (Pg is activated to plasmin to degrade fibrin clots; **Fig 3A**) was detected on human and mouse urinary catheters (*21*). Despite the presence of Pg, fibrin accumulation persisted resulting in exacerbated colonization and dissemination during CAUTIs (*21*). Based on this, we investigated fibrinolysis in CAUTIs by focusing on Pg and its key activators on modulating Mϕ polarization by flow cytometry analysis. For increased fibrin levels, we utilized C57BL/6-background mice that were deficient in fibrinolytic factors, plasminogen (Pg^-/-^) or urokinase plasminogen activator (uPA^-/-^) (*21, 44, 45*). For reduced fibrin levels, we used mice deficient in plasminogen activator inhibitor-I (PAI-1^-/-^), which display uncontrolled plasmin activation and subsequent fibrin degradation (**Fig. 3A and table S6**) (*46*). Mice that were only catheterized or catheterized and infected with *E. coli* and *E. faecalis* for 24 hours were analyzed.

We found that M2-Mϕs polarization was predominant in fibrinolytic-mutant mice with excessive fibrin accumulation (Pg^-/-^: 52.9% of total Mϕs; uPA^-/-^: 63.0% total of Mϕs) (**Fig. 3B and table S5**). Compared to Fg^AEK^ catheterized uninfected mice, excess fibrin levels in catheterized Pg^-/-^ and uPA^-/-^ mice led to a significant increase in M2-Mϕs by ∼5- and ∼7-fold, respectively (**Fig. 3B and fig. S10E**). Furthermore, persistent fibrin accumulation also resulted in a similar robust increase in M2 Mϕs in Pg^-/-^ and uPA^-/-^ mice during *E. coli* and *E. faecalis* CAUTIs (**Fig. 3B and fig. S10E**).

Unlike *E. faecalis* CAUTI in Pg^-/-^ mice, *E. coli* infection significantly elevated the hybrid M1/M2 population by 3.3-fold from Fg^AEK^ mice infected with the same pathogen (**Fig. 3B, fig. S10D, and table S5**). Although not significant, *E. faecalis* infection also displayed a trend of higher hybrid M1/M2 population by 3.1-fold from Fg^AEK^ mice infected with the same pathogen (**Fig. 3B, fig. S10D, and table S5**). Similar to WT mice with either *E. coli* or *E. faecalis* CAUTI, mixed polarization signals from LPS on *E. coli* or lipoteichoic acid (LTA) on *E. faecalis*, as M1-Mϕs inducers, and fibrin, as a M2-Mϕs inducer, within the bladder may contribute to driving this hybrid M1/M2 phenotype. Collectively, fibrinolysis impairment resulting in continuous fibrin accumulation enhanced M2 polarization upon catheterization.

Different from Pg^-/-^ and uPA^-/-^ mice, PAI-1 deficiency significantly elevated M1-Mϕ presence in catheterized bladders. These mice display fibrin reduction and elevated production of fibrin degradation products (FDPs), due to plasmin hyperactivity continuously degrading fibrin (**Fig. 3 and table S5**). Similar to Fg^AEK^ catheterized bladders, PAI-1 deficiency significantly elevated M1 Mϕs in catheterized uninfected mice by ∼36-, ∼35-, and ∼12-fold, compared to WT, Pg^-/-^, and uPA^-/-^ catheterized uninfected mice, respectively (**Fig. 3B and fig. S10C**). Notably, *E. coli* and *E. faecalis* CAUTIs in PAI-1^-/-^ mice robustly increased the hybrid M1/M2 population by ∼2-fold, respectively, compared to PAI-1^-/-^ catheterized uninfected mice (**Fig. 3B**). These data indicate that both pathogens and reduced fibrin may also contribute to this hybrid M1/M2-Mϕs polarization. Interestingly, constitutive fibrin degradation (PAI-1^-/-^) decreased colonization and dissemination during CAUTIs (*21*), possibly due to the higher presence of M1 Mϕs contributing to the reduction in microbial burden.

In summary, soluble Fg and fibrin exhibited differential effects on Mϕ polarization in catheterized bladders. Together, this data revealed that catheterization of bladders with impaired fibrinolysis resulting in continuous fibrin accumulation shifted polarization towards M2, which may worsen CAUTI outcome. Furthermore, fibrin reduction by impairing soluble Fg to form fibrin clots or persistent fibrin degradation enhanced M1-Mϕs polarization, which may stimulate antimicrobial response during CAUTIs.

### Fibrin impairs Mϕ antimicrobial response against CAUTI pathogens

To assess *ex vivo* if Fg or fibrin directly modulates the phagocytic and bactericidal activity of Mϕs against CAUTI pathogens, we used BMDMϕs from C57BL/6 mice (**Fig. 2**) treated with GM-CSF as a control for M1-Mϕs polarization since elevated GM-CSF levels were detected in the bladder upon catheterization (*21*). To assess antimicrobial activity against phagocytized bacteria, we used an *ex vivo* gentamicin protection assay performed on cytokine-, Fg-, and fibrin-stimulated BMDMϕs (**Fig. 4**) infected with either *E. coli* or *E. faecalis* to measure antimicrobial activity.

**Figure 4.**
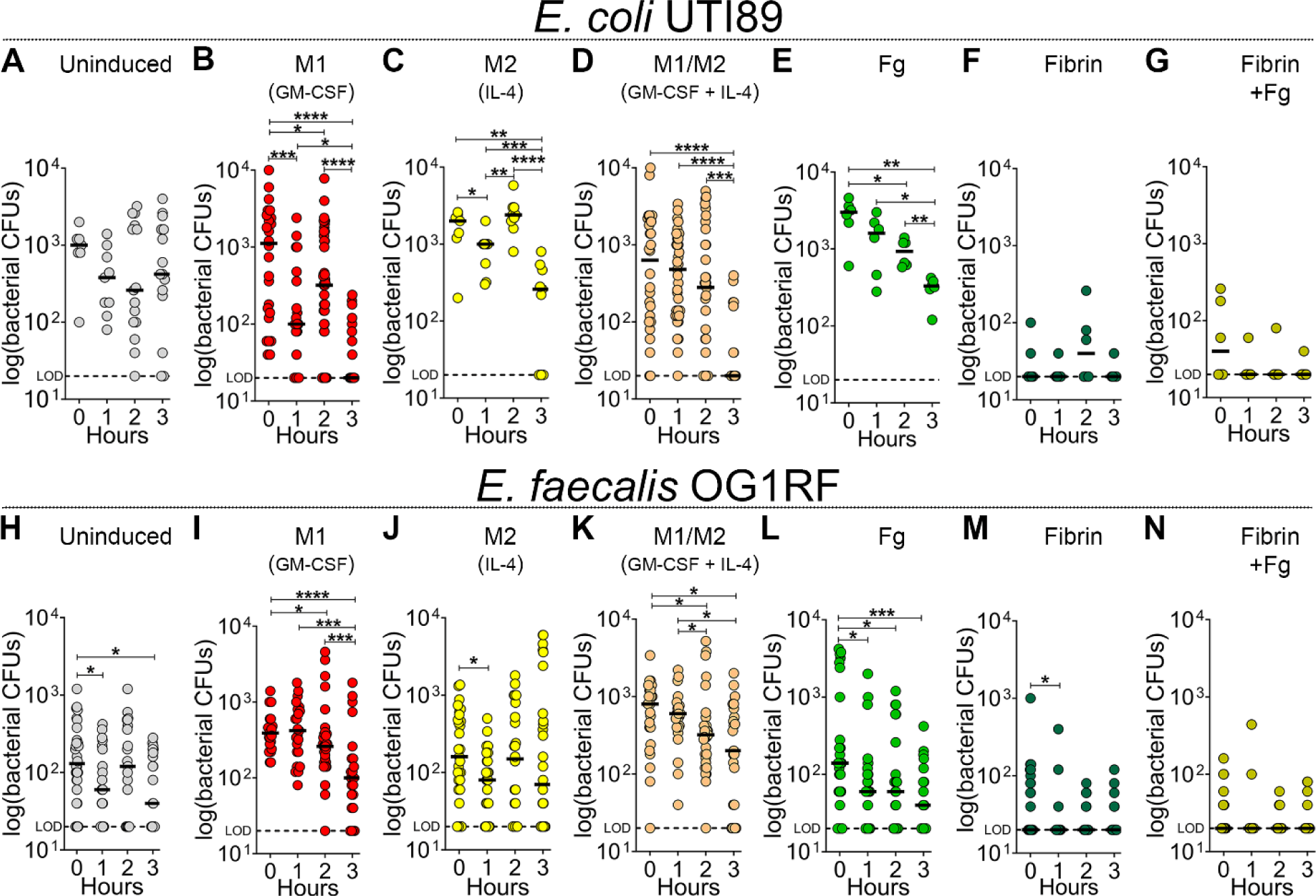
Fibrin impairs the antimicrobial response of Mϕs against pathogens. BMDMϕs were either uninduced (M0, **A & H**) or stimulated with 100 ng/mL of GM-CSF (M1, **B & I**), IL-4 (M2, **C & J**), or cotreated with GM-CSF and IL-4 (M1/M2, **D & K**), 1.5 mg/mL of Fg (**E & L**) or fibrin (**F & M**) or both simultaneously (**G & N**) for 24 hours. Mϕs were then infected with either (**A-G**) opsonized *E. coli* UTI89 or (**H-N**) *E. faecalis* OG1RF at a multiplicity-of-infection ratio of 1 (50,000 bacteria to 50,000 Mϕs) for initial uptake (45 minutes). Supernatant was removed after initial uptake and Mϕs were then treated with gentamicin for 1-3 hours to remove extracellular bacteria. Then, Mϕs were bursted with pure sterile distilled water for intracellular bacteria retrieval and CFU counts. Horizontal line represents median value. Each data point is one replicate. The horizontal broken line represents the limit of detection (LOD) of viable bacteria. The Mann-Whitney U test was used to determine significant difference between time points; *, *P* ≤ 0.05; **, *P* ≤ 0.005; ***, *P* ≤ 0.0005; ****, *P* ≤ 0.0001.

Uninduced (UI) BMDMϕs displayed higher phagocytosis of *E. coli* than *E. faecalis* by ∼8-fold (**fig. S11**). However, intracellular bactericidal response was significant for *E. faecalis* at 1 and 3 hrs post phagocytosis (hpp) (**Fig. 4H**). M1-Mϕs (GM-CSF), M1/M2-Mϕs (GM-CSF + IL-4), and M2-Mϕs (IL-4) displayed no difference in uptake of *E. coli*, compared to UI while M1-Mϕs and M1/M2-Mϕs were higher for *E. faecalis* (**fig. S11B**). M1-Mϕs and M1/M2-Mϕs exhibited antimicrobial activity at all time points against *E. coli* and for *E. faecalis* after 2 hpp (**Fig. 4, B, D, I, and K**). By 3 hpp, all M1-Mϕs, M1/M2-Mϕs, and M2-Mϕs significantly reduced intracellular bacterial burden for both pathogens (**Fig. 4**), except for M2 *E. faecalis*-infected Mϕs (**Fig. 4J**). BMDMϕs exposed to Fg showed increased phagocytosis and killing activity against both pathogens (**Fig. 4, E and L; fig. S11**). Interestingly, fibrin-exposed BMDMϕs exhibited deficient uptake and suppressed antimicrobial activity against either pathogen (**Fig. 4, F and M**; **fig. S11**). Similar to fibrin-exposed BMDMϕs, Mϕ exposure to both fibrin and Fg hindered uptake and antimicrobial activity against both pathogens (**Fig. 4, G and N**). Together, these results showed that Mϕs exhibited higher phagocytosis of *E. coli* than *E. faecalis.* Importantly, soluble Fg enhanced M1 antimicrobial response while fibrin suppressed Mϕs’ phagocytic and antimicrobial ability even in presence of Fg.

### GM-CSF outcompetes IL-4 and fibrin in promoting M1 polarization in CAUTIs

Since the catheterized bladder promotes M2-Mϕs polarization, we assessed whether Mϕ reprogramming was possible during CAUTI. Reprogramming to M1-Mϕs could be a potential therapeutic strategy to optimize Mφ response and maximize infection control while orchestrating proper healing of damaged tissue. Specifically, GM-CSF has been used as an immunostimulant in clinical studies (*47-50*). Therefore, we assessed whether GM-CSF cytokine treatment could overcome fibrin-driven programming of Mϕs to promote a switch to M1 polarization and enhanced antimicrobial activity.

Similarly to **Fig. 3B**, vehicle-treated catheterized bladder without infection significantly increased M2 and M1/M2 polarization compared with uUTIs and naïve bladders (**Fig. 5B; fig. S13D and E; and fig. S14D**). Furthermore, M1 and M1/M2 Mϕs significantly increased in vehicle-treated catheterized bladders during *E. coli* infection when compared with bladders from *E. faecalis* CAUTI and catheterized only (**Fig. 5B to D and fig. S14D**), suggesting that presence of LPS on *E. coli* counteract the M2 polarization by catheterized bladder environment (**Fig. 5C and fig. S14D**). In *E. faecalis* CAUTIs of vehicle-treated mice, M1/M2 Mϕs significantly increased compared to the vehicle-treated catheterized only (**Fig. 5D and fig. S14D).** During uUTI, the presence of either pathogen significantly increased M1 and M1/M2 Mϕs but the presence of *E. faecalis* significantly induced M2 Mφs (**fig. S14A**).

**Figure 5.**
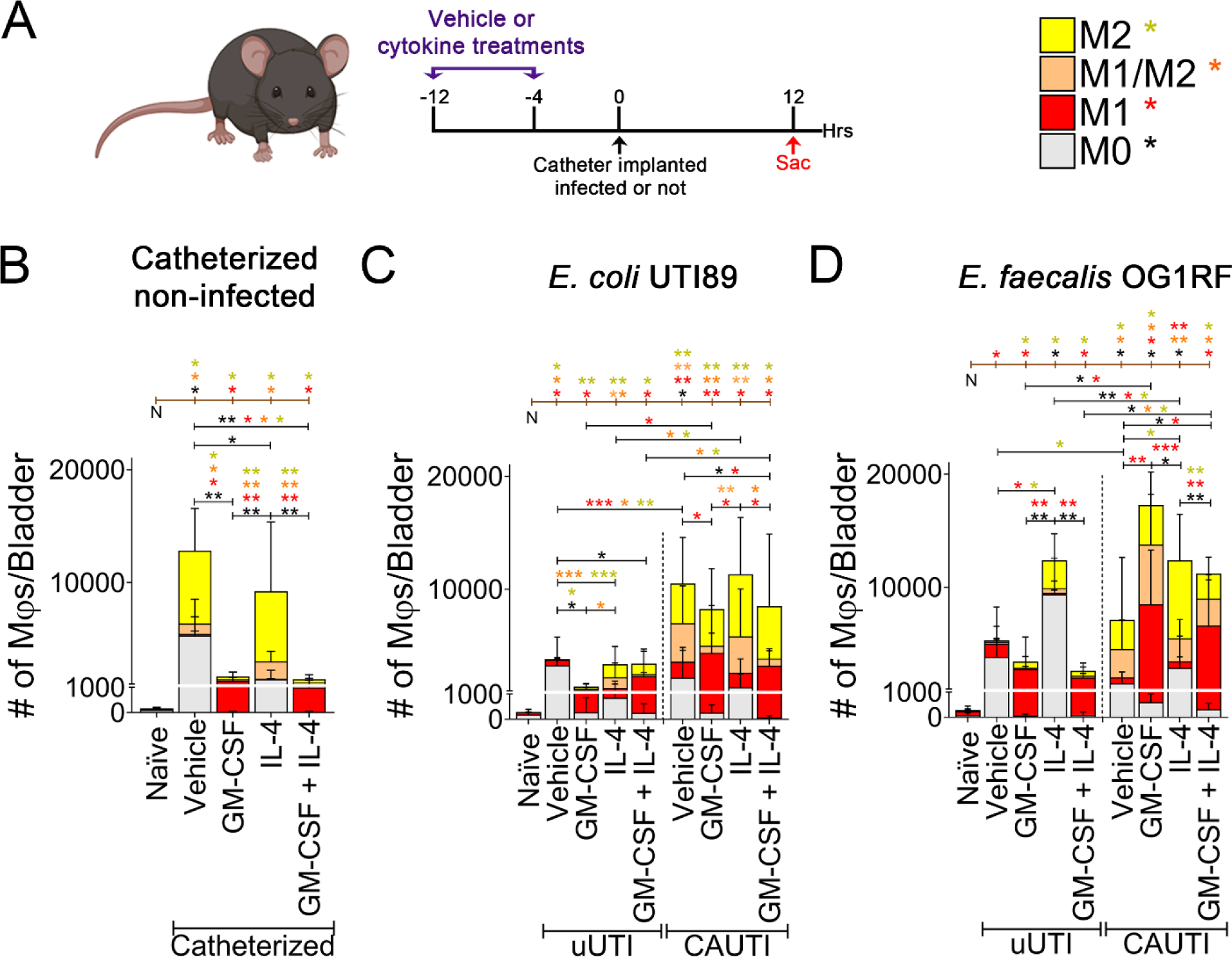
GM-CSF outcompetes IL-4 and fibrin in promoting M1 polarization in CAUTIs. **(A)** C57BL/6 WT mice were intraperitoneally (i.p.) injected with two doses of either sterile 1X PBS (vehicle), 200 ng/mouse of GM-CSF, IL-4, or co-treatment with GM-CSF and IL-4 at 12 and 4 hours prior to catheterization and/or infection. Twelve hours post-catheterization and/or infection, bladders were harvested and digested to isolate single cells for flow cytometry analysis. Cells were then stained with a viability dye and for conjugated antibodies for CD45, CD11b, F4/80, iNOS, and Arginase-1. Mϕs was gated from live single cell populations before further gating for M0-Mϕs (iNOS^-^ Arg1^-^), M1-Mϕs (iNOS^+^Arg1^-^), M1/M2-Mϕs (iNOS^+^Arg1^+^), and M2-Mϕs (iNOS^-^Arg1^+^) populations (**fig. S1, gating strategy**). Naïve (N) mice were neither catheterized nor infected. Mice were either (**B**) catheterized without infection, (**C**) infected with 10^8^ CFUs of *E. coli* UTI89 or (**D**) infected with 10^8^ CFUs of *E. faecalis* OG1RF in absence (uUTI) or presence of a catheter (CAUTI). Values represent mean ± SD. Differences between groups were tested for significance using the Mann-Whitney U test. Statistical significance between each group and naïve (N) shown as brown brackets on top of each graph. *, *P* ≤ 0.05; **, *P* ≤ 0.005; ***, *P* ≤ 0.0005; ****, *P* ≤ 0.0001. Individual data points in **fig. S13**.

Mice were treated with GM-CSF, IL-4, both cytokines, or phosphate-buffered saline (PBS) vehicle control prior to catheterization and/or infection with either *E. coli* or *E. faecalis* (**Fig. 5A**). We first examined bladder inflammation and edema by weight to understand how cytokine treatment changes the bladder environment during uUTIs and CAUTIs. Regardless of cytokine treatment and pathogen, catheterization significantly increased bladder weights compared to uUTIs (**fig. S12**). Compared to IL-4-treated mice, GM-CSF + IL-4 treatment significantly increased bladder weights during *E. faecalis* CAUTI (**fig. S12B**). This further supports that urinary catheterization causes damage and induces bladder edema. Next, we assessed whether cytokine treatment impact in Mϕ recruitment and phenotype. For all treatment groups, catheterization significantly increased total Mϕ numbers over uUTI (**fig. S13A and table S8**). However, the total Mφ population was significantly lower in the GM-CSF and GM-CSF + IL-4 groups during catheterization without infection, compared to vehicle and IL-4-treated mice (**fig. S13A**). In *E. coli* CAUTIs, there was no difference in total Mϕ recruitment between treatments (**fig. S13A**). In contrast, GM-CSF, IL-4, and GM-CSF+IL-4 treatment significantly increased the total Mφ population over the vehicle control during *E. faecalis* CAUTI by 2.4-, 1.7-, and 1.6-fold (**fig. S13A and table S8**). GM-CSF-treated mice showed significantly increased M1 polarized Mϕs in the bladder while M2 polarized Mϕs were significantly elevated in IL-4-treated mice during urinary catheterization, uUTI, and CAUTI (**Fig. 5B to D and fig. S13, C and D**). Similar to *ex vivo* BMDMϕ polarization (**Fig. 2**), GM-CSF and IL-4-cotreated mice exhibited a significant increase in M1 population, further demonstrating that (i) GM-CSF promotes M1 polarization in the catheterized bladder and (ii) GM-CSF is able to suppress IL-4-induced M2 polarization *in vivo* (**Fig. 5B to D and fig. S13C**). During *E. faecalis* CAUTI, hybrid M1/M2 population increased in GM-CSF- and GM-CSF and IL-4-treated mice (**Fig. 5D and fig. S13D**).

Our results suggest that *E. coli* presence promoted M1 polarization while *E. faecalis* promoted M2 and M1/M2 polarization (**Fig. 5B to C and fig. S14, A and D**). Additionally, urinary catheterization further promoted M2 polarization. Importantly, GM-CSF treatment prior to catheterization favored M1 polarization, suggesting that GM-CSF could induce M1 polarization, overcoming the M2 induction by the catheterized bladder environment.

### The catheterized bladder outcompetes Mϕ reprogramming by suppressing Mϕ antimicrobial activity

Next, we assessed whether reprogramming treatment modulated Mϕs’ phagocytic ability of either uropathogen during catheterization. For this, we performed the same experimental setup described in **Fig. 5A** and infected mice with either *E. coli*-or *E. faecalis*-GFP-expressing strains to quantify the number and percent of Mϕs that phagocytize the pathogen (GFP^+^ Mϕs) by flow cytometry (**Fig. 6, fig. S14, fig. S15,** gating strategy**, and table S9**). In *E. coli* uUTI, GM-CSF, IL-4, and GM-CSF+IL-4 significantly increased the number of Mϕs that phagocytize the pathogen by 3.9-, 10-, and 8.7-fold, respectively, compared to vehicle-treated mice (**Fig. 6A and table S9**). In *E. faecalis* uUTI, compared to vehicle-treated mice, GM-CSF, IL-4, and cotreatment with GMCSF and IL-4 significantly elevated phagocytosis by Mϕs by 18.6-, 14.4-, and 7.7-fold, respectively (**Fig. 6E and table S9**). Interestingly, each cytokine treatment elicited a significant increase in Mϕs that phagocytized either pathogen with catheterization (**Fig. 6, A and E**). Particularly in *E. faecalis* CAUTI, GM-CSF and GM-CSF+IL-4 enhanced uptake of the uropathogen compared to IL-4 (**Fig. 6E**). These data showed that the catheterized bladder environment promoted Mϕ uptake of both pathogens.

**Figure 6.**
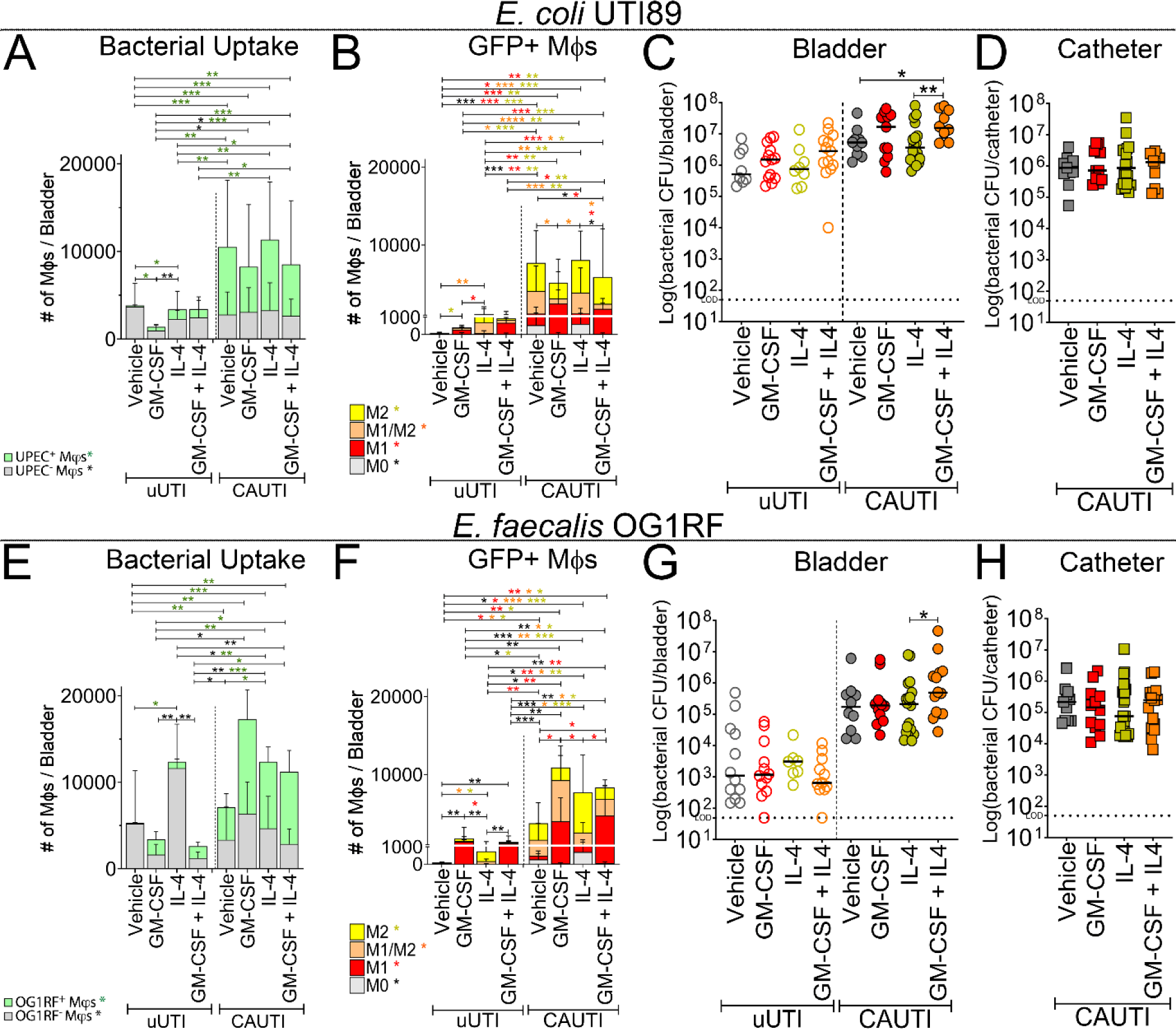
Fibrin outcompetes cytokines in suppressing Mϕ antimicrobial response during catheterization. WT-C57BL/6 mice were i.p. injected with two doses of either sterile 1X PBS (vehicle) or cytokines at 12 and 4 hours prior to catheterization and/or infection as described in Fig. 5A. Naïve (N) mice were neither catheterized nor infected. Mice were either (**A-D**) infected with 10^8^ CFUs of *E. coli* UTI89-GFP strain or (**E-G**) infected with 10^8^ CFUs of *E. faecalis* OG1RF-GFP strain in absence (uUTI) or presence of a catheter (CAUTI). Twelve hours post-catheterization and/or infection, bladders were harvested and digested to isolate single cells for flow cytometry analysis and staining as described in Fig. 5A. (**A, E**) GFP^+^Mϕs (phagocytized the pathogen, UPEC^+^ or OG1RF^+^) and GFP^-^Mϕs (did not phagocytize the pathogen, UPEC^-^ or OG1RF^-^) were each gated (**fig. S15, gating strategy**). (**B, F)** Evaluation of polarization state of GFP^+^ Mϕs populations phagocytizing the pathogen**. (C-D, G-H**) Twelve hours post-catheterization and/or infection, the bacterial burden of each strain was determined by CFU count of (**C, G**) bladders and (**D, H**) catheters. Horizontal line represents median value. Each data point is one mouse. The Mann-Whitney U test was used to test for significance between groups. *, *P* ≤ 0.05; **, *P* ≤ 0.005; ***, *P* ≤ 0.0005; ****, *P* ≤ 0.0001. The horizontal broken line represents the limit of detection (LOD) of viable bacteria. Three to four independent experiments were performed with 6-16 mice for each experimental group.

We further analyzed which Mϕ subpopulation was responsible for the uptake of the uropathogen (GFP^+^ Mϕs) (**Fig. 6, B and F; fig. S14, B and E, table S10 and S11**). During urinary catheterization, the number of M1/M2 and M2 Mϕs phagocytizing either pathogen significantly increased in all treatments when compared with uUTI (**Fig. 6, B and F; and table S10**). In both *E. coli* and *E. faecalis* uUTI, in GM-CSF and GM-CSF+IL-4-treated mice, M1 population predominantly phagocytized *E. coli* when compared with the vehicle control (**Fig. 6, B and F; and table S10**). In *E. coli* and *E. faecalis* uUTI, hybrid M1/M2s predominantly phagocyted the pathogen in IL-4-treated mice (**Fig. 6, B and F; and table S10**). During *E. coli* CAUTI, GM-CSF- and GM-CSF+IL-4-treated mice showed increased number of M1s uptaking the pathogen, representing ∼52% and ∼49% of Mϕs that phagocytized the pathogen, respectively. Of the remaining Mϕs that phagocytized *E. coli* during catheterization, approximately 25% and 38% were M2s in GM-CSF- and GM-CSF+IL-4-treated mice, respectively (**Fig. 6B and table S10**). In the vehicle- and IL-4-treated mice, M1/M2 and M2 were the main population phagocytizing *E. coli* (**Fig. 6B**). In *E. faecalis* CAUTI, GM-CSF- and GM-CSF+IL-4-treated mice, M1 and M1/M2 phagocytosis of the pathogen significantly increased when compared with vehicle- and IL-4-treated mice (**Fig. 6F**). Of these two subpopulations in this treatment, M1 comprised the majority of Mϕs that engulfed the pathogen. In the vehicle- and IL-4 treated mice, M2 were the main population phagocytizing *E. coli* (**Fig. 6F**). These data demonstrated that both the catheterized bladder and the inducer influence the Mϕ population uptaking the pathogen.

Based on these results, we investigated whether repolarization influences uropathogen colonization and dissemination (**Fig. 6 and fig. S16**). After 12-hour post-catheterization and infection, bladders, catheters, kidneys, spleens, and hearts were collected for bacterial enumeration. In uUTIs with either pathogen, there was no difference in bladder infection, regardless of the treatment (**Fig. 6, C and G**). Interestingly, GM-CSF+IL-4-cotreatment led to significantly higher dissemination into kidneys and spleen in *E. coli* uUTI, correlating with the higher bladder burden even though this increase in the bladder was not significant (**Fig. 6C and fig. S16A to B**). For *E. coli* CAUTI, bacterial bladder colonization significantly increased in the GM-CSF+IL-4-cotreatment but catheter colonization and dissemination were not significantly different (**Fig. 6D and fig. S16, A to C**). In *E. faecalis* CAUTIs, there was a significant increase in bladder colonization between GM-CSF+IL-4- and IL-4-treated mice, which also led to significantly increased bacterial dissemination into kidneys, spleen, and heart for both treatments. (**Fig. 6, G and H; fig. S16, D to F**). Together, these data demonstrated that the catheterized bladder environment promotes M2 polarization that correlates with pathogen persistence. Importantly, inducing M1 polarization with GM-CSF was not sufficient to reduce bacterial burden. Moreover, GM-CSF + IL-4 co-treatment exacerbated pathogen bladder colonization and dissemination during CAUTI with either pathogen, suggesting the mixed polarization signals may benefit the pathogen.

## DISCUSSION

A major paradox in the field is how uropathogens are able to persist during CAUTI and urosepsis (*20, 21*), despite the robust immune cell recruitment into the catheterized bladder, which includes monocyte-derived Mφs (*5, 18-20*). Here, we showed that the Mφs’ ability to clear pathogens is hindered by the complexity of the catheterized bladder environment, which modulates Mφ polarization. Our study revealed that fibrin effects on Mφ polarization overcame Fg-induced M1 polarization, resulting in a sustained M2 polarization (**Fig. 2**). Consistently, mice with coagulopathies that resulted in fibrin clot accumulation significantly promoted M2 polarization during urinary catheterization (**Fig. 3**). Importantly, we observed that Mϕ interaction with Fg promoted a bactericidal response against both *E. coli* and *E. faecali*s, while interaction with fibrin inhibited pathogen clearance *in vitro* and *ex vivo* (**Fig. 4**). This suggested that reprogramming of the Mϕs with GM-CSF could help control the infection. However, M1 reprogramming with GM-CSF treatment did not contribute to reduction of infection *in vivo* (**Fig. 5 and 6**). This indicates that the catheter-induced bladder inflammation produces mixed signals that contributes to Mϕ polarization dysfunction, resulting in suppressed antimicrobial response against infection.

Similar to our *in vitro* results, we showed that Fg and fibrin differentially modulate Mϕ polarization. In catheterized Fg^AEK^ (soluble Fg; no fibrin clot formation) and PAI-1^-/-^ (uncontrolled fibrin degradation; Fg would predominate) mice bladders, M1 polarization was prominent (**Fig. 3**), correlating with decreased microbial colonization (*21*). Furthermore, catheterized mice with impaired fibrin clot degradation (Pg^-/-^, and uPA^-/-^), where fibrin predominates, further enhanced M2 polarization when compared to WT mice (**Fig. 3**). This increase in M2 Mφs correlates with higher pathogen colonization during CAUTI and systemic dissemination in mice with coagulopathies that resulted in elevated fibrin clot accumulation (*21*). Previously, we have shown that fibrin accumulation creates ideal conditions for biofilm formation and persistence in the bladder, which is an open and dynamic system, where urine is constantly passing (*21*). However, the fact that fibrin induces M2 polarization, this may indicate that pathogen persistence during CAUTI and systemic dissemination is also, in part, due to the impairing of Mφs antimicrobial functions. Interestingly, pathogens have adapted strategies to modulate macrophage polarization to favor persistence or proliferation (*51*). For example, we have shown that *E. faecalis* uses its secreted protease, SprE, to target the fibrinolytic system, resulting in fibrin accumulation and promoting biofilm formation, consequently enhancing its colonization and systemic dissemination (*21*). Thus, by increasing fibrin accumulation, *E. faecalis* is indirectly promoting M2 polarization, making the catheterized environment advantageous for other pathogens by increasing platforms for biofilm formation and reducing immune surveillance.

We found that Mϕs stimulated with Fg exhibited higher phagocytosis and killing of pathogens while fibrin decreased phagocytosis and pathogen killing *ex vivo* (**Fig. 4**). The fact that Fg in soluble or polymerized form exerts differential effects on Mϕ polarization is intriguing. This could be due to Fg and fibrin having different structural conformations affecting its engagement and binding affinity with Mϕs receptors (*52-54*). Fg/fibrin have been reported to interact with Mϕs receptors, CD11b/CD18 (αMβ2, Mac-1, CR3), CD11c/CD18 (αXβ2, CR4), and toll-like receptor-4 (TLR-4), modulating the inflammatory response (*27, 54-57*). Hsieh *et al*. showed that BMDMϕs stimulated by soluble Fg activates proinflammatory cytokine secretion of TNF-⍺, IL-6, MCP-1, MIG, MIP-1⍺, MIP-1 β, and CCL5 similar to LPS/IFN-γ treatment, while Mϕs on fibrin matrices express IL-10, G-CSF and TGF-β1, similar to IL-4/IL-13 treatment (*27*). However, it is unclear whether Fg and fibrin uses the same receptors to differentially modulate Mϕ polarization in the catheterized bladder and how this interaction affects specific signaling pathway and transcriptional profiles.

Interestingly, a prior study has identified GM-CSF as the critical regulator of M1 antimicrobial activation in mouse models of intestinal *Citrobacter rodentium* infection and colitis while suppressing M2 wound-healing Mφ response associated with intestinal fibrosis (*58*). Despite being successful in repolarizing the Mφs based on the inducer used, when inducing M1 polarization (**Fig. 5**), *E. coli* and *E. faecalis*’s burden did not decrease in uUTI and CAUTI (**Fig. 6, C and G**). In the uUTI model, this could be explained by low Mφs phagocytosis of the pathogen (GFP^+^ Mφs) (**Fig. 6, A and E**) and their limited role in pathogen clearance in a first uUTI (*59, 60*). Even though GFP^+^ Mφs were significantly higher in all treatments during CAUTI, a large portion of Mφs were M0, M2, and hybrid M1/M2 Mφs. Based on our *ex-vivo* data showing that their killing activity was reduced compared to M1-GM-CSF Mφs (**Fig. 4)**, this suggests that their killing capabilities *in vivo* may also be impaired. Furthermore, in the catheterized bladder, the pathogen interacts with fibrin (**Fig. 1C**), which may prevent direct recognition, phagocytosis, and killing by the Mφs. Interestingly, mice treated with GM-CSF + IL-4 exhibited significantly higher bacterial burden by both pathogens and higher systemic dissemination by *E. faecalis* during CAUTI (**Fig. 6C and G, fig. S16**) and significantly higher *E. coli* kidney and spleen colonization during uUTI. This indicates that mixed polarization signals in the bladder environment may drive persistent CAUTI and systemic dissemination.

M1 and M2 Mϕs are not necessarily mutually exclusive and can coexist depending on the tissue microenvironment. Instead, Mϕs polarization occurs on a continuum towards the extreme ends of the spectrum (M1 to M2), depending on the tissue microenvironment. In our results, a hybrid M1/M2 Mϕ population was identified in the catheterized bladder (**Fig. 3**) or when combination signals were used such as GM-CSF/IL-4 or Fg/fibrin *ex vivo* and *in vivo* (**Fig. 2 to 6 and fig. S8 to S9**). This hybrid phenotype was characterized by expression of both iNOS and Arg1. This hybrid M1/M2 Mϕ-induced with GM-CSF/IL-4 exhibited an intermediate pathogen phagocytosis and killing, when compared with M1-(GM-CSF) and M2-(IL-4) (**Fig. 5B to C**). However, hybrid M1/M2 Mϕ-induced with Fg/fibrin exhibited decreased pathogen phagocytosis and killing, similar to M2-(fibrin) Mϕ. These data suggest that hybrid M1/M2 populations have a compromised antimicrobial activity, and hybrid polarization mediated by cytokines or fibrin may have different transcriptional profiles.

When comparing *in vivo* Mϕs’ capability to phagocytize uropathogen during uUTI and CAUTI by using GFP+ bacteria, we found that not only there were significantly less Mϕs but also *E. coli* and *E. faecalis* phagocytosis by Mɸs (GFP+ Mϕs) was significantly reduced during uUTI over CAUTI (**Fig. 1B**, **Fig. 6, A and E**). Other studies have shown that in mice experiencing a first time *E. coli* uUTI, depletion of tissue-resident Mϕs prior to infection does not change bacterial clearance (*59, 60*), which is consistent with the reduced bacterial phagocytosis observed in our data (**Fig. 6 and fig. S14A to C)**.

The constant catheter-induced bladder damage and inflammation (*18, 19, 21*) results in the disruption of the wound healing process and the host’s inability to resolve inflammation. Dysregulation of wound healing causes Mϕ dysfunction, leading to chronic wounds (*61, 62*). Our results showed that urinary catheterization drives M2 and hybrid M1/M2 polarization and the fact that M2 Mϕs increased in mice with coagulopathies that results in fibrin accumulation, this suggests that the catheterized bladder environment is promoting Mϕ dysfunction. Importantly, excessive M2 polarization is associated with cancers, asthma, and promoting bacterial and fungal infections (*63-66*). Consistently, prolonged urinary catheterization is correlated with increased bladder cancer risk in catheterized patients (*67-70*). Thus, this could indicate that exacerbated catheter-induced bladder inflammation and the uncontrolled M2 polarization may play a role in bladder cancer risk. However, this would require further investigation.

As our population ages, the use of urinary catheters is becoming more frequent due to increased incidence of chronic and lifestyle-related diseases, which will render patients susceptible to developing CAUTI (*2-6, 11*). CAUTIs comprise most healthcare-associated infections and has led to increased morbidities and mortalities due to urosepsis (*11*). With a high prevalence of CAUTIs and the fact that 25% of sepsis cases are from urinary isolates (*71*), it is critical to understand how the catheter-induced bladder inflammation promotes pathogen persistence despite the robust immune response. The catheterized environment provides mixed signals, which promotes dysregulating Mϕ polarization, thus impairing pathogen clearance. This study opens new avenues to understanding Mϕ polarization in the catheterized bladder by differentiating between the effect of Fg and fibrin on Mϕ antimicrobial response. It is essential to understand the catheterized bladder environment to develop therapeutics to prevent infections in patients receiving a urinary catheter.

## MATERIALS AND METHODS

### Study design

This study was performed using a preclinical CAUTI mouse model and transgenic mouse strains in accordance with approved protocols (see next subsection) to investigate how urinary catheterization affects Mϕs’ antimicrobial response in CAUTIs. Our objective was to understand how coagulopathies that results in impaired fibrin clot dissolution modulates Mϕs’ polarization and impairs its function to control bacterial infections. Having found M1, hybrid M1/M2, and M2 Mϕ subpopulations in our studies, we investigated whether soluble Fg without clot formation and fibrin accumulation that differentially polarize these subpopulations also modulates its phagocytic and bactericidal activity against both *E. coli* and *E. faecalis*. Female 6-to 8-week-old female mice were assigned to groups based on its genotype (WT, Fg^AEK^, Pg^-/-^, uPA^-/-^, and PAI-1^-/-^) and then randomly partitioned into one of these two groups first: (1) catheterized or (2) non-catheterized before being sub-grouped further into either uninfected or infected with either pathogen. Furthermore, for *in vivo* Mϕ polarization studies, mice were further subdivided into either receiving an i.p. injection of vehicle (1X PBS) or cytokines (GM-CSF, IL-4, or GM-CSF+IL4). In all experiments, a minimum of 3 and maximum of 16 mice comprise an experimental group and 3 to 4 independent experiments were performed. Sample size was determined based on previous CAUTI studies and was unchanged in this study (*17, 29, 72*). Data collection is detailed in following subsections. All data including outliers are included in statistical analysis. For quantifying Mϕ populations in flow cytometry analysis, bladders from multiple mice were not pooled together. Mice who lost catheters before time of sacrifice were excluded from all analyses. End points for mouse studies were determined before start of experiments and researchers were not blinded to experimental groups. Statistical analysis is detailed at end of this section.

### Mice handling and ethical approvals

This study utilized 6-to 8-week-old female wildtype C57BL/6 mice purchased from Jackson Laboratory, Charles River, and transgenic mice bred in W.M. Keck Center for Transgenic Research at the University of Notre Dame. All mouse studies including infections and procedures are approved by The University of Notre Dame Institutional Animal Care and Use Committee as part of protocol number 22-01-6971. All animal handling, use, and care followed approved protocol and Guide for the Care and Use of Laboratory Animals from the National Research Council (*72*).

### Bacterial strains and growth conditions

Uropathogenic *E. coli* (UPEC) UTI89 HK::GFP, *E. coli* UTI89 (no GFP), *E. faecalis* OG1RF, and *E. faecalis* OG1RF pAOJ20 (GFP) strains were used in mouse, *in vitro*, and *ex vivo* studies. UPEC UTI89 was first inoculated in 10 mL of Luria Broth (LB, MP Biomedicals). For UPEC UTI89 HK::GFP strain, LB was supplemented with the final concentration of 50 µg/mL kanamycin (Santa Cruz Biotechnologies) shaking for four hours at 37°C. After shaking, the inoculum was diluted in LB with kanamycin at 1:1000 and then grown for 24 hours in static conditions overnight at 37°C before being grown again for the next 24 hours in the same static conditions. *E. faecalis* OG1RF was grown in static conditions overnight at 37°C in 10 mL of brain heart infusion (BHI, Hardy Diagnostics) supplemented with final concentrations of 25 µg/mL rifampicin and 25 µg/mL fusidic acid (Chem-Implex). For *E. faecalis* OG1RF pAOJ20 (GFP), the inoculum was grown in static conditions overnight at 37°C in 10 mL of brain heart infusion supplemented with final concentrations of 20 µg/mL rifampicin, 20 µg/mL fusidic acid, and 10 µg/mL chloramphenicol.

### In vivo experimental model

Female mice were first anesthetized by inhaling isoflurane and then were transurethrally implanted through the urethra with a 5-mm-long silicone tubing (Braintree Scientific SIL 025) on a 30-gauge needle for 24 hours as previously described (*72*). For CAUTIs, mice were then immediately transurethrally injected with 50 µl of approximately 10^7^ colony-forming units (CFUs) of either *E. coli* UTI89 or *E. faecalis* OG1RF in 1X PBS. For uncomplicated UTIs, mice were infected as described without implantation. Naïve mice were neither implanted nor infected. After 24 hours, mice were anesthetized and then sacrificed by cervical dislocation for collection of bladders, kidneys, spleen, heart, and catheters (where applicable) for downstream applications. For bacterial enumeration, organs were homogenized, and catheters were sonicated for CFU counts.

### In vivo Mϕ polarization model

Female mice were intraperitoneally injected with two doses of either sterile 1X PBS as vehicle control, 200 ng/mouse of GM-CSF (IrvineScientific) (*73*), 200 ng/mouse of IL-4 (PeproTech) (*74*), or 200 ng/mouse of both GM-CSF+IL-4 simultaneously 12 and 4 hours prior to catheterization and/or infection with 10^8^ CFUs of either *E. coli* UTI89 or *E. faecalis* OG1RF in 1X PBS. Twelve hours post-catheterization and/or infection, mice were sacrificed for collection of bladders, kidneys, spleen, heart, and catheters for either single-cell isolation for flow cytometry analysis or bacterial CFU enumeration.

### Cell culturing

The RAW 264.7 Mϕ-like cell line (ATCC TIB-71) was cultured in Dulbecco Modified Eagle Medium (DMEM, Corning) with 4.5 g/L glucose, 2 mM L-glutamine (VWR), 1 mM sodium pyruvate (Corning), 10% fetal bovine serum (FBS, ThermoFisher Scientific), 100 units/mL penicillin and 100 µg/mL streptomycin (Corning). Cells were passaged no more than ten times. Upon isolation and separation of monocytes from bone marrow, bone marrow-derived monocytes were cultured in DMEM media supplemented with 4.5 g/L glucose, 2 mM L-glutamine, and 1 mM sodium pyruvate, 10% FBS, 100 units/mL penicillin and 100 µg/mL streptomycin, and 25 ng/mL recombinant mouse Macrophage Colony Stimulating Factor (M-CSF, IrvineScientific) for 5-7 days before use. Cells were maintained in 37°C 5% CO_2_ incubator before use.

### Isolation of bone-marrow derived monocytes

Bone marrow from femurs of 6-to 8-week-old female C57BL/6 mice were flushed with Roswell Park Memorial Institute (RPMI) 1640 media with 1X L-glutamine supplemented with 5.95 g/L HEPES (VWR), and 1 mM sodium pyruvate to retrieve all cells. Peripheral blood mononuclear cells (PBMCs) were separated from red blood cells and neutrophils using centrifugation separation with density gradients Histopaque-1077 (Sigma-Aldrich) and Histopaque-1119 (Sigma-Aldrich). After centrifugation, PBMCs were then isolated, washed with 1X DPBS, and treated in DMEM media supplemented with 4.5 g/L glucose, 2 mM L-glutamine, 1 mM sodium pyruvate, 10% FBS, 100 units/mL penicillin and 100 µg/mL streptomycin, and 25 ng/mL recombinant mouse M-CSF for 5-7 days for differentiation into Mϕs.

### Fibrin gel formation

For forming fibrin gels, soluble human Fg (Enzyme Research Laboratories) was treated with 2 U/mL of thrombin (Sigma-Aldrich) at 37°C for one hour.

### In vitro and ex vivo polarization of Mϕs

Uninduced (M0) RAW 264.7 and BMDMϕs were cultured in its respective media without cytokine treatment. For M1 polarization, cells were treated with 100 ng/mL GM-CSF (IrvineScientific) or IFN-γ (R&D Systems) and LPS for 24 hours. For M2 polarization, cells were treated with 100 ng/mL IL-4 (PeproTech) for 24 hours. To obtain a hybrid M1/M2 phenotype, cells were induced with 100 ng/mL of GM-CSF and IL-4 or 100 ng/mL of each LPS, IFN-γ, and IL-4 for 24 hrs.

### Immunofluorescent staining of RAW 264.7 and BMDMϕs

For IF analysis, RAW 264.7 or BMDMϕs were stimulated with the listed concentrations for 24 hours as described in the preceding section. For each condition, approximately 500,000 Mφs were seeded on glass-bottom petri dishes for 24 hours before staining. For Fg, Mϕs were resuspended in 1.5 mg/mL of soluble Alexa Fluor 650-conjugated Fg (Fg^650^, Invitrogen). To form fibrin gels on dishes, the glass bottom was first coated with 1.5 mg/mL of unconjugated Fg before polymerization with 2 U/mL of thrombin as described previously. For Mϕs simultaneously interacting with both fibrin and Fg, the glass bottom was first coated with the 1.5 mg/mL fibrin gel then Mϕs were resuspended in cell culture media containing 1.5 mg/mL of soluble Alexa Fluor 650-conjugated Fg. After a 24-hour incubation, Mϕs were first fixed with 10% formalin, then washed in 1X PBS three times before permeabilization with 0.3% Triton X-100 for 20 minutes. Cells were then blocked with 1X PBS with 1% BSA and 0.3% Triton X-100 for 1 hour before incubation with rabbit anti-iNOS (1:100) & goat anti-Arg1 (1:100) primary antibodies at 4°C overnight. After incubation, cells were washed in 1X PBS three times then incubated with DyLight 488 donkey anti-goat and DyLight 550 donkey anti-rabbit secondary antibodies (1:500) at room temperatures for two hours. Sections were then washed with 1X PBS before staining with Hoeshst dye (1:10,000) for imaging.

### Immunofluorescent staining and imaging of bladders

Bladders were harvested and fixed in 10% formalin overnight at 4°C, before being processed and sectioned for staining as previously described (*33*). Sections were deparaffinized with xylene, rehydrated with isopropanol, and washed with water. Antigen retrieval was done by boiling the sections in 10 mM sodium citrate, rinsed in water, and then washed in 1X PBS. Sections were then blocked with 1X PBS with 1% BSA and 0.3% Triton X-100 for 1 hour before incubation with corresponding primary antibodies at 4°C overnight. One set of bladders were stained with goat anti-Fg (1:100), rabbit anti-*E. coli* (1:100) or anti-Group D *Streptococcus* (1:100), and rat anti-F4/80 (1:100) antibodies. Another set of bladders were stained with rabbit anti-iNOS (1:100), goat anti-Arg1 (1:100), and rat anti-F4/80 (1:100) antibodies. After incubation, sections were washed in 1X PBS three times then incubated with corresponding secondary antibodies (1:500) at room temperatures for two hours. Next, sections were washed with 1X PBS before staining with Hoechst dye (1:10,000). Imaging was performed using the Zeiss Axio Observer and analyzed using Zeiss Zen Pro, Zeiss Apotome, and ImageJ software.

### Flow cytometry analysis

Mouse bladders were harvested and treated with digestion solution (0.34 U/mL Liberase^TM^ (Roche) and 100 µg/mL DNaseI (ThermoFisher) in 1X DPBS). Bladders were incubated on a heat block shaker for 1 hour at 37°C and 250 rpm. Bladders were vortexed every 15 minutes at a high speed for 30 seconds. After 1 hour, digestion was arrested with a 1X DPBS supplemented with 2% FBS and 0.2 µM EDTA before passing through 40-µm cell strainers. Following a one-hour incubation with anti-mouse CD16/32 monoclonal antibodies, cells were stained with far red viability dye (Invitrogen) and the conjugated antibodies: PE-Cy5 anti-CD45, Brilliant Violet (BV) 650 anti-CD11b, Super Bright (SB) 780 anti-F4/80, eFluor 450 anti-Arginase-1, and PE anti-iNOS monoclonal antibodies. Manufacturers of listed antibodies provided in Supplementary Materials (**Table S12**).

### Gentamicin protection assay

Approximately 50,000 BMDMϕs were seeded per well in a 96-well flat-bottom culture plate. Mϕs were stimulated with the listed concentrations as described in the ***In vitro and ex vivo polarization of Mφs*** section. Furthermore, Mϕs were also seeded with either 1.5 mg/mL Fg, fibrin gel, or both simultaneously. During the 24-hour stimulation at 37°C 5% CO_2_, Mϕs were cultured in DMEM media supplemented with 10% FBS and 1X penicillin/streptomycin. After 24 hours, supernatant of cell culture media was removed, cells were washed three times with 1X DPBS, and replaced with fresh cell culture media free of penicillin/streptomycin. Cells were then infected with either *E. coli* UTI89 or *E. faecalis* OG1RF strains at a multiplicity-of-infection ratio of 1 for a period of 45 minutes to allow uptake of the pathogen in a 37°C 5% CO_2_ incubator. After 45 minutes, supernatant was removed and Mϕs were washed three times with 1X DPBS before being replaced with DMEM media supplemented with 100 µg/mL gentamicin and 1X penicillin/streptomycin. Cells were then incubated in a 37°C 5% CO_2_ incubator for 1-, 2-, and 3-hours post-uptake. After each time point, media was removed, and cells were washed three times with 1X DPBS before treatment with distilled water for 15 minutes. Water was then plated on either LB or BHI media plates and incubated overnight at 37°C for CFU counts.

### Western blot

Approximately 10^6^ RAW 264.7 were seeded per well in a 6-well flat-bottom culture plate. Mϕs were stimulated as described in the ***In vitro and ex vivo polarization of M***ϕ***s*** section, but at concentrations of 12.5, 25, 50, and 100 ng/mL of inducers. Additionally, Mϕs were also seeded with Fg, fibrin gel, or both simultaneously at concentrations of 0.1875, 0.375, 0.75, and 1.5 mg/mL. After 24 hours, supernatant was removed, cells were harvested and centrifuged into a pellet, and resuspended in 5X SDS buffer. The cell lysate was then boiled at 95°C for 5 minutes before 15 µL of the sample was loaded in a SDS-PAGE gel. For probing Arginase-1, samples were run on 14% acrylamide gels for 4 and half hours at 120 V. The gel was then transferred to polyvinylidene difluoride membrane (PVDF, Millipore Sigma) using a semi-dry transfer. After transfer, membranes were blocked in 5% non-fat milk in 1X PBS for one hour at room temperature before incubation with rabbit anti-beta actin (1:7500) and goat anti-Arginase-1 (1:100) primary antibodies diluted in 2% non-fat milk in 1X PBS-Tween (0.1%, PBS-T) overnight at 4°C. Membranes were washed with 1X PBS-T before being probed with IRDye680RD donkey anti-rabbit and IRDye800CW donkey anti-goat secondary antibodies (LI-COR Bioscience) diluted in 2% non-fat milk in 1X PBS-T-SDS (0.01% SDS) for one hour at room temperature. For probing iNOS, samples were run on 7% acrylamide gels for 2 hours at 120 V. Then, the gel was transferred to PVDF membrane using a wet tank transfer method running at 4°C for 2 and half hours. After transfer, membranes were blocked before incubation with rat anti-beta actin (1:7500) and rabbit anti-iNOS (1:100) primary antibodies as previously described. After overnight, membranes were probed with secondary antibodies IRDye680RD donkey anti-rat and IRDye800CW donkey anti-rabbit as previously described. Membranes were then visualized on an Odyssey CxL Reader.

### Statistical analysis

Data derived from these studies were entered into GraphPad Prism 8 software to generate statistical results and graphs. At least more than three independent experiments with replicates were performed for all studies. For data that was parametric, unpaired student two-tailed t-tests was used to determine significance between samples. When data was nonparametric, medians was used to represent the distribution and two-tailed Mann-Whitney U tests was performed. For all tests performed, \**p≤0.05*, \*\**p≤0.005*, \*\*\**p≤0.001*, and \*\*\*\**p≤0.0001*. Statistical test listed in each figure legend.

### List of all materials, reagents, antibodies, and dyes used in this study

List of all materials, reagents, antibodies, and dyes can be found in (**table S12**) table in Supplementary Materials.

## Acknowledgments

We thank the members of both A.L.F.M and F.H.S.T laboratories for their guidance on experimental design, materials, and comments. We also thank the Freimann Life Science Center for mouse breeding and husbandry. Additionally, we thank Dr. Sara Cole from the Notre Dame Integrated Imaging Facility for tissue processing.

## Author contributions

ALFM conceived and supervised the research. Below are each author’s contributions:

Conceptualization: AMM, ALFM

Formal analysis: AMM, JJM, ALFM

Methodology: AMM, JJM, ALFM, DD FHST, VAP, MJF, FJC

Funding acquisition: ALFM, FHST, AMM, VAP, MJF, FJC

Investigation: AMM, ALFM, JJM, MJA, CG, AAL, ERL, EW, TU, KAP, RW, PVS, KNK

Visualization: AMM, ALFM Supervision: ALFM

Writing - original draft: AMM, ALFM

Writing - review & editing: AMM, JJM, MJA, CG, AAL, ERL, EW, TU, KAP, RW, PVS, KNK, VAP, MJF, FJC, FHST, ALFM

## Data and materials availability

The data that support the findings and conclusions in this study are present in this paper and/or supplemental materials. Further inquiries and requests about additional data availability be directed to corresponding author A.L.F.M.

