## Supplemental data for "The catheterized bladder environment induces dysregulation of macrophage polarization exacerbating bacterial UTI"

**Short Title (49 characters):** Catheterized bladder impairs macrophages promoting UTI

**Teaser (121 characters):** Fibrin promotes M2 macrophage polarization resulting in enhanced uropathogen colonization during urinary catheterization.

**Funding:** This work was done in the Flores-Mireles Laboratory and funded by institutional funds from the University of Notre Dame (to A.L.F.M), the National Institute of Health's National Institute of Diabetes and Digestive and Kidney Diseases (NIDDK) grants R01DK128805 (to A.L.F.M. and A. M. M.), the Diversity supplement R01DK12880501-A1S1 (to A. M. M.); R01-HL013423 (to D.D., V.A.P., and F.J.C.); R01-HL160046 and U01-HL143403 (to M.J.F.); R01AI177875 and R21AI71742 (to F.H.S-T); and from Good Venture Foundation (Open Philanthropy) grant (to A.L.F.M., A. M. M, J.J.M., M.J.A., C.G., and E.W.).

**Author contributions:**

ALFM conceived and supervised the research. Below are each author's contributions:

Conceptualization: AMM, ALFM

Formal analysis: AMM, JJM, ALFM

Methodology: AMM, JJM, ALFM, DD FHST, VAP, MJF, FJC

Funding acquisition: ALFM, FHST, AMM, VAP, MJF, FJC

Investigation: AMM, ALFM, JJM, MJA, CG, AAL, ERL, EW, TU, KAP, RW, PVS, KNK

Visualization: AMM, ALFM

Supervision: ALFM

Writing - original draft: AMM, ALFM

Writing - review & editing: AMM, JJM, MJA, CG, AAL, ERL, EW, TU, KAP, RW, PVS, KNK, VAP, MJF, FJC, FHST, ALFM

**Competing interests:** The authors declare no competing financial interests.

**Data and materials availability:** The data that support the findings and conclusions in this study are present in this paper and/or supplemental materials. Further inquiries and requests about additional data availability be directed to corresponding author A.L.F.M.

#### FIGURES

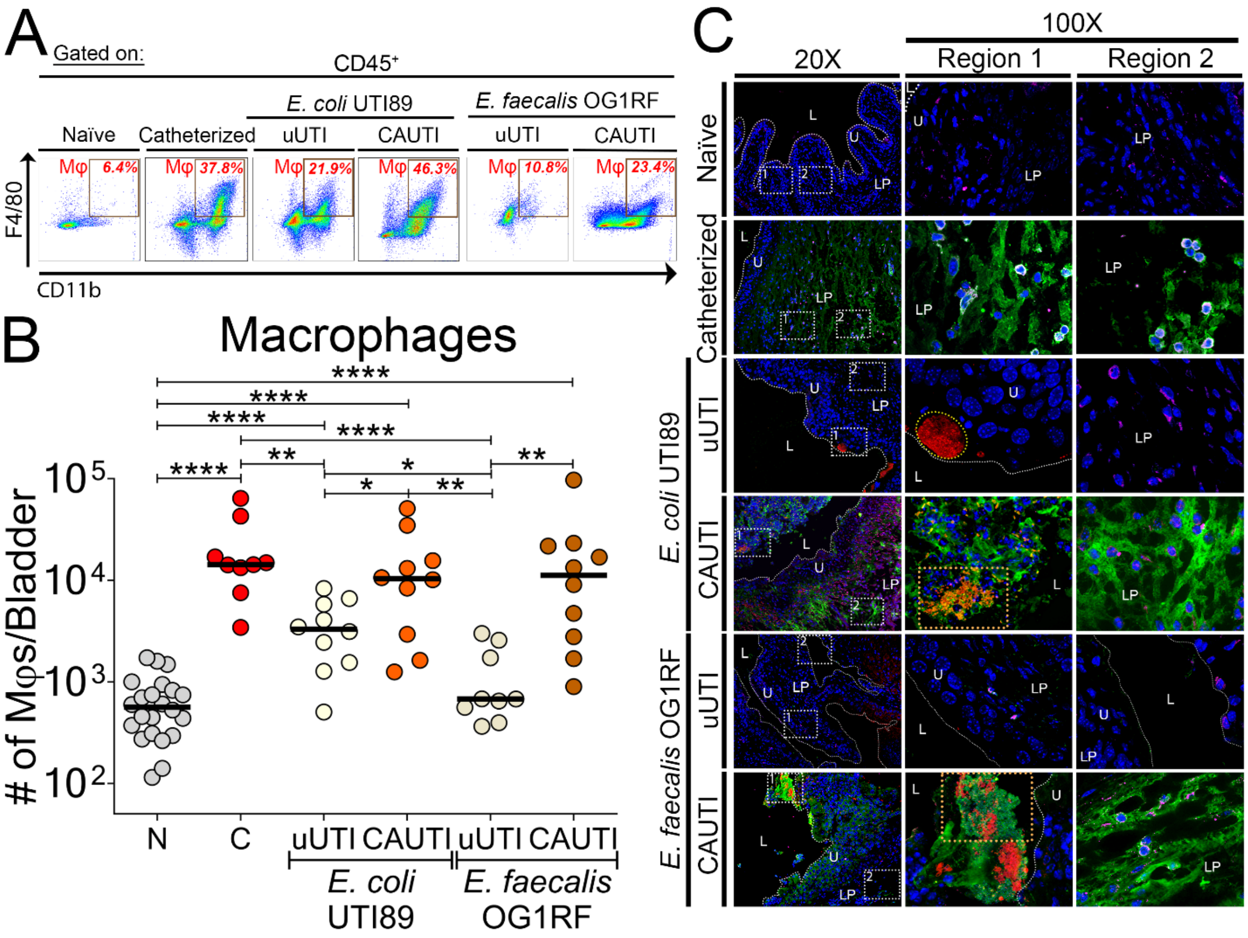


**Figure 1. Urinary catheterization induces robust Fg/fibrin accumulation and Mφ response.** **(A)** Representative dot plots of Mφs (Live single CD45$^+$CD11b$^+$F4/80$^+$; **fig. S1, gating strategy**). **(B)** Quantification of number of Mφs in the bladder. Each dot represents one mouse; horizontal lines are medians. The Mann-Whitney U test was used for determining statistical significance; *, $P \leq 0.05$; **, $P \leq 0.005$; ***, $P \leq 0.0005$; ****, $P \leq 0.0001$. **(C)** C57BL/6 female mice bladders were either naïve (N), catheterized without infection (C), or infected in absence (uUTI) or presence (CAUTI) of a catheter with either *E. coli* UTI89 or *E. faecalis* OG1RF for 24 hours. At 24 hours, bladders were stained with DAPI for cell nuclei (blue) and antibodies to detect Fg/fibrin (green), *E. coli* or *E. faecalis* (red), and Mφs (anti-F4/80; magenta). **fig. S2-S7, individual channels**. For

all representative images, white boxes at 20x represent zoomed-in areas of 100x magnification and the white broken line separates the lumen (L) from the urothelium surface (U) and the lamina propria (LP). Orange dotted rectangle depicts uropathogen-Fg biofilms.

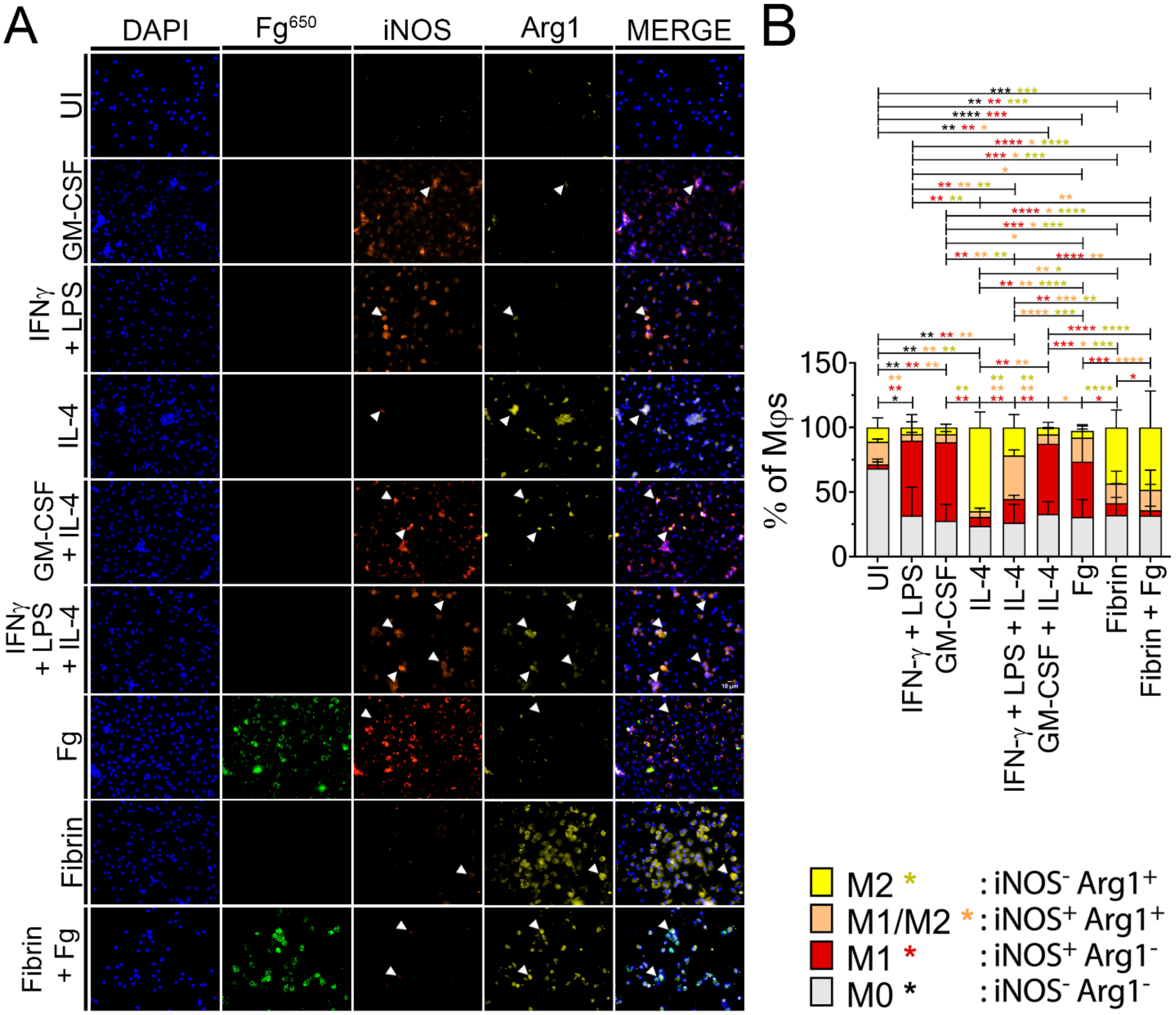


**Figure 2. Fibrin suppresses Fg-induced M1-Mφ activation. (A)** BMD-Mφs were either uninduced in DMEM cell culture media (UI) or induced with either 100 ng/ml of either GM-CSF or IL-4, both GM-CSF+IL-4, 1.5 mg/ml of either Alexa Fluor 650-conjugated Fg (Fg$^{650}$; green) or fibrin or both (fibrin + Fg) simultaneously for 24 hours. Cells were stained with DAPI for cell nuclei and antibodies to detect iNOS (orange) and Arginase-1 (yellow) for IF analysis (representative images). Magnification is at 40x. (**B**) Percent of Mφs that are either M0 (gray; iNOS$^-$Arg1$^-$), M1 (red; iNOS$^+$Arg1$^-$), hybrid M1/M2 (orange; iNOS$^+$Arg1$^+$), or M2 (yellow; iNOS$^-$ Arg1$^+$). Percent of Mφs per phenotype was calculated by the number of Mφs per phenotype

divided by total number of Mφs in each field (**table S2**, n=6-12 fields). Values represent mean ± SD. The Mann-Whitney U test was used where $P < 0.05$ was considered statistically significant; *, $P \leq 0.05$; **, $P \leq 0.005$; ***, $P \leq 0.0005$; ****, $P \leq 0.0001$.

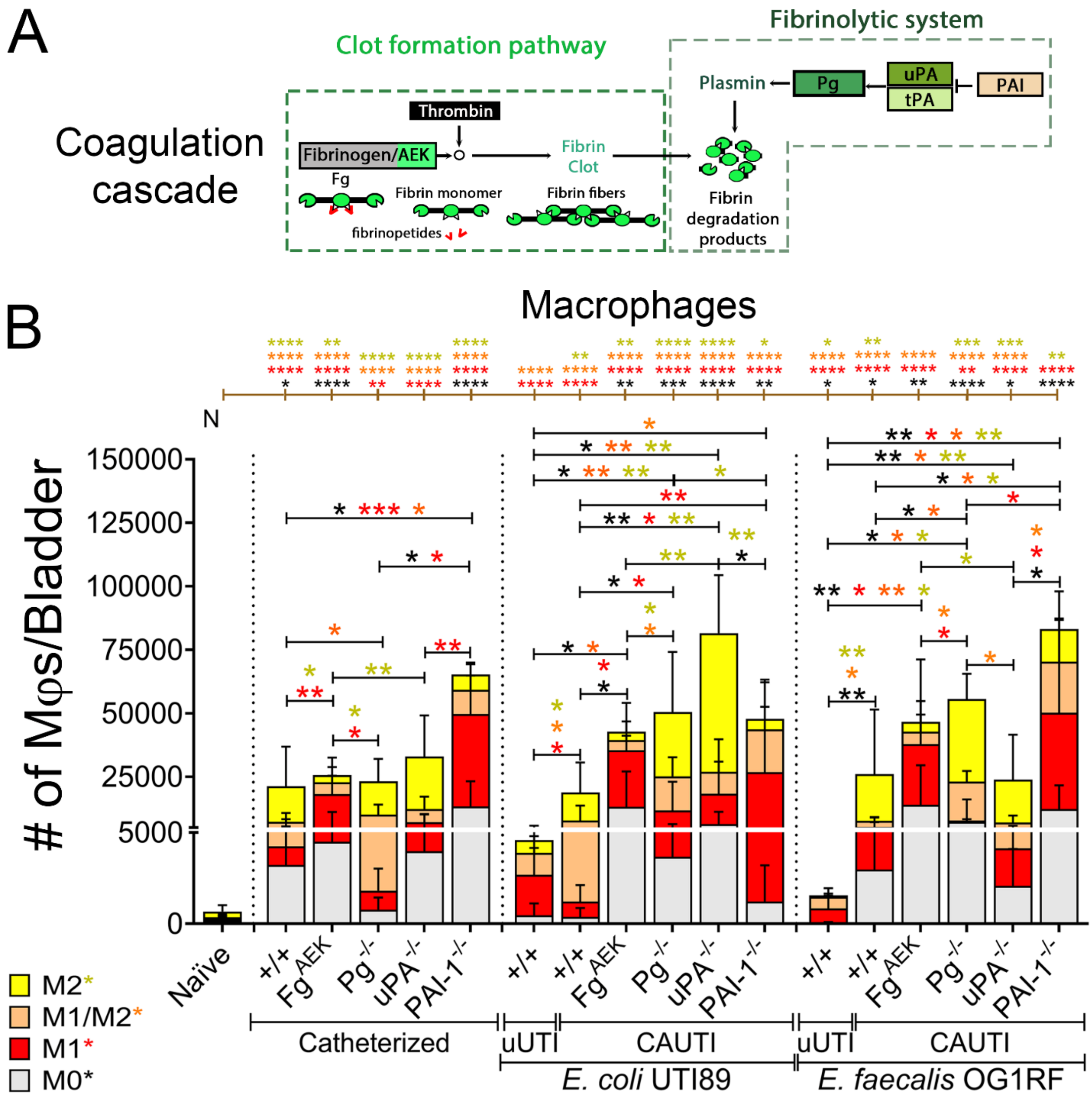


**Figure 3. Fibrinogen and fibrin differentially polarize Mφs during catheterization. (A)** Coagulation cascade diagram. C57BL/6 female WT ($^{+/+}$) and coagulation-transgenic mice bladders (**table S7**) were uninfected or infected with ~$10^7$ CFUs of *E. coli* UTI89 or *E. faecalis* OG1RF in absence (uUTI) or presence of a catheter (CAUTI) for 24 hours. Coagulation-transgenic mice used are: Fg$^{AEK}$ (soluble Fg, no fibrin formation), Pg$^{-/-}$ and uPA$^{-/-}$ (elevated fibrin accumulation, no

plasmin activation), and PAI-$1^{-/-}$ (continuous fibrin clot degradation). Naïve (N) mice were neither catheterized nor infected. Bladders were then harvested and digested to isolate single cells before incubation with CD16/32 antibodies for blocking and viability dye. For flow cytometry analysis, cells were then stained with conjugated anti-mouse primary antibodies for CD45, CD11b, F4/80, iNOS, and Arginase-1. (**B**) Mφs ($CD45^+CD11b^+F4/80^+$) were gated from live single cell populations before further gating for M0 ($Arg1^-iNOS^-$), M1 ($Arg1^-iNOS^+$), M1/M2 ($Arg1^+iNOS^+$), and M2 ($Arg1^+iNOS^-$) populations (**fig. S1, gating strategy**). The values represent mean ± SD. Differences between groups were tested for significance using the Mann-Whitney U test. Statistical significance between each group and naïve shown as brown brackets on top of each graph. *, $P \leq 0.05$; **, $P \leq 0.005$; ***, $P \leq 0.0005$; ****, $P \leq 0.0001$. Individual data points in **fig. S10**.

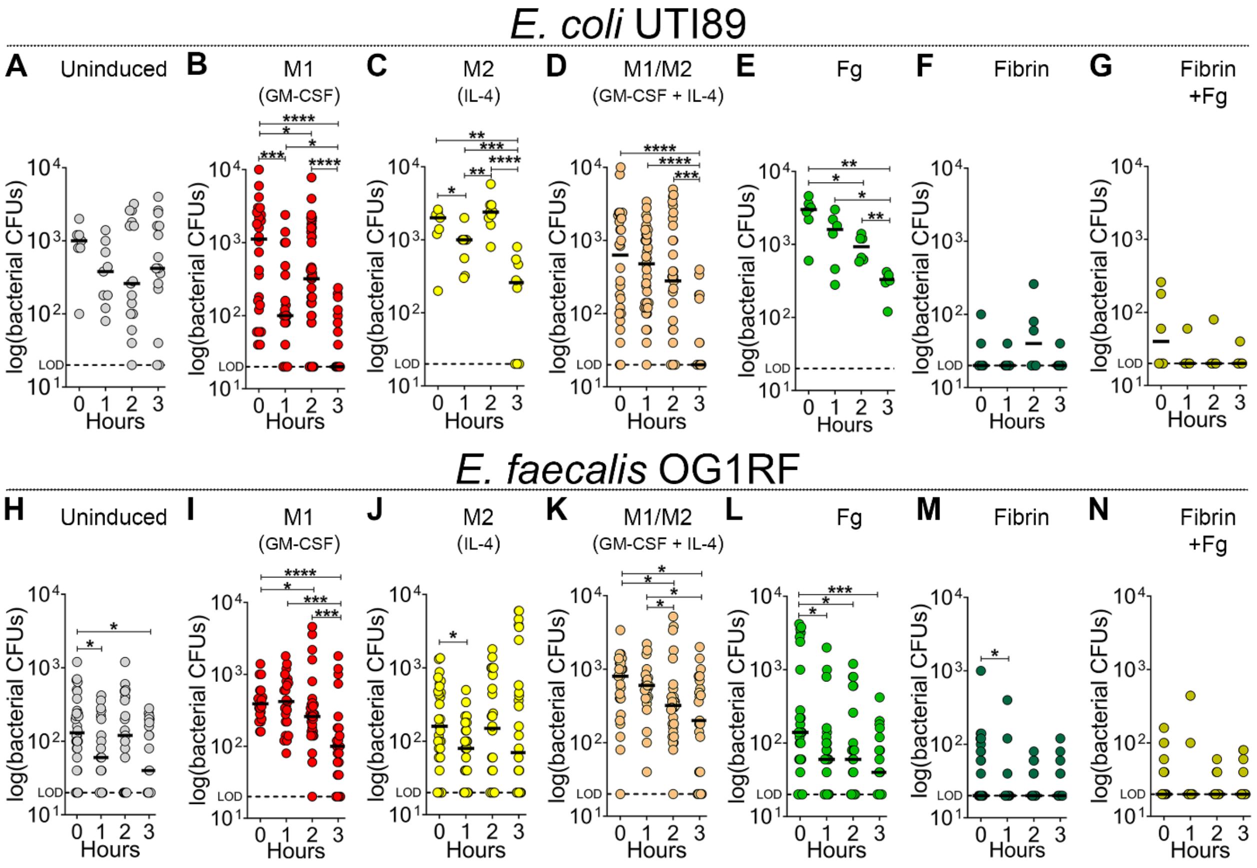


**Figure 4. Fibrin impairs the antimicrobial response of Mφs against pathogens.** BMDMφs were either uninduced (M0, **A & H**) or stimulated with 100 ng/mL of GM-CSF (M1, **B & I**), IL-4 (M2, **C & J**), or cotreated with GM-CSF and IL-4 (M1/M2, **D & K**), 1.5 mg/mL of Fg (**E & L**) or fibrin (**F & M**) or both simultaneously (**G & N**) for 24 hours. Mφs were then infected with either (**A-G**) opsonized *E. coli* UTI89 or (**H-N**) *E. faecalis* OG1RF at a multiplicity-of-infection ratio of 1 (50,000 bacteria to 50,000 Mφs) for initial uptake (45 minutes). Supernatant was removed after initial uptake and Mφs were then treated with gentamicin for 1-3 hours to remove extracellular bacteria. Then, Mφs were bursted with pure sterile distilled water for intracellular bacteria retrieval and CFU counts. Horizontal line represents median value. Each data point is one replicate. The horizontal broken line represents the limit of detection (LOD) of viable bacteria. The Mann-Whitney U test was used to determine significant difference between time points; *, *P*

$\leq 0.05$; **, $P \leq 0.005$; ***, $P \leq 0.0005$; ****, $P \leq 0.0001$.

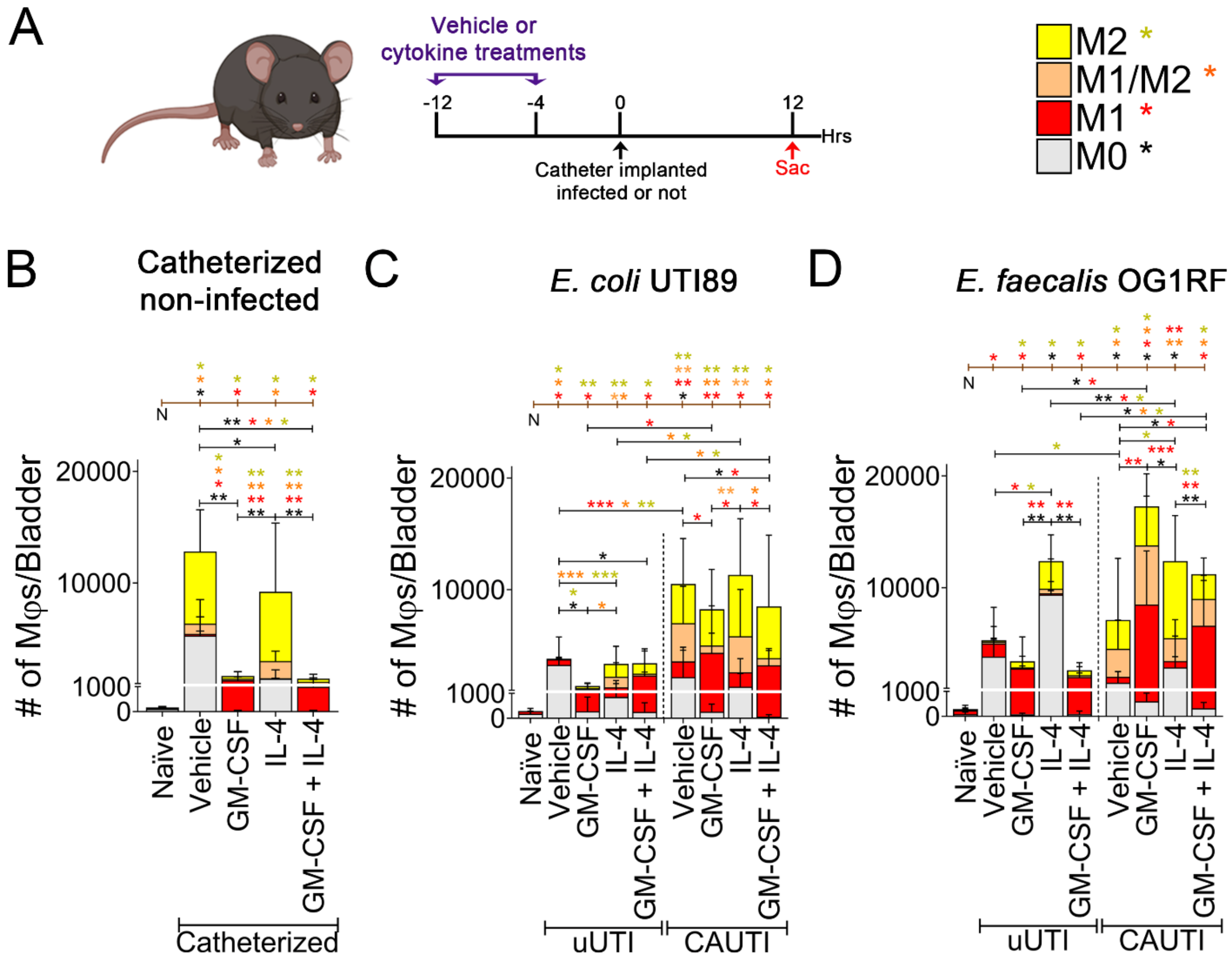


**Figure 5. GM-CSF outcompetes IL-4 and fibrin in promoting M1 polarization in CAUTIs.** **(A)** C57BL/6 WT mice were intraperitoneally (i.p.) injected with two doses of either sterile 1X PBS (vehicle), 200 ng/mouse of GM-CSF, IL-4, or co-treatment with GM-CSF and IL-4 at 12 and 4 hours prior to catheterization and/or infection. Twelve hours post-catheterization and/or infection, bladders were harvested and digested to isolate single cells for flow cytometry analysis. Cells were then stained with a viability dye and for conjugated antibodies for CD45, CD11b, F4/80, iNOS, and Arginase-1. Mφs was gated from live single cell populations before further gating for M0-Mφs (iNOS$^-$ Arg1$^-$), M1-Mφs (iNOS$^+$Arg1$^-$), M1/M2-Mφs (iNOS$^+$Arg1$^+$), and M2-Mφs (iNOS$^-$Arg1$^+$) populations (**fig. S1, gating strategy**). Naïve (N) mice were neither catheterized nor infected. Mice were either (**B**) catheterized without infection, (**C**) infected with $10^8$ CFUs of *E.*

*coli* UTI89 or (**D**) infected with $10^8$ CFUs of *E. faecalis* OG1RF in absence (uUTI) or presence of a catheter (CAUTI). Values represent mean ± SD. Differences between groups were tested for significance using the Mann-Whitney U test. Statistical significance between each group and naïve (N) shown as brown brackets on top of each graph. *, $P \leq 0.05$; **, $P \leq 0.005$; ***, $P \leq 0.0005$; ****, $P \leq 0.0001$. Individual data points in **fig. S13**.

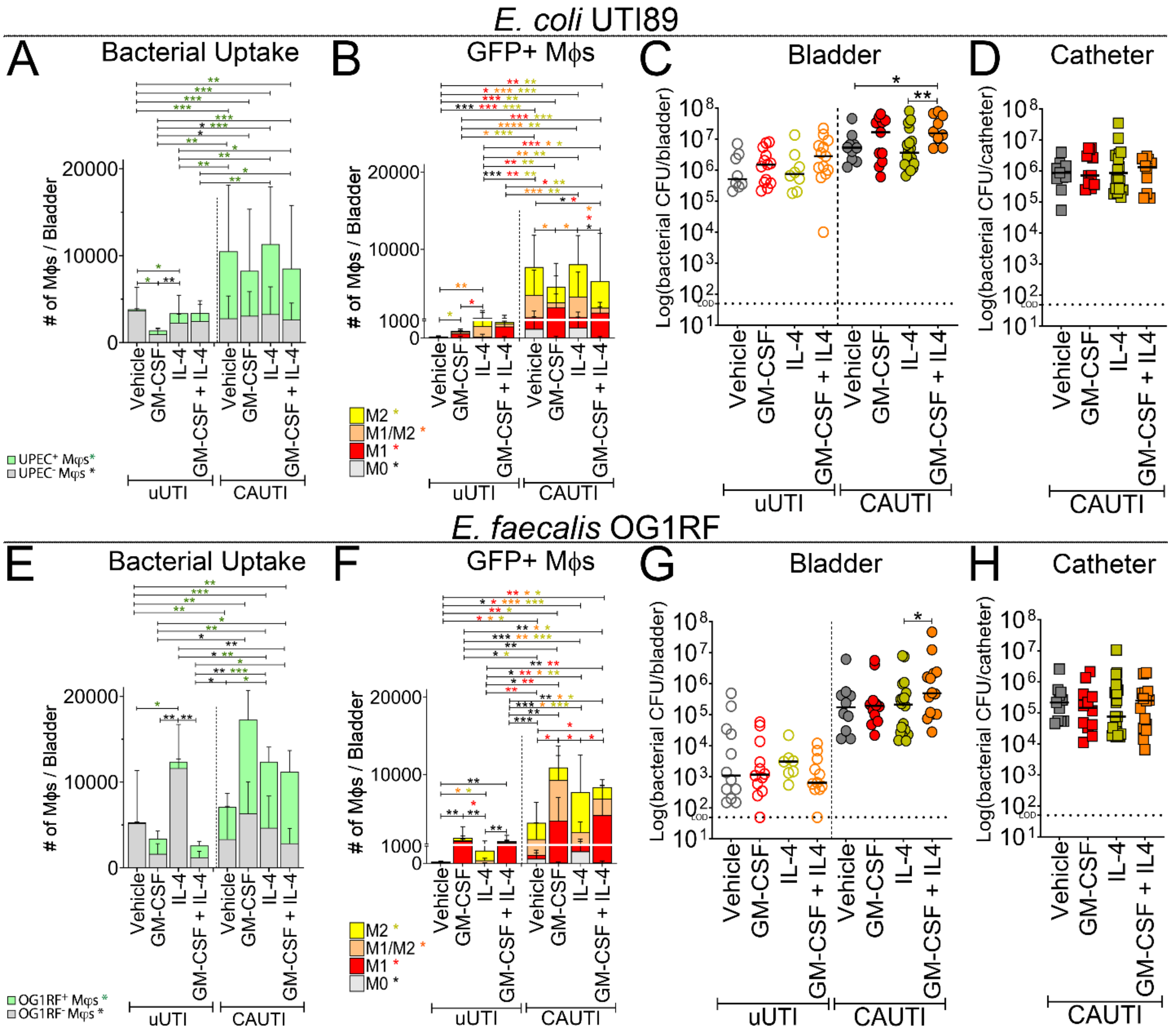


**Figure 6. Fibrin outcompetes cytokines in suppressing Mφ antimicrobial response during catheterization.** WT-C57BL/6 mice were i.p. injected with two doses of either sterile 1X PBS (vehicle) or cytokines at 12 and 4 hours prior to catheterization and/or infection as described in **Fig. 5A**. Naïve (N) mice were neither catheterized nor infected. Mice were either (**A-D**) infected with $10^8$ CFUs of *E. coli* UTI89-GFP strain or (**E-G**) infected with $10^8$ CFUs of *E. faecalis* OG1RF-GFP strain in absence (uUTI) or presence of a catheter (CAUTI). Twelve hours post-catheterization and/or infection, bladders were harvested and digested to isolate single cells for flow cytometry analysis and staining as described in **Fig. 5A**. (**A, E**) GFP$^+$Mφs (phagocytized the

pathogen, UPEC$^+$ or OG1RF$^+$) and GFP$^-$Mφs (did not phagocytize the pathogen, UPEC$^-$ or OG1RF$^-$) were each gated (**fig. S15, gating strategy**). (**B, F)** Evaluation of polarization state of GFP$^+$ Mφs populations phagocytizing the pathogen**. (C-D, G-H**) Twelve hours post-catheterization and/or infection, the bacterial burden of each strain was determined by CFU count of (**C, G**) bladders and (**D, H**) catheters. Horizontal line represents median value. Each data point is one mouse. The Mann-Whitney U test was used to test for significance between groups. *, $P \leq 0.05$; **, $P \leq 0.005$; ***, $P \leq 0.0005$; ****, $P \leq 0.0001$. The horizontal broken line represents the limit of detection (LOD) of viable bacteria. Three to four independent experiments were performed with 6-16 mice for each experimental group.

Science Advances
AAAS

### Supplementary Materials for

#### Catheterized bladder environment induces anti-inflammatory macrophage polarization resulting in enhanced bacterial UTI

Armando Magallanes Marrufo *et al.*

**This PDF file includes:**

Supplementary Text
Figs. S1 to S16
Tables S1 to S12

**SUPPLEMENTARY MATERIALS:**

**Supplementary Figures:**

**Supplementary Figure 1.** Macrophage gating strategy.

**Supplementary Figure 2.** Low presence of bladder Mφs in naïve bladders.

**Supplementary Figure 3.** Mφ recruitment upon urinary catheterization.

**Supplementary Figure 4.** Mφ response to *E. coli* UTI89 bladder colonization in absence of a catheter.

**Supplementary Figure 5.** Mφ recruitment upon *E. coli* UTI89 bladder colonization during CAUTI.

**Supplementary Figure 6.** Mφ recruitment during *E. faecalis* OG1RF bladder colonization in absence of a catheter.

**Supplementary Figure 7.** Mφ recruitment in response to *E. faecalis* OG1RF bladder colonization during CAUTI.

**Supplementary Figure 8.** Fibrinogen and fibrin promote iNOS and Arginase-1 in RAW 264.7.

**Supplementary Figure 9.** Fibrin suppresses Fg-induced M1 Mφ activation in RAW 264.7.

**Supplementary Figure 10.** Total number of Mφs and polarization states in bladders of WT and coagulation-transgenic mice that were catheterized and/or infected with either *E. coli* UTI89 or *E. faecalis* OG1RF.

**Supplementary Figure 11.** Fibrin suppressed Mφ phagocytosis of bacteria.

**Supplementary Figure 12**. Catheterization promotes bladder inflammation regardless of inducer.

**Supplementary Figure 13**. Total number of Mφs and polarization states in the bladder treated with different inducers before catheterization and/or infection with either *E. coli* UTI89 or *E. faecalis* OG1RF.

**Supplemental Figure 14.** Comparison of Mφ polarization between vehicle-treated mice during uUTI and CAUTI.

**Supplementary Figure 15.** Representative gating strategy to determine Mφ phagocytic activity.

**Supplementary Figure 16**. Bacterial colonization of the kidneys, spleen, and heart during uUTI and CAUTI.

**Supplementary Tables:**

**Supplementary Table 1.** Number of Mφs in bladders of WT mice that were catheterized and/or infected with either *E. coli* UTI89 or *E. faecalis* OG1RF.

**Supplementary Table 2.** Percentage of BMDMφs in each condition that are M0 ($iNOS^-$ $Arg1^-$), M1 ($iNOS^+$ $Arg1^-$), M1/M2 ($iNOS^+$ $Arg1^+$), and M2 ($iNOS^-$ $Arg1^+$).

**Supplementary Table 3.** Percentage of RAW 264.7 Mφs in each condition that are M0 ($iNOS^-$ $Arg1^-$), M1 ($iNOS^+$ $Arg1^-$), M1/M2 ($iNOS^+$ $Arg1^+$), and M2 ($iNOS^-$ $Arg1^+$).

**Supplementary Table 4.** Total number of Mφs in bladders of WT and coagulation-transgenic mice that were catheterized and/or infected with either *E. coli* UTI89 or *E. faecalis* OG1RF.

**Supplementary Table 5.** Number and percentage of Mφs that are M0, M1, M1/M2, and M2 in bladders of WT and coagulation-transgenic mice that were catheterized and/or infected with either *E. coli* UTI89 or *E. faecalis* OG1RF.

**Supplementary Table 6.** Table of mouse strains used in this study.

**Supplementary Table 7.** Total number of Mφs in bladders of WT mice treated with inducers before catheterization and/or infection with either *E. coli* UTI89 or *E. faecalis* OG1RF.

**Supplementary Table 8.** Number and percentage of Mφs that are M0, M1, M1/M2, or M2 in bladders of WT mice treated with inducers before catheterization and/or infection with either *E.*

*coli* UTI89 or *E. faecalis* OG1RF.

**Supplementary Table 9.** Number and percentage of Mφs that phagocytized pathogens in bladders of WT mice treated with inducers before catheterization and infection with either *E. coli* UTI89 or *E. faecalis* OG1RF.

**Supplementary Table 10.** Number and percentage of GFP+ Mφs that are M0, M1, M1/M2, or M2 in bladders of WT mice treated with inducers before catheterization and infection with either *E. coli* UTI89 or *E. faecalis* OG1RF.

**Supplementary Table 11.** Number and percentage of GFP- Mφs that are M0, M1, M1/M2, or M2 in bladders of WT mice treated with inducers before catheterization and infection with either *E. coli* UTI89 or *E. faecalis* OG1RF.

**Supplementary Table 12.** Table of materials, reagents, antibodies, and dyes used in this study.

**Supplementary Figures**

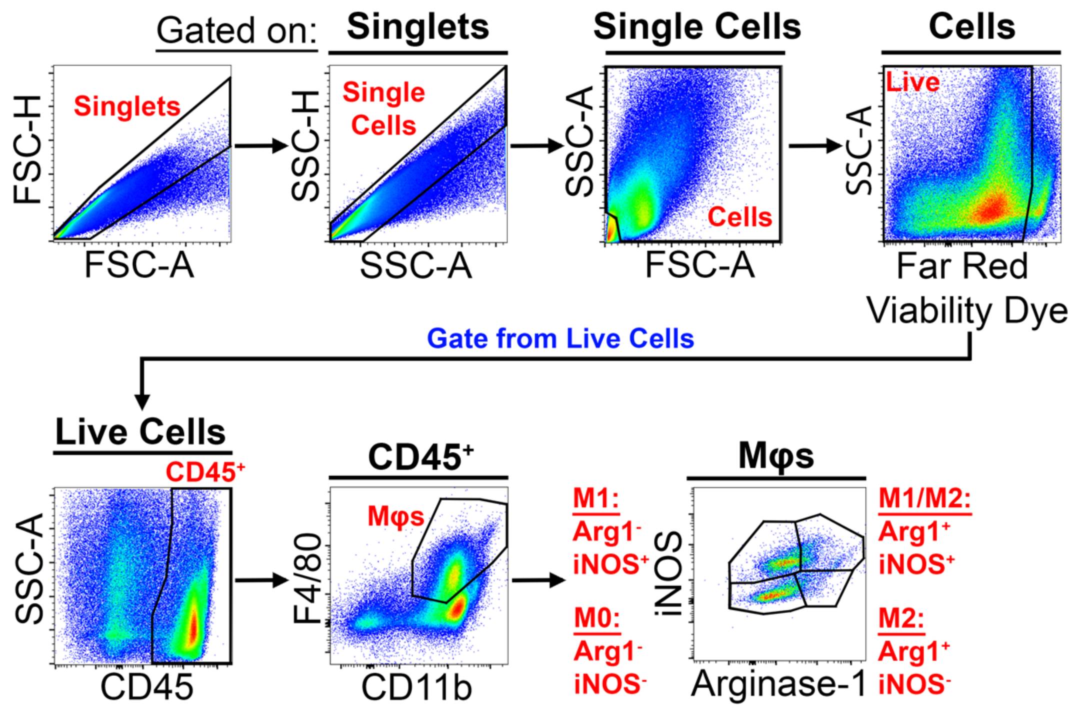


**Supplementary Figure 1. Macrophage gating strategy.** Gating strategy to identify M0 (Arg1$^-$iNOS$^-$), M1 (Arg1$^-$iNOS$^+$), M1/M2 (Arg1$^+$iNOS$^+$), and M2 (Arg1$^+$iNOS$^-$) Mφs populations. Bladders were harvested and digested in buffer containing Liberase$^{TM}$ for single cell isolation. Fc receptors in single-cell suspensions were blocked with anti-mouse CD16/CD32 fragments and stained with Far-Red viability dye and antibodies for CD45, CD11b, F4/80, Arginase-1, and iNOS. Mφs (CD45$^+$CD11b$^+$F4/80$^+$) were gated from live single CD45$^+$ cells.

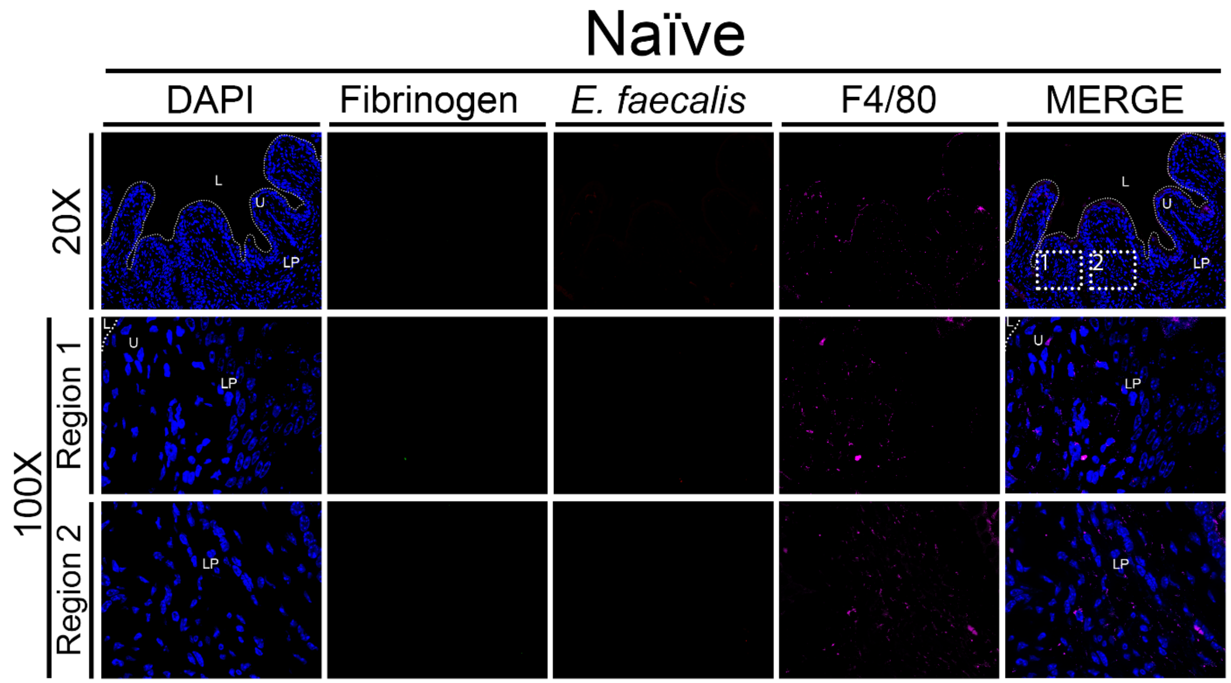


**Supplementary Figure 2. Low presence of bladder Mφs in naïve bladders.** C57BL/6 female mice bladders were neither implanted nor infected. Bladder tissues were harvested, formalin-fixed, and paraffin-embedded. Bladders were subjected to IF analysis using antibodies to detect for DAPI for cell nuclei (blue), Fg (green), *E. faecalis* (red), and Mφs (anti-F4/80; magenta). White boxes represent zoomed-in sections of higher magnification at 100x.

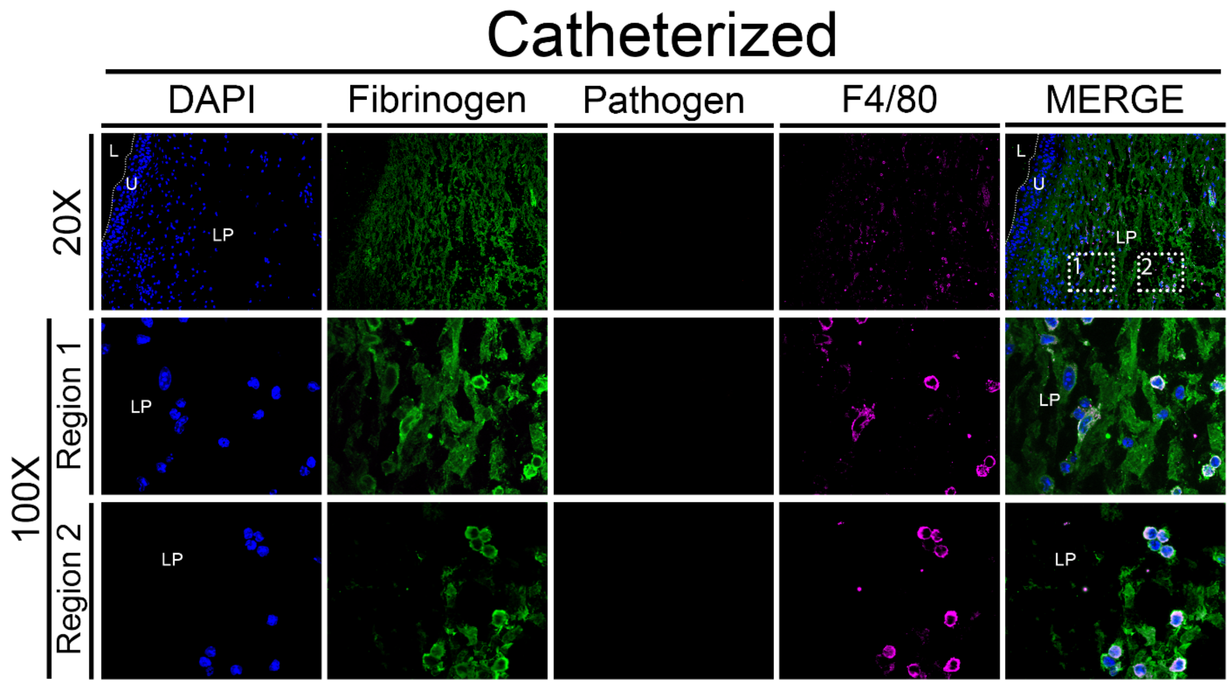


**Supplementary Figure 3. Mφ recruitment upon urinary catheterization.** Mice bladders were implanted without infection. At 24 hours post-catheterization (hpc), bladder tissues were harvested, formalin-fixed, and paraffin-embedded. Bladders were subjected to IF analysis using antibodies to detect for DAPI for cell nuclei (blue), Fg (green), *E. faecalis* (red), and Mφs (anti-F4/80; magenta). White boxes represent zoomed-in sections of higher magnification at 100x.

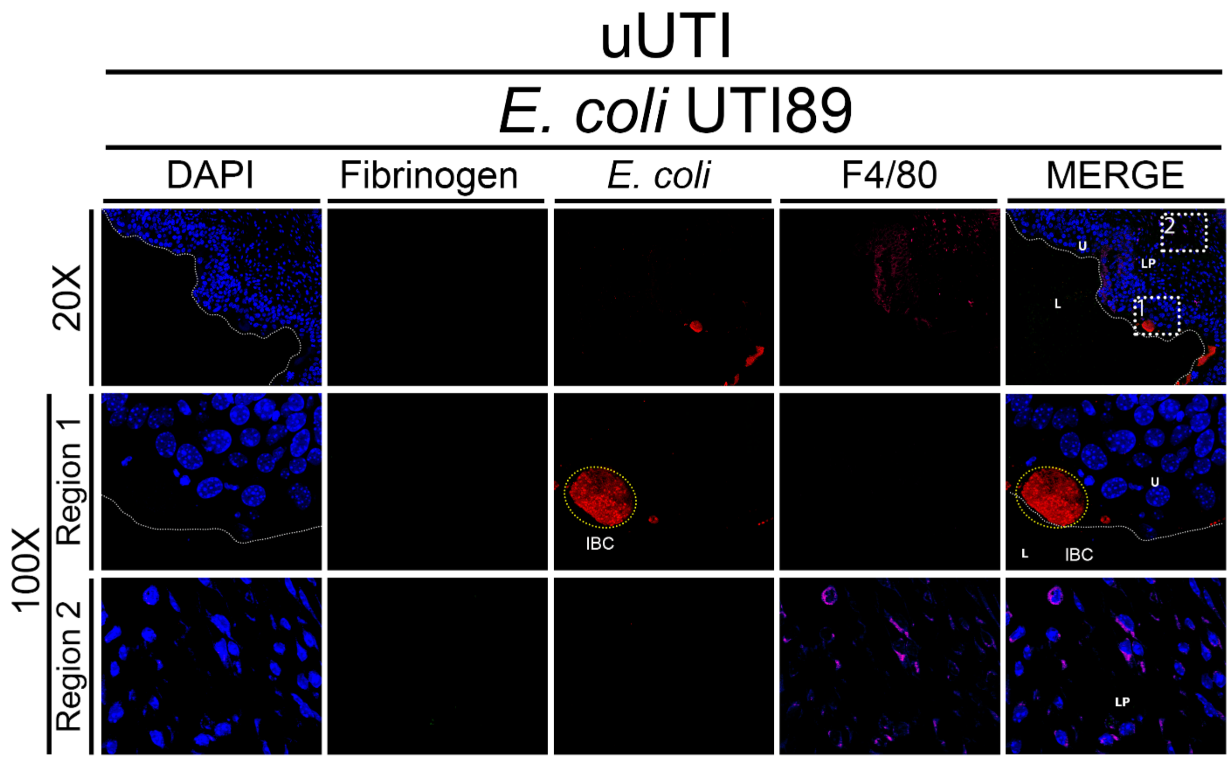


**Supplementary Figure 4. Mφ response to *E. coli* UTI89 bladder colonization in absence of a catheter.** Non-implanted mice bladders were infected with $10^7$ CFUs of *E. coli* strain UTI89. At 24 hpi, bladder tissues were harvested, formalin-fixed, and paraffin-embedded. Bladders were subjected to IF analysis using antibodies to detect for DAPI for cell nuclei (blue), Fg (green), *E. coli* (red), and Mφs (anti-F4/80; magenta). White boxes represent zoomed-in sections of higher magnification at 100x. Yellow dotted ellipse depicts *E. coli* intracellular bacterial communities (IBCs).

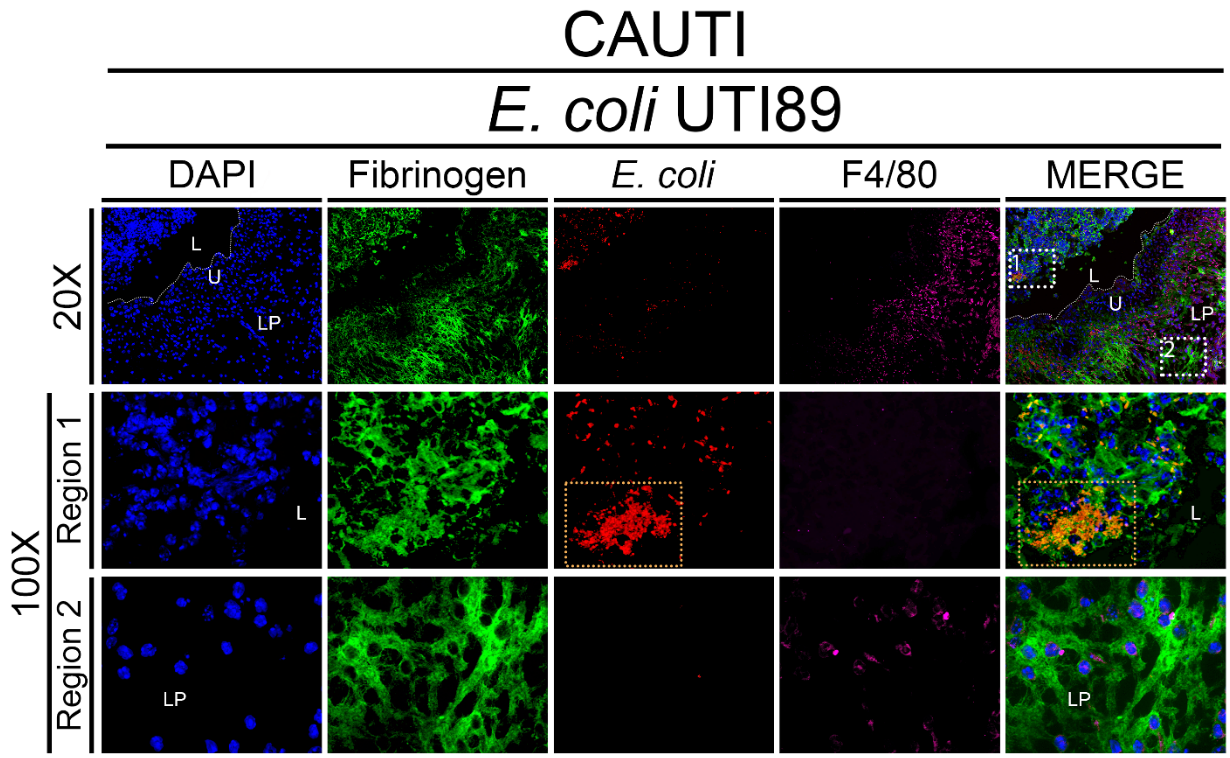


**Supplementary Figure 5. Mφ recruitment upon *E. coli* UTI89 bladder colonization during CAUTI.** Mice bladders were implanted and infected with $10^7$ CFUs of *E. coli* strain UTI89. At 24 hpic, bladder tissues were harvested, formalin-fixed, and paraffin-embedded. Bladders were subjected to IF analysis using antibodies to detect for DAPI for cell nuclei (blue), Fg (green), *E. coli* (red), and Mφs (anti-F4/80; magenta). White boxes represent zoomed-in sections of higher magnification at 100x. Orange dotted rectangle depicts *E. coli*-Fg biofilms.

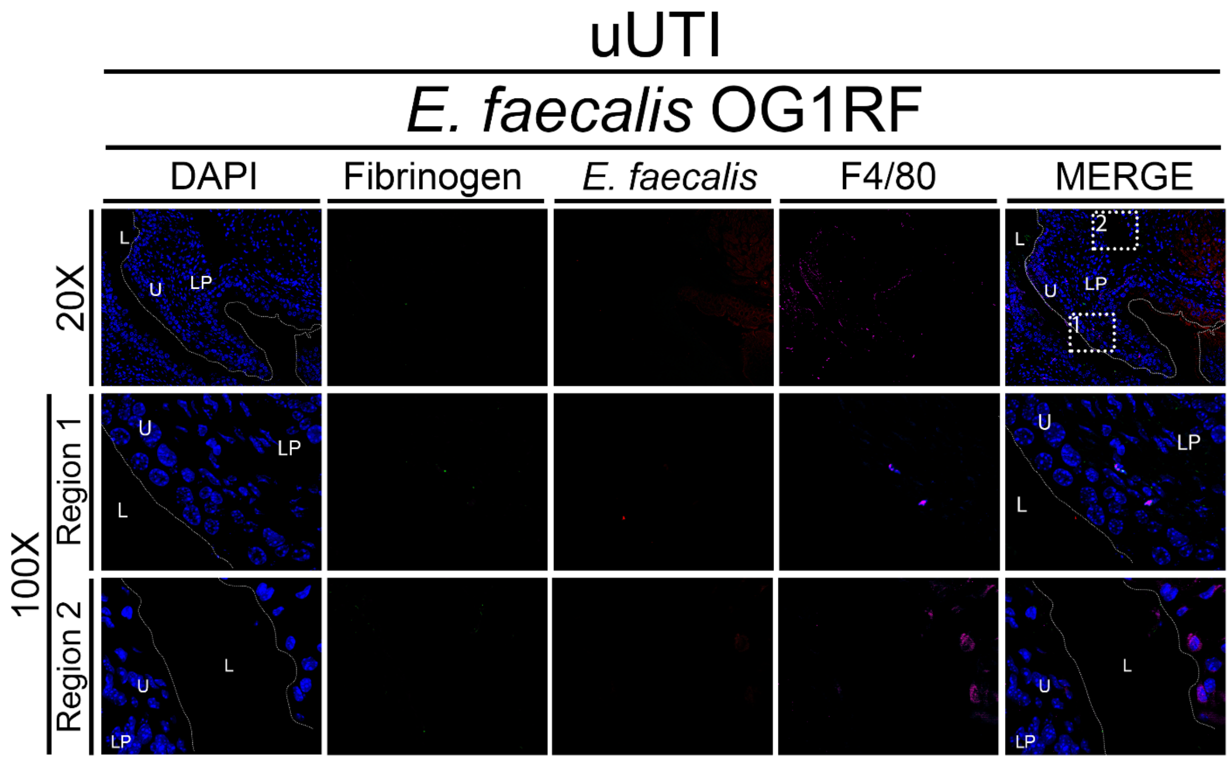


**Supplementary Figure 6. Mφ recruitment during *E. faecalis* OG1RF bladder colonization in absence of a catheter.** Non-implanted mice bladders were infected with $10^7$ CFUs of *E. faecalis* strain OG1RF. At 24 hpi, bladder tissues were harvested, formalin-fixed, and paraffin-embedded. Bladders were subjected to IF analysis using antibodies to detect for DAPI for cell nuclei (blue), Fg (green), *E. faecalis* (red), and Mφs (anti-F4/80; magenta). White boxes represent zoomed-in sections of higher magnification at 20x and 100x.

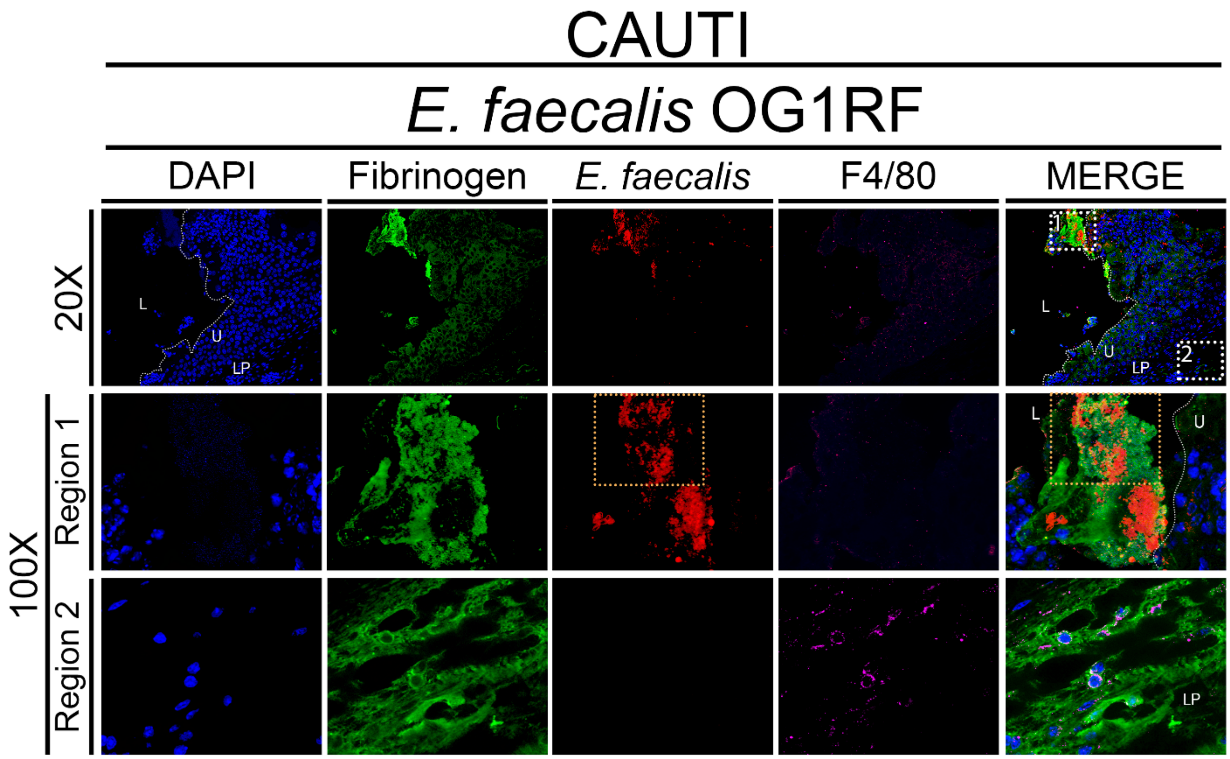


**Supplementary Figure 7. Mφ recruitment in response to *E. faecalis* OG1RF bladder colonization during CAUTI.** Mice bladders were implanted and infected with $10^7$ CFUs of *E. faecalis* strain OG1RF. At 24 hpic, bladder tissues were harvested, formalin-fixed, and paraffin-embedded. Bladders were subjected to IF analysis using antibodies to detect for DAPI for cell nuclei (blue), Fg (green), *E. faecalis* (red), and Mφs (anti-F4/80; magenta). White boxes represent zoomed-in sections of higher magnification at 20x and 100x. Orange dotted rectangle depicts *E. faecalis*-Fg biofilms.

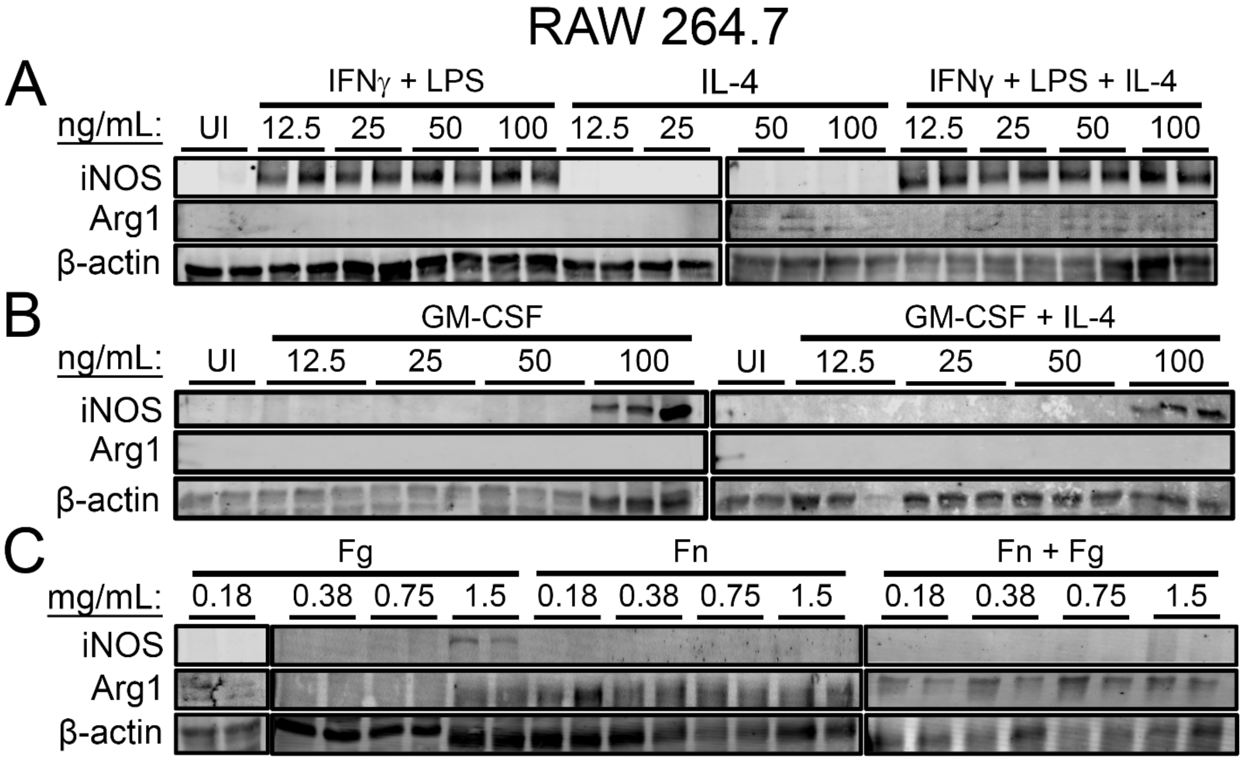


**Supplementary Figure 8. Fibrinogen and fibrin promote iNOS and Arginase-1 in RAW 264.7. (A)** One million RAW 264.7 Mφs was either uninduced in DMEM cell culture media (UI) or induced with either IFNγ + LPS, GM-CSF, IL-4, or mixed signals (IFNγ + LPS +IL-4 or GM-CSF + IL-4) at 12.5, 25, 50, and 100 ng/ml for 24 hours. (**B**) Mφs were stimulated with either Fg, fibrin (Fn) or both (Fn + Fg) simultaneously at 0.18, 0.375, 0.75, and 1.5 mg/ml for 24 hours. iNOS and Arginase-1 protein abundance was assessed by western blot analysis with β-actin as the loading control.

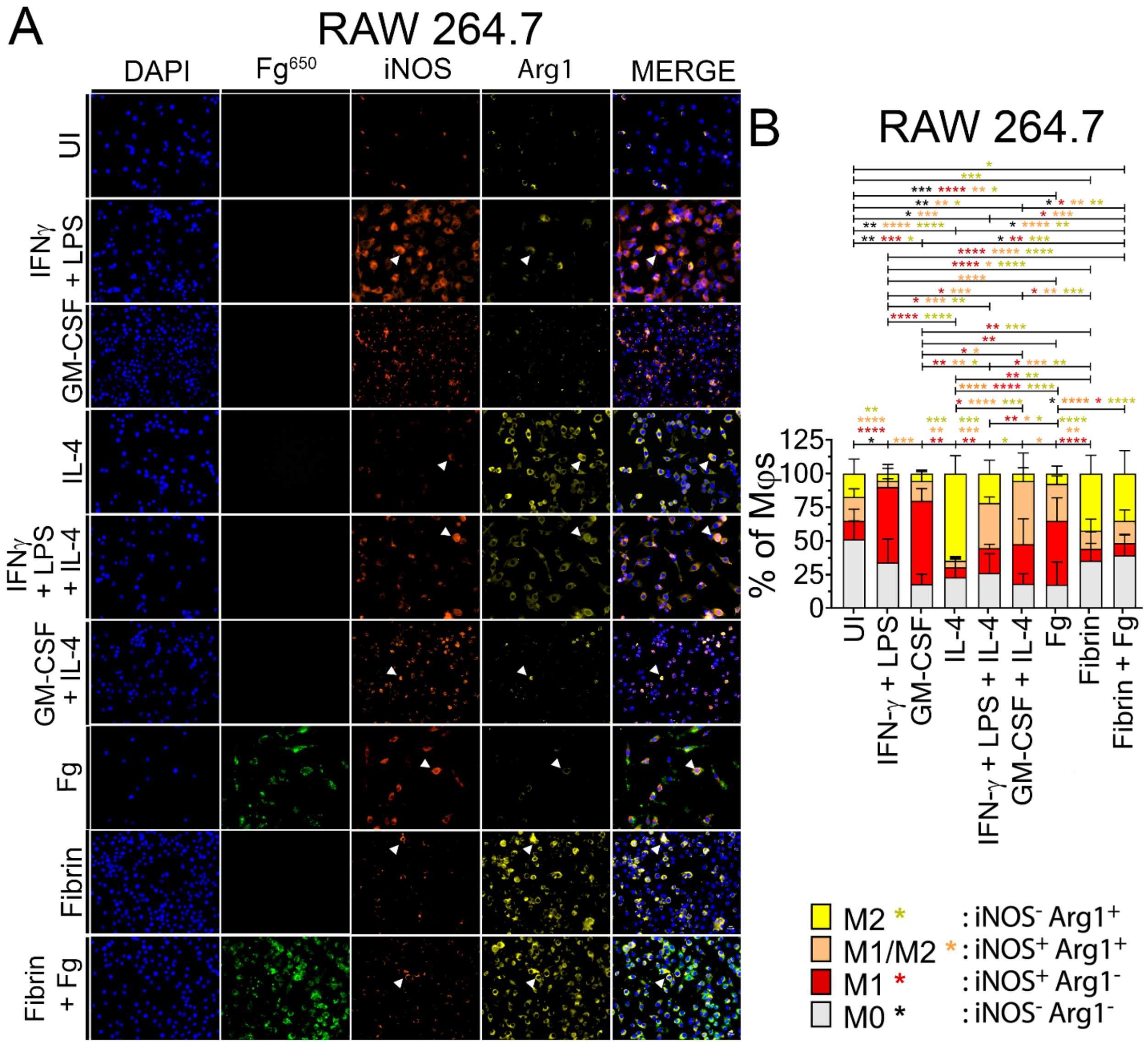


**Supplementary Figure 9. Fibrin suppresses Fg-induced M1 Mφ activation in RAW 264.7.** **(A)** RAW 264.7 Mφs was either uninduced in DMEM cell culture media (UI) or induced with either 100 ng/ml of IFNγ + LPS, GM-CSF, IL-4, all three (IFNγ + LPS + IL-4), or both GM-CSF+IL-4. Mφs were also stimulated with 1.5 mg/ml of either Alexa Fluor 650-conjugated Fg ($Fg^{650}$; green) or fibrin or both (Fibrin + Fg) simultaneously for 24 hours. Cells were stained with DAPI for cell nuclei and antibodies to detect iNOS (orange) and Arginase-1 (yellow) for IF analysis (representative images). Magnification is at 40x. (**B**) Percent of Mφs that are either M0 (gray; $iNOS^-Arg1^-$), M1 (red; $iNOS^+Arg1^-$), hybrid M1/M2 (orange; $iNOS^+Arg1^+$), and M2 (yellow; $iNOS^-Arg1^+$). Percent of Mφs per phenotype was calculated by the number of Mφs per phenotype divided by total number of Mφs in each field (n = 6-12 fields). Values represent mean ± SD. The Mann-Whitney U test was used where $P < 0.05$ was considered statistically significant; *, $P \leq 0.05$; **, $P \leq 0.005$; ***, $P \leq 0.0005$; ****, $P \leq 0.0001$.

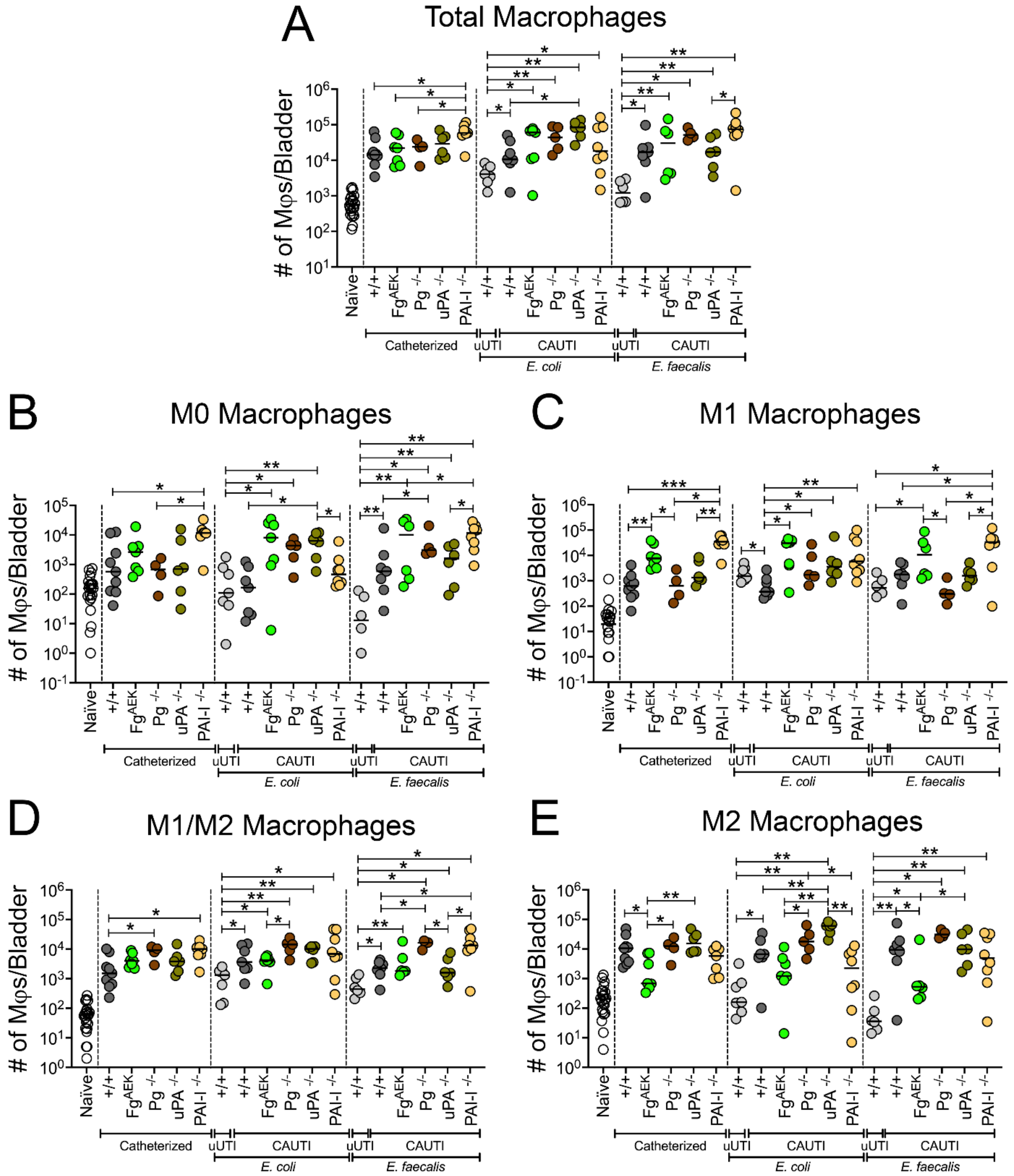


**Supplementary Figure 10. Total number of Mφs and polarization states in bladders of WT and coagulation-transgenic mice that were catheterized and/or infected with either *E. coli* UTI89 or *E. faecalis* OG1RF.** C57BL/6 female WT (+/+) and coagulation-transgenic mice bladders were uninfected or infected with ~$10^7$ CFUs of *E. coli* UTI89 or *E. faecalis* OG1RF in absence (uUTI) or presence of a catheter (CAUTI) for 24 hours. Coagulation-transgenic mice used are: $Fg^{AEK}$ (soluble Fg, no fibrin formation), $Pg^{-/-}$ and $uPA^{-/-}$ (elevated fibrin accumulation, no plasmin activation), and PAI-$1^{-/-}$ (continuous fibrin clot degradation). Naïve (N) mice were neither catheterized nor infected. Bladders were then harvested and digested to isolate single cells before incubation with CD16/32 antibodies for blocking and viability dye. For flow cytometry analysis, cells were then stained with conjugated anti-mouse primary antibodies for CD45, CD11b, F4/80, iNOS, and Arginase-1. (**A**) All Mφs ($CD45^+CD11b^+F4/80^+$) were gated from live single cell populations before further gating for (**B**) M0 ($Arg1^-iNOS^-$), (**C**) M1 ($Arg1^-iNOS^+$), (**D**) M1/M2 ($Arg1^+iNOS^+$), and (**E**) M2 ($Arg1^+iNOS^-$) populations. Values represent median. Differences between groups were tested for significance using the Mann-Whitney U test. Statistical significance between each group and naïve shown as brown brackets on top of each graph. *, $P \leq 0.05$; **, $P \leq 0.005$; ***, $P \leq 0.0005$; ****, $P \leq 0.0001$.

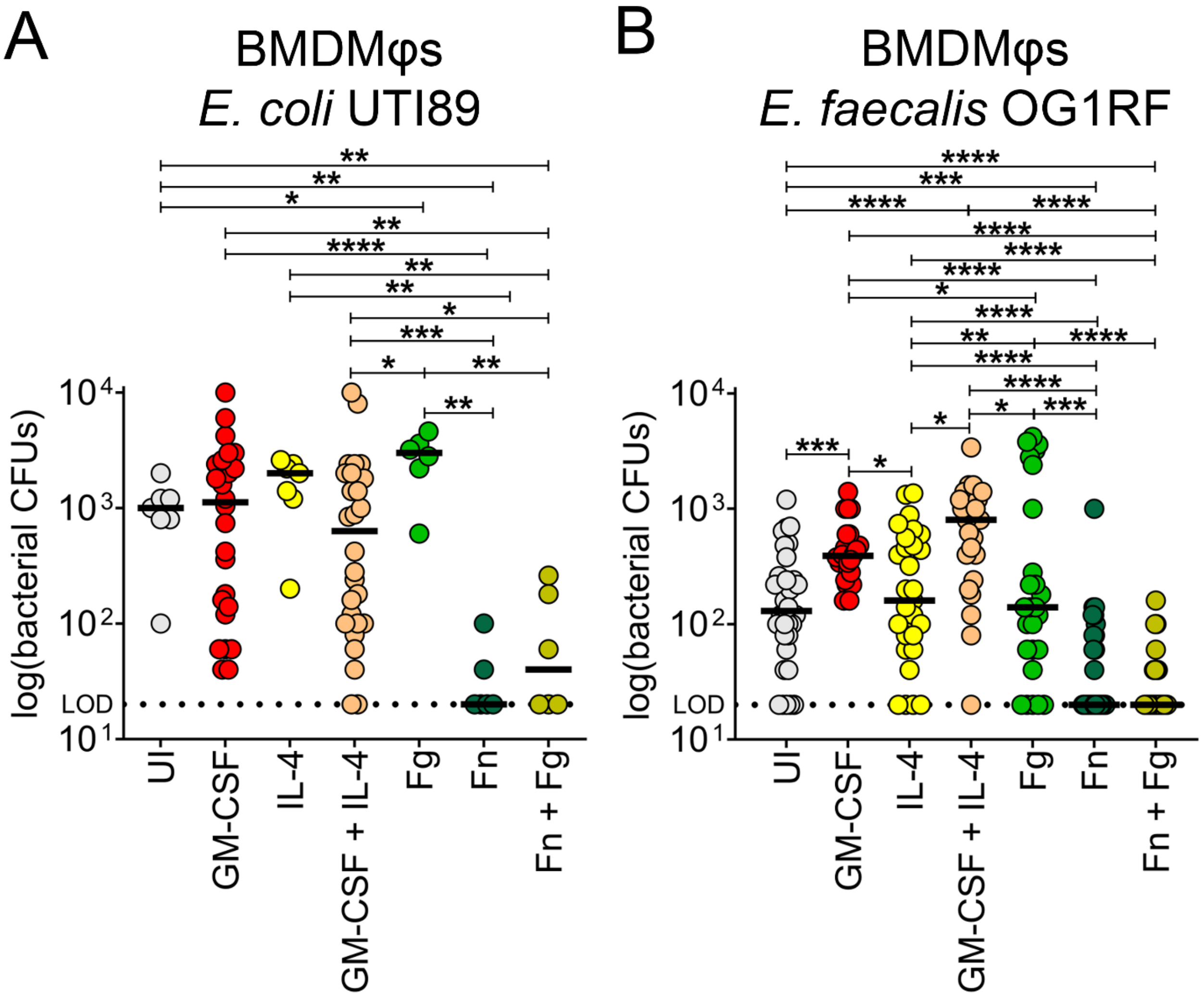


**Supplementary Figure 11. Fibrin suppressed Mφ phagocytosis of bacteria**. BMDMφs were either uninduced or stimulated with 100 ng/mL of GM-CSF, IL-4, or cotreated with GM-CSF+IL-4, 1.5 mg/mL of Fg, fibrin, or both simultaneously (fibrin+ Fg) for 24 hours. Mϕs were then infected with either opsonized (**A**) *E. coli* UTI89 or (**B**) *E. faecalis* OG1RF at multiplicity-of-infection ratio of 1 (50,000 bacteria to 50,000 Mφs) for an initial 45-minute uptake. Supernatant was then removed and Mφs were treated with gentamicin for 30 minutes to remove non-internalized bacteria before being exposed to pure sterile distilled water for intracellular bacteria retrieval and CFU counts. Horizontal line represents median value. Each data point is one replicate.

The horizontal broken line represents the limit of detection (LOD) of viable bacteria. The Mann-Whitney U test was used to determine significant difference between time points; *, $P \leq 0.05$; **, $P \leq 0.005$; ***, $P \leq 0.0005$; ****, $P \leq 0.0001$.

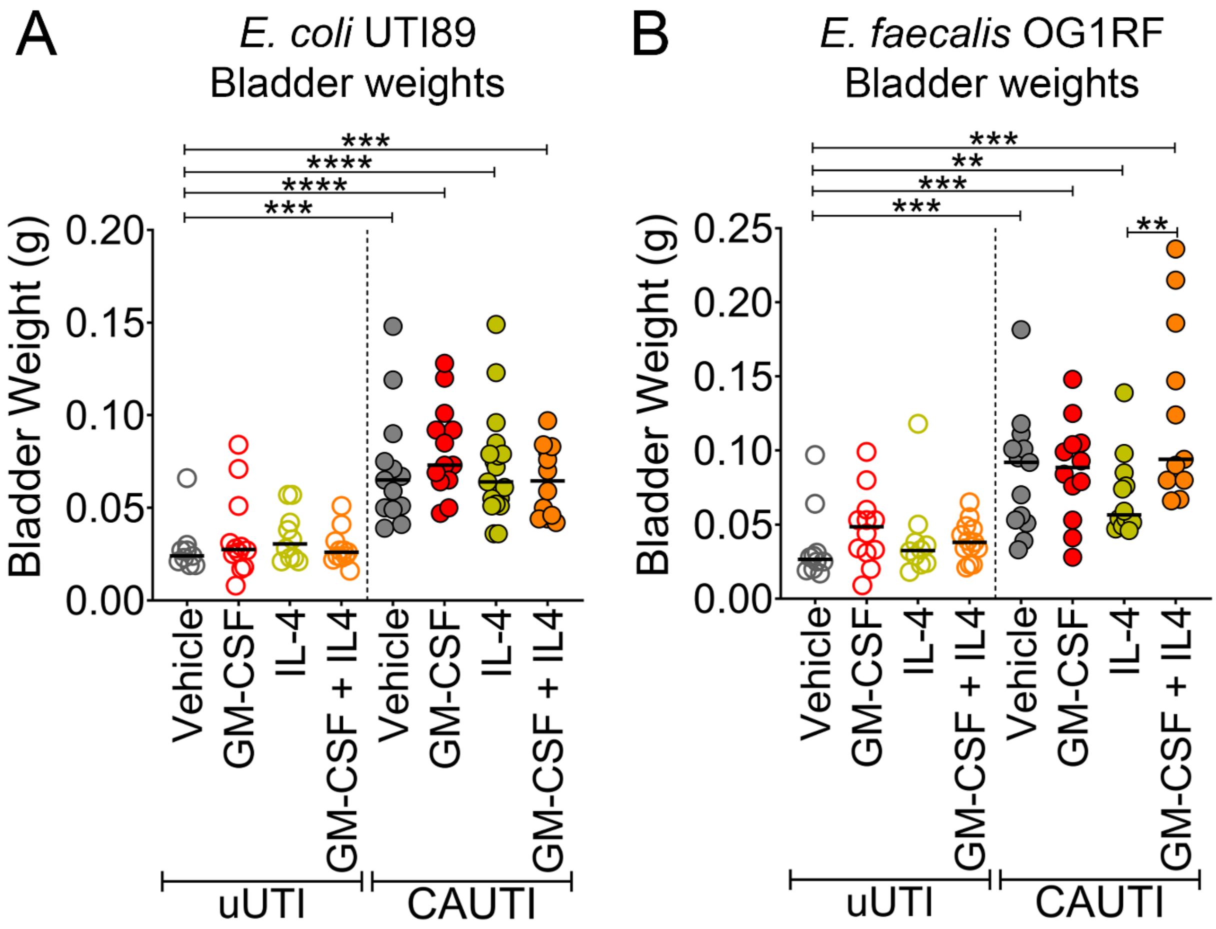


**Supplementary Figure 12**. **Catheterization promotes bladder inflammation regardless of inducer.** C57BL/6 WT mice were intraperitoneally (i.p.) injected with two doses of either sterile 1X PBS (vehicle), 200 ng/mouse of GM-CSF, IL-4, or co-treatment with GM-CSF and IL-4 at 12 and 4 hours prior to catheterization and/or infection with either (**A**) *E. coli* UTI89 or (**B**) *E. faecalis* OG1RF. Twelve hours post-catheterization and/or infection, bladders were harvested and weighted. The Mann-Whitney U test was used to determine significant difference between treatment; * **, $P \leq 0.005$; ***, $P \leq 0.0005$; ****, $P \leq 0.0001$.

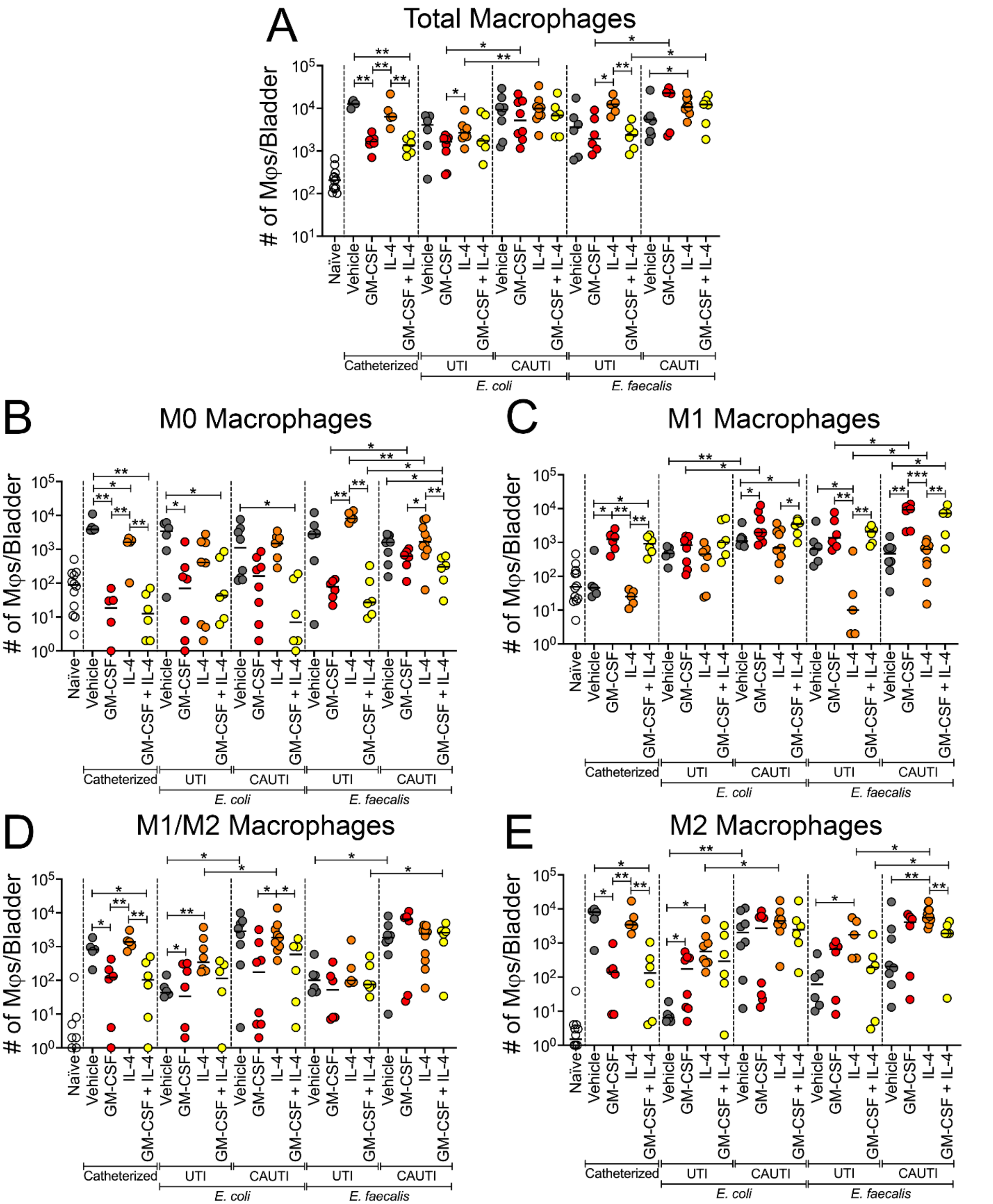


**Supplementary Figure 13**. **Total number of Mφs and polarization states in the bladder treated with different inducers before catheterization and/or infection with either *E. coli***

**UTI89 or *E. faecalis* OG1RF.** C57BL/6 WT mice were intraperitoneally (i.p.) injected with two doses of either sterile 1X PBS (vehicle), 200 ng/mouse of GM-CSF, IL-4, or co-treatment with GM-CSF and IL-4 at 12 and 4 hours prior to catheterization and/or infection with either *E. coli* UTI89 or *E. faecalis* OG1RF. Twelve hours post-catheterization and/or infection, single cells were isolated from the bladders. Cells were then stained with a viability dye and for conjugated antibodies for CD45, CD11b, F4/80, iNOS, and Arginase-1. (**A**) Total Mφs was gated from live single cell populations before further gating for (**B**) M0 (iNOS$^-$Arg1$^-$), (**C**) M1 (iNOS$^+$Arg1$^-$), (**D**) M1/M2 (iNOS$^+$Arg1$^+$), and (**E**) M2 (iNOS$^-$Arg1$^+$) populations. Values represent median. Differences between groups were tested for significance using the Mann-Whitney U test. Statistical significance between each group and naïve shown as brown brackets on top of each graph. *, $P \leq 0.05$; **, $P \leq 0.005$; ***, $P \leq 0.0005$; ****, $P \leq 0.0001$.

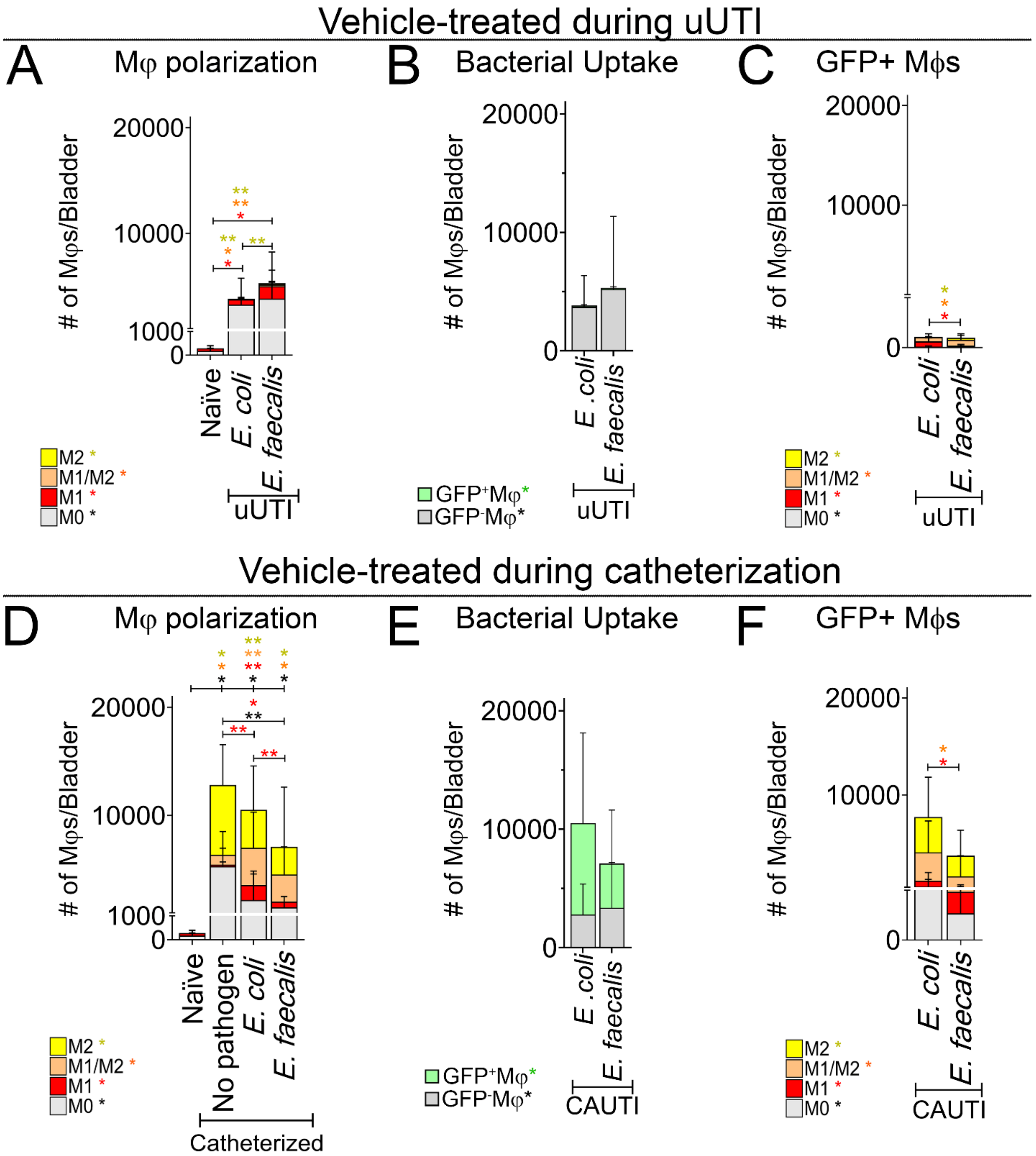


**Supplemental Figure 14. Comparison of Mφ polarization and bacterial uptake between vehicle-treated mice during uUTI and CAUTI.** C57BL/6 WT mice were i.p. injected with two doses of either sterile 1X PBS (vehicle) at 12 and 4 hours prior to catheterization and/or infection as described in **Fig. 5A**. Naïve (N) mice were neither catheterized nor infected. Mice were either infected with $10^8$ CFUs of *E. coli* UTI89-GFP strain or with $10^8$ CFUs of *E. faecalis* OG1RF-GFP strain in absence (uUTI) (**A-C**) or presence of a catheter (CAUTI) (**D-F**). Twelve hours post-catheterization and/or infection, bladders were harvested and digested to isolate single cells for flow cytometry analysis and staining as described in **Fig. 5A**. (**A, D**) Mφs polarization was evaluated. (**B, E**) $GFP^+$ Mφs (phagocytized the pathogen, $UPEC^+$ or $OG1RF^+$) and $GFP^-$ Mφs (did not phagocytize the pathogen, $UPEC^-$ or $OG1RF^-$) were each gated (**fig. S15, gating strategy**). (**C-F)** Mφs polarization was evaluated on $GFP^+$ Mφs populations phagocytizing the pathogen**.** The Mann-Whitney U test was used to test for significance between groups. *, $P \leq 0.05$; **, $P \leq 0.005$. The horizontal broken line represents the limit of detection (LOD) of viable bacteria. Three to four independent experiments were performed with 6-16 mice for each experimental group. Values represent mean ± SD.

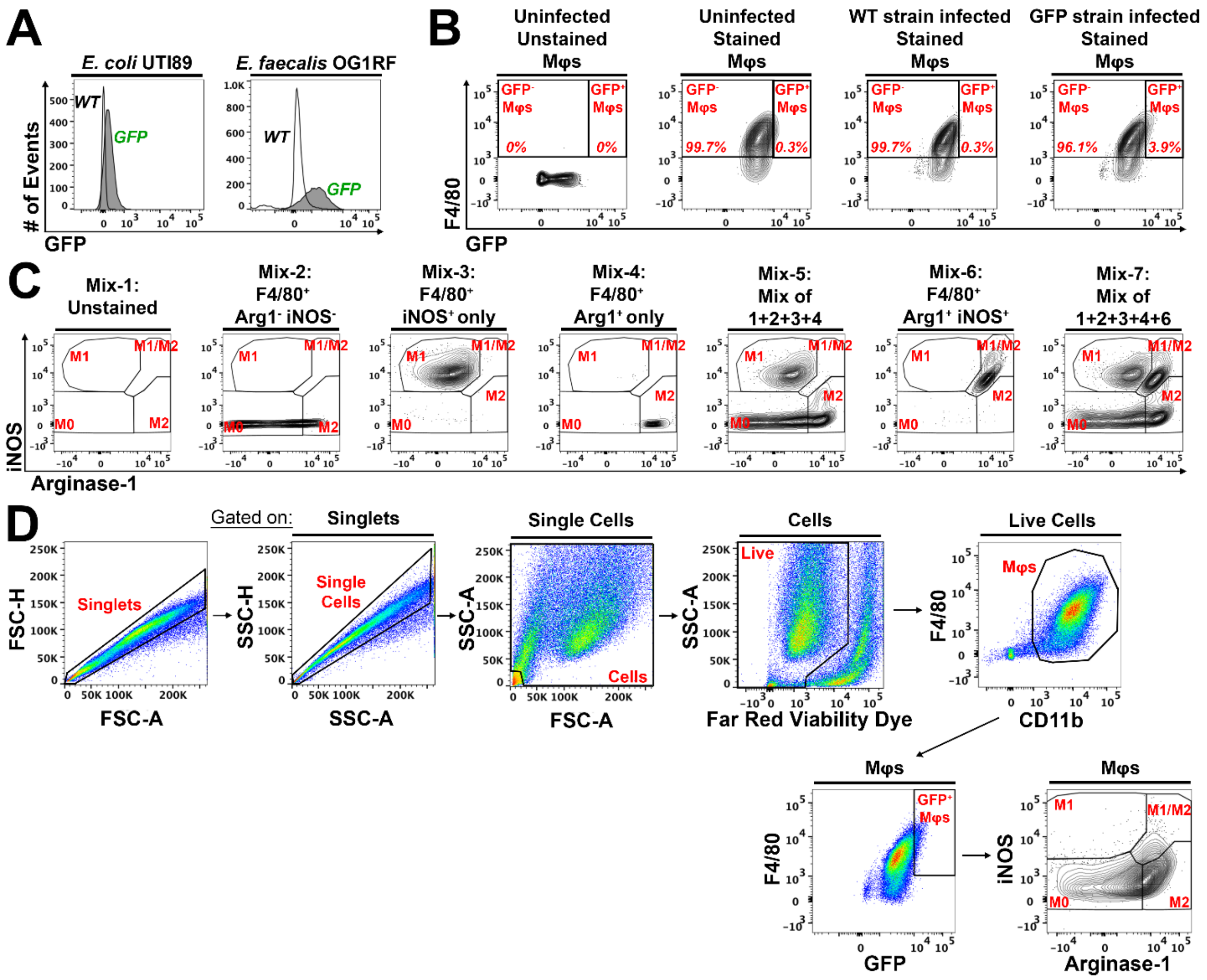


**Supplemental Figure 15. Representative gating strategy to determine macrophage phagocytotic activity. (A)** Flow cytometry analysis of *E. coli* UTI89 and *E. faecalis* OG1RF wildtype (WT) strain or GFP strain. **(B)** Gating strategy to determine positive signal cutoff of Mφs that phagocytosed the pathogen containing the induced *gfp* gene (GFP$^+$ Mφs) and Mϕs that did not engulf the pathogen (GFP$^-$ Mφs). RAW 264.7 Mϕs were uninfected or infected with either the *E. coli* UTI89 or *E. faecalis* OG1RF WT or GFP strain. **(C)** Gating strategy to determine M0 (Arg1$^-$ iNOS$^-$), M1 (Arg1$^-$ iNOS$^+$), M1/M2 (Arg1$^+$ iNOS$^+$), and M2 (Arg1$^+$ iNOS$^-$) Mφs. Cells in Mix-1 were unstained, while Mix-2 thru -4 & -6 were stained for both far-red viability dye and F4/80 Mφ marker. Cells in Mix-3 were additionally stained for iNOS, Mix-4 for Arg1, and Mix-6 for both iNOS and Arg1. Some part of the cell populations from Mix-1, -2, -3, and -4 were combined into Mix-5 and populations from Mix-1, -2, -3, -4, & -6 were combined into Mix-7 to distinguish gating between the 4 different populations. **(D)** Using the gating strategy determined in **B**, $GFP^+$ and $GFP^-$ Mϕs were subgated from the preceding bladder Mφ population. Using the gating strategy determined in **C**, M0, M1, M1/M2, and M2 Mφs were subgated from the preceding bladder Mφ population.

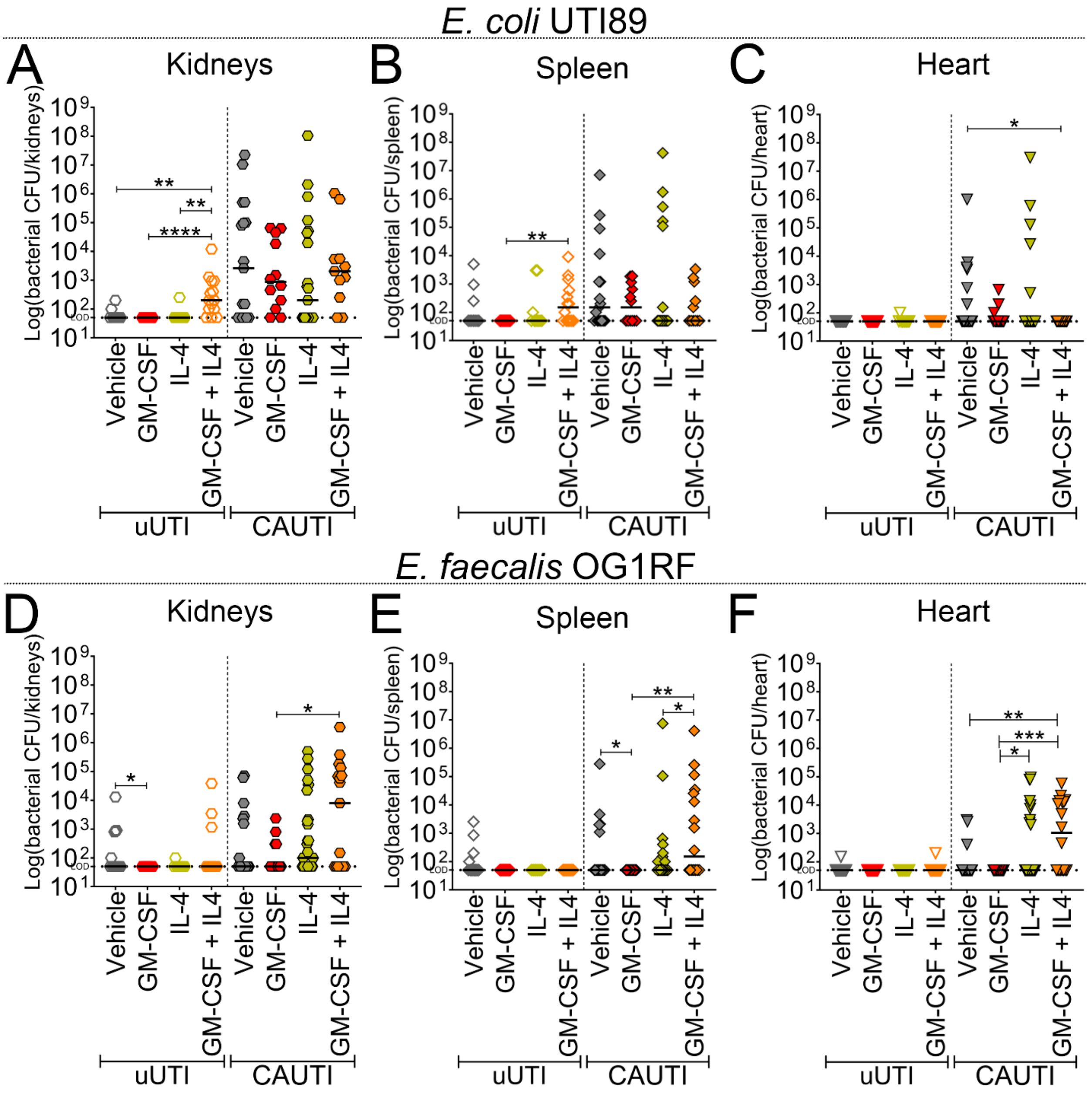


**Supplementary Figure 16**. **Bacterial colonization of the kidneys, spleen, and heart during uUTI and CAUTI.** C57BL/6 WT mice were i.p. injected with two doses of either sterile 1X PBS (vehicle) or cytokines at 12 and 4 hours prior to catheterization and/or infection as described in **Fig. 5A**. Naïve (N) mice were neither catheterized nor infected. Mice were either (**A-C**) infected with $10^8$ CFUs of *E. coli* UTI89 GFP strain or (**D-F**) infected with $10^8$ CFUs of *E. faecalis* OG1RF GFP strain in absence (uUTI) or presence of a catheter (CAUTI). Twelve hours post-catheterization and/or infection, the bacterial burden of each strain was determined by CFU count of (**A, D**) kidneys, (**B, E**) spleen, and (**C, F**) heart. Horizontal line represents median value. Each data point is one mouse. The Mann-Whitney U test was used to test for significance between groups. *, $P \leq 0.05$; **, $P \leq 0.005$; ***, $P \leq 0.0005$; ****, $P \leq 0.0001$. The horizontal broken line represents the limit of detection (LOD) of viable bacteria. Three to four independent experiments were performed with 6-16 mice for each experimental group.

#### SUPPLEMENTARY MATERIALS

**Supplementary Table 1. Number of Mφs in bladders of WT mice that were catheterized and/or infected with either *E. coli* UTI89 or *E. faecalis* OG1RF.** WT ($Fg^{+/+}$) are C57BL/6 female mice. Naïve mice are neither catheterized nor infected. Values in table are mean ± SD. Values are median displayed in **Figure 1B**.

| Condition | N (#) | # of Mφs / Bladder |
|---|---|---|
| Naïve | 22 | 665 ± 454 |
| Catheterized without infection | 9 | 21332 ± 19494 |
| UTI UPEC UTI89 | 10 | 3721 ± 2517 |
| CAUTI UPEC UTI89 | 10 | 14960 ± 15953 |
| UTI *E. faecalis* OG1RF | 9 | 1182 ± 1000 |
| CAUTI *E. faecalis* OG1RF | 10 | 19166 ± 28613 |

**Supplementary Table 2. Percentage of BMDMφs in each condition that are M0 (iNOS$^{-}$ Arg1$^{-}$), M1 (iNOS$^{+}$ Arg1$^{-}$), M1/M2 (iNOS$^{+}$ Arg1$^{+}$), and M2 (iNOS$^{-}$ Arg1$^{+}$).** Values in mean % ± SD.

| Condition | N | M0 (%) | M1 (%) | M1/M2 (%) | M2 (%) |
|---|---|---|---|---|---|
| Uninduced | 6 | 68.3 ± 7.2 | 3.1 ± 1.8 | 17.5 ± 2.1 | 11.1 ± 7.4 |
| GM-CSF | 6 | 27.7 ± 12.7 | 60.9 ± 7.2 | 6.2 ± 4.6 | 5.2 ± 2.5 |
| IFNγ + LPS | 6 | 31.8 ± 22.0 | 58.0 ± 20.2 | 5.0 ± 1.4 | 5.2 ± 4.4 |
| IL-4 | 6 | 23.6 ± 13.8 | 7.2 ± 6.9 | 4.4 ± 2.2 | 64.9 ± 12.0 |
| GM-CSF + IL-4 | 6 | 33.0 ± 9.5 | 54.3 ± 11.7 | 7.4 ± 5.3 | 5.4 ± 4.0 |
| IFNγ + LPS + IL-4 | 6 | 26.2 ± 14.3 | 18.3 ± 3.0 | 33.7 ± 4.4 | 21.8 ± 5.4 |
| Fibrinogen (Fg) | 12 | 30.8 ± 13.4 | 42.7 ± 25.4 | 18.7 ± 10.1 | 5.3 ± 3.8 |
| Fibrin (Fn) | 7 | 32.1 ± 24.4 | 9.2 ± 4.5 | 15.2 ± 9.5 | 43.4 ± 13.5 |
| Fn + Fg | 13 | 31.8 ± 24.1 | 4.0 ± 3.1 | 15.8 ± 15.3 | 48.4 ± 28.2 |

**Supplementary Table 3. Percentage of RAW 264.7 Mφs in each condition that are M0 (iNOS$^-$ Arg1$^-$), M1 (iNOS$^+$ Arg1$^-$), M1/M2 (iNOS$^+$ Arg1$^+$), and M2 (iNOS$^-$ Arg1$^+$).** Values in mean % ± SD.

| Condition | N | M0 (%) | M1 (%) | M1/M2 (%) | M2 (%) |
|---|---|---|---|---|---|
| Uninduced | 10 | 51.1 ± 13.8 | 13.8 ± 8.7 | 17.7 ± 6.0 | 17.4 ± 10.7 |
| GM-CSF | 5 | 17.9 ± 7.3 | 61.7 ± 9.1 | 14.8 ± 7.0 | 5.5 ± 2.6 |
| IFNγ + LPS | 10 | 34.1 ± 17.4 | 55.9 ± 16.8 | 4.5 ± 1.7 | 5.5 ± 4.0 |
| IL-4 | 10 | 23.1 ± 15.3 | 7.3 ± 6.2 | 4.9 ± 2.2 | 64.7 ± 13.2 |
| GM-CSF + IL-4 | 5 | 18.3 ± 7.3 | 29.2 ± 18.9 | 47.0 ± 20.8 | 5.5 ± 4.3 |
| IFNγ + LPS + IL-4 | 6 | 26.2 ± 14.3 | 18.3 ± 3.9 | 33.7 ± 4.4 | 21.8 ± 9.9 |
| Fibrinogen (Fg) | 11 | 17.6 ± 16.7 | 47.3 ± 17.1 | 27.4 ± 6.0 | 7.7 ± 5.5 |
| Fibrin (Fn) | 10 | 35.5 ± 21.4 | 8.7 ± 4.1 | 13.5 ± 8.6 | 42.4 ± 13.6 |
| Fn + Fg | 9 | 39.4 ± 15.6 | 9.1 ± 6.0 | 16.5 ± 8.1 | 35.1 ± 17.0 |

**Supplementary Table 4. Total number of Mφs in bladders of WT and coagulation-transgenic mice that were catheterized and/or infected with either *E. coli* UTI89 or *E. faecalis* OG1RF.** WT ($Fg^{+/+}$) are C57BL/6 female mice. Values in mean ± SD. Number of Mφs means and SD are rounded to nearest whole number.

| Condition | Infection | Genotype | N | # of Mφs |
|---|---|---|---|---|
| Naïve | No infection | WT ($Fg^{+/+}$) | 22 | 665 ± 454 |
| Catheterized | | WT ($Fg^{+/+}$) | 9 | 21332 ± 19494 |
| | | $Fg^{AEK}$ | 7 | 25635 ± 21478 |
| | | $Pg^{-/-}$ | 4 | 23154 ± 12803 |
| | | $uPA^{-/-}$ | 6 | 32955 ± 23881 |
| | | $PAI\text{-}1^{-/-}$ | 7 | 65241 ± 32942 |
| uUTI | *E. coli* UTI89 | WT ($Fg^{+/+}$) | 7 | 4574 ± 2468 |
| CAUTI | | WT ($Fg^{+/+}$) | 7 | 18842 ± 17606 |
| | | $Fg^{AEK}$ | 7 | 42723 ± 33144 |
| | | $Pg^{-/-}$ | 5 | 50495 ± 34976 |
| | | $uPA^{-/-}$ | 6 | 81527 ± 39931 |
| | | $PAI\text{-}1^{-/-}$ | 8 | 47774 ± 56864 |
| uUTI | *E. faecalis* OG1RF | WT ($Fg^{+/+}$) | 6 | 1552 ± 1049 |
| CAUTI | | WT ($Fg^{+/+}$) | 7 | 26063 ± 32284 |
| | | $Fg^{AEK}$ | 6 | 46555 ± 56144 |
| | | $Pg^{-/-}$ | 4 | 55624 ± 20252 |
| | | $uPA^{-/-}$ | 6 | 23870 ± 20985 |
| | | $PAI\text{-}1^{-/-}$ | 8 | 83183 ± 62071 |

**Supplementary Table 5. Number and percentage of Mφs that are M0, M1, M1/M2, and M2 in bladders of WT and coagulation-transgenic mice that were catheterized and/or infected with either *E. coli* UTI89 or *E. faecalis* OG1RF.** WT ($Fg^{+/+}$) are C57BL/6 female mice. Values in mean ± SD. Number of Mφs means and SD are rounded to nearest whole number.

| | | | | # of Mφs | | | | % Out of Mφs | | | |
|---|---|---|---|---|---|---|---|---|---|---|---|
| **Condition** | **Infection** | **Genotype** | **N** | **M0** | **M1** | **M1/M2** | **M2** | **M0 (%)** | **M1 (%)** | **M1/M2 (%)** | **M2 (%)** |
| Naïve | No infection | WT ($Fg^{+/+}$) | 22 | 199 ± 209 | 81 ± 246 | 75 ± 71 | 311 ± 347 | 31.5 ± 28.4 | 9.1 ± 15.5 | 14.6 ± 13.7 | 44.9 ± 26.5 |
| Catheterized | | WT ($Fg^{+/+}$) | 9 | 3213 ± 5110 | 1018 ± 1257 | 2962 ± 3586 | 14139 ± 15529 | 9.3 ± 9.7 | 7.9 ± 8.8 | 21.0 ± 24.1 | 61.8 ± 23.3 |
| | | $Fg^{AEK}$ | 7 | 4496 ± 6615 | 13686 ± 14369 | 4632 ± 2608 | 2822 ± 3163 | 13.8 ± 11.0 | 47.7 ± 10.7 | 27.2 ± 14.7 | 11.3 ± 10.0 |
| | | $Pg^{-/-}$ | 4 | 771 ± 667 | 1040 ± 1209 | 8269 ± 3941 | 13074 ± 8886 | 2.8 ± 1.5 | 6.8 ± 7.2 | 37.5 ± 5.7 | 52.9 ±11.3 |
| | | $uPA^{-/-}$ | 6 | 3966 ± 6240 | 3126 ± 3255 | 5141 ± 4954 | 20723 ± 16209 | 7.6 ± 9.3 | 12..4 ± 12.8 | 17.1 ± 11.3 | 63.0 ± 11.8 |
| | | PAI-$1^{-/-}$ | 7 | 13297 ± 9938 | 36346 ± 19644 | 9594 ± 5620 | 6005 ± 4734 | 18.4 ± 8.7 | 53.2 ± 8.9 | 14.6 ± 3.5 | 13.9 ± 15.8 |
| uUTI | *E. coli* UTI89 | WT ($Fg^{+/+}$) | 7 | 465 ± 651 | 2216 ± 1470 | 1195 ± 924 | 698 ± 1145 | 7.4 ± 7.9 | 51.5 ± 23.8 | 30.2 ± 22 | 11.0 ± 12.9 |
| CAUTI | | WT ($Fg^{+/+}$) | 7 | 372 ± 488 | 843 ± 897 | 6566 ± 5883 | 11061 ± 11770 | 4.2 ± 8.0 | 8.8 ± 9.6 | 37.0 ± 12.9 | 50.0 ± 20.3 |
| | | $Fg^{AEK}$ | 7 | 13162 ± 13947 | 22283 ± 18651 | 4016 ± 1710 | 3262 ± 4010 | 21.7 ± 17.1 | 46.3 ± 10.6 | 24.9 ± 24.1 | 7.2 ± 5.4 |
| | | $Pg^{-/-}$ | 5 | 3673 ± 2719 | 8053 ± 11315 | 13448 ± 7514 | 25322 ± 23629 | 10.1 ± 12.1 | 13.2 ± 12.1 | 30.9 ± 9.6 | 46.2 ± 17.5 |
| | | $uPA^{-/-}$ | 6 | 6398 ± 4790 | 11908 ± 21447 | 8569 ± 4085 | 54652 ± 22863 | 7 .1± 4.7 | 11.7 ± 15.1 | 10.7 ± 2.4 | 70.4 ± 12.6 |
| | | PAI-$1^{-/-}$ | 8 | 1208 ± 1992 | 25553 ± 35451 | 16896 ± 19590 | 4119 ± 4784 | 5.6 ± 7.2 | 49.1 ± 18.7 | 34.7 ± 13.3 | 10.6 ± 18.3 |
| uUTI | *E. faecalis* OG1RF | WT ($Fg^{+/+}$) | 6 | 40 ± 54 | 793 ± 705 | 644 ± 465 | 74 ± 95 | 1.7 ± 1.8 | 49.0 ± 14.9 | 45.3 ± 18.0 | 4.1 ± 2.6 |
| CAUTI | | WT ($Fg^{+/+}$) | 7 | 2964 ± 6153 | 2264 ± 1927 | 2428 ± 1329 | 18407 ± 25398 | 10.2 ± 12.9 | 14.3 ± 9.7 | 19.8 ± 15.8 | 55.6 ± 26.7 |
| | | $Fg^{AEK}$ | 6 | 13939 ± 15578 | 23863 ± 33402 | 4974 ± 6665 | 3779 ± 8244 | 24.4 ± 22.6 | 44.3 ± 9.3 | 24.5 ± 18.7 | 6.8 ± 5.6 |
| | | $Pg^{-/-}$ | 4 | 7243 ± 8825 | 513 ± 551 | 15276 ± 4232 | 32592 ± 9906 | 11.1 ± 9.4 | 0.9 ± 0.9 | 28.5 ± 7.1 | 59.3 ± 3.8 |
| | | $uPA^{-/-}$ | 6 | 2076 ± 2025 | 2053 ± 1639 | 2854 ± 2718 | 16887 ± 17695 | 8.5 ± 8.4 | 18.2 ± 15.6 | 13.0 ± 4.5 | 60.3 ± 14.5 |
| | | PAI-$1^{-/-}$ | 8 | 12256 ± 9388 | 37985 ± 36525 | 19998 ± 17090 | 12944 ± 14848 | 21.9 ± 19.1 | 37.8 ± 20.0 | 24.8 ± 13.4 | 15.5 ± 21.1 |

**Supplementary Table 6. Table of mouse strains used in this study.**

| Mouse strain | Genotype | Phenotype | References |
|---|---|---|---|
| Wildtype (WT) mice | $Fg^{+/+}$ | Presence of both Fg and fibrin | |
| Fg AEK mutant form mice | $Fg^{AEK}$ | Fibrinopeptide A cannot be cleaved by thrombin resulting in absence of fibrin polymer formation | Prasad et al., 2015 |
| Plasminogen (Pg)-deficient mice | $Pg^{-/-}$ | Impaired fibrinolysis resulting in elevated fibrin accumulation | Ploplis et al., 2005 |
| Urokinase plasminogen activator (uPA)-deficient mice | $uPA^{-/-}$ | Impaired activation of PG resulting in fibrinolysis deficiency leading to elevated fibrin accumulation | Carmeliet et al., 1994 |
| Plasminogen activator inhibitor-1 (PAI-1)-deficient mice | $PAI\text{-}1^{-/-}$ | Impaired inhibition of UPA activity resulting in continuous degradation of fibrin clots leading to elevated fibrin degradation products | Carmeliet et al., 2003 I/II |

**Supplementary Table 7. Total number of Mφs in bladders of WT mice treated with inducers before catheterization and/or infection with either *E. coli* UTI89 or *E. faecalis* OG1RF.** Values in mean ± SD. Number of Mφs' means and SD are rounded to nearest whole number.

| Condition | Infection | Inducer | N | # of Mφs |
|---|---|---|---|---|
| Naïve | None | None | 12 | 245 ± 167 |
| Catheterized only | None | Vehicle | 5 | 12830 ± 2087 |
| | | GM-CSF | 6 | 1693 ± 689 |
| | | IL-4 | 5 | 9220 ± 7180 |
| | | GM-CSF+IL-4 | 6 | 1447 ± 630 |
| uUTI | *E. coli* UTI89 | Vehicle | 6 | 3823 ± 2710 |
| | | GM-CSF | 8 | 1397 ± 802 |
| | | IL-4 | 8 | 3371 ± 2452 |
| | | GM-CSF+IL-4 | 6 | 3414 ± 3339 |
| CAUTI | | Vehicle | 8 | 10515 ± 9261 |
| | | GM-CSF | 8 | 8257 ± 7608 |
| | | IL-4 | 9 | 11330 ± 9159 |
| | | GM-CSF+IL-4 | 6 | 8517 ± 7655 |
| uUTI | *E. faecalis* OG1RF | Vehicle | 6 | 5309 ± 6240 |
| | | GM-CSF | 6 | 3423 ± 3289 |
| | | IL-4 | 5 | 12369 ± 5806 |
| | | GM-CSF+IL-4 | 6 | 2614 ± 1737 |
| CAUTI | | Vehicle | 8 | 7109 ± 8073 |
| | | GM-CSF | 6 | 17272 ± 11835 |
| | | IL-4 | 10 | 12376 ± 5357 |
| | | GM-CSF+IL-4 | 6 | 11039 ± 6874 |

**Supplementary Table 8. Number and percentage of Mφs that are M0, M1, M1/M2, or M2 in bladders of WT mice treated with inducers before catheterization and/or infection with either *E. coli* UTI89 or *E. faecalis* OG1RF.** Values in mean ± SD. Number of Mφs means and SD are rounded to nearest whole number.

| | | | | **# of Mφs** | | | | **% Out of Mφs** | | | |
|---|---|---|---|---|---|---|---|---|---|---|---|
| **Condition** | **Infection** | **Inducer** | N | **M0** | **M1** | **M1/M2** | **M2** | **M0 (%)** | **M1 (%)** | **M1/M2 (%)** | **M2 (%)** |
| Naïve | None | None | 12 | 122±141 | 106±130 | 12±35 | 5±11 | 53.9±38.9 | 40.0±35.8 | 1.7±3.6 | 4.3±11.5 |
| Catheterized only | None | Vehicle | 5 | 5319±3203 | 147±243 | 912±599 | 6452±3724 | 42.5±26.5 | 1.2±2.0 | 7.0±4.1 | 49.3±25.4 |
| | | GM-CSF | 6 | 23±27 | 1285±774 | 146±157 | 239±362 | 1.9±2.1 | 76.5±28.0 | 8.2±9.2 | 13.3±20.3 |
| | | IL-4 | 5 | 1454±786 | 26±13 | 1533±896 | 6207±6151 | 18.5±13.5 | 0.4±0.2 | 20.2±12.0 | 60.9±10.9 |
| | | GM-CSF+IL-4 | 6 | 25±28 | 970±536 | 176±201 | 277±399 | 1.7±2.0 | 68.2±27.5 | 10.6±8.8 | 19.4±24.2 |
| uUTI | *E. coli* UTI89 | Vehicle | 6 | 3267±2506 | 494±211 | 54±48 | 8±5 | 71.9±27.9 | 26.2±27.5 | 1.3±0.9 | 0.6±0.9 |
| | | GM-CSF | 8 | 281±567 | 798±582 | 110±140 | 209±217 | 26.9±32.4 | 55.8±20.3 | 5.5±6.8 | 11.8±10.6 |
| | | IL-4 | 8 | 853±1032 | 418±334 | 949±1251 | 1151±1575 | 27.3±34.3 | 20.2±21.1 | 22.7±18.1 | 29.9±23.5 |
| | | GM-CSF+IL-4 | 6 | 256±369 | 2077±2192 | 149±159 | 932±1299 | 19.2±28.4 | 59.5±23.4 | 4.2±5.6 | 17.0±14.7 |
| CAUTI | | Vehicle | 8 | 2179±2678 | 1402±1017 | 3433±3297 | 3501±4068 | 17.5±15.2 | 28.9±29.2 | 30.6±21.8 | 23.0±17.1 |
| | | GM-CSF | 8 | 260±311 | 4099±4202 | 655±1109 | 3243±3570 | 11.3±12.5 | 57.8±23.2 | 4.9±8.0 | 26.0±29.2 |
| | | IL-4 | 9 | 1315±1195 | 1305±1213 | 3220±4184 | 5490±5025 | 12.8±13.5 | 19.2±23.9 | 23.7±12.5 | 44.3±17.3 |
| | | GM-CSF+IL-4 | 6 | 57±83 | 3174±1511 | 651±688 | 4635±6366 | 2.5±3.9 | 51.0±23.2 | 5.8±5.2 | 40.6±22.9 |
| uUTI | *E. faecalis* OG1RF | Vehicle | 6 | 3847±4406 | 1165±1522 | 172±211 | 126±182 | 62.7±31.6 | 31.0±28.8 | 4.1±2.2 | 2.2±1.3 |
| | | GM-CSF | 6 | 77±43 | 2697±2827 | 100±133 | 548±450 | 3.1±1.8 | 76.9±20.4 | 2.3±2.3 | 17.6±19.1 |
| | | IL-4 | 5 | 9381±3192 | 119±243 | 411±643 | 2458±2358 | 80.2±15.4 | 0.6±1.1 | 2.5±2.7 | 16.7±13.3 |
| | | GM-CSF+IL-4 | 6 | 86±125 | 1926±934 | 173±191 | 427±684 | 3.3±3.7 | 80.6±13.6 | 6.1±4.7 | 10.0±11.7 |
| CAUTI | | Vehicle | 8 | 1507±992 | 532±467 | 2505±2554 | 2565±5518 | 29.4±21.8 | 17.5±29.4 | 13.9±15.8 | 39.2±22.0 |
| | | GM-CSF | 6 | 598±332 | 7909±4785 | 5269±4413 | 3496±2917 | 5.1±4.0 | 58.5±23.6 | 21.5±16.2 | 14.9±10.5 |
| | | IL-4 | 10 | 2869±2805 | 564±406 | 2060±1640 | 6883±4080 | 19.9±14.9 | 4.5±3.3 | 18.6±15.8 | 56.9±20.1 |
| | | GM-CSF+IL-4 | 6 | 323±234 | 6281±4501 | 2398±1648 | 2205±1447 | 3.4±2.4 | 56.1±25.2 | 21.9±17.9 | 18.7±11.7 |

**Supplementary Table 9. Number and percentage of Mφs that phagocytized pathogens in bladders of WT mice treated with inducers before catheterization and infection with either *E. coli* UTI89 or *E. faecalis* OG1RF.** GFP$^{+}$ are Mφs that phagocytized the pathogen and GFP$^{-}$ are Mφs that did not phagocytize the pathogen. Values in mean ± SD. Number of Mφs means and SD are rounded to nearest whole number.

| | | | | **# of Mφs** | | **% Out of Mφs** | |
|---|---|---|---|---|---|---|---|
| **Condition** | **Infection** | **Inducer** | **N** | **GFP+** | **GFP-** | **GFP+ (%)** | **GFP- (%)** |
| uUTI | *E. coli* UTI89 | Vehicle | 6 | 109 ± 74 | 3714 ± 2644 | 3.4 ± 1.1 | 96.6 ± 1.1 |
| | | GM-CSF | 8 | 422 ± 276 | 975 ± 635 | 32.0 ± 15.9 | 68.0 ± 15.9 |
| | | IL-4 | 8 | 1093 ± 2078 | 2278 ± 983 | 23.6 ± 22.1 | 76.4 ± 22.1 |
| | | GM-CSF+IL-4 | 6 | 949 ± 1006 | 2465 ± 2353 | 23.7 ± 11.3 | 76.3 ± 11.3 |
| CAUTI | | Vehicle | 8 | 7725 ± 7616 | 2790 ± 2577 | 66.4 ± 24.2 | 33.6 ± 24.2 |
| | | GM-CSF | 8 | 5170 ± 7099 | 3087 ± 2777 | 43.3 ± 30.0 | 56.7 ± 30.0 |
| | | IL-4 | 9 | 8063 ± 6598 | 3268 ± 3151 | 68.1 ± 22.8 | 31.9 ± 22.8 |
| | | GM-CSF+IL-4 | 6 | 5890 ± 7265 | 2627 ± 1923 | 63.5 ± 19.4 | 36.5 ± 19.4 |
| uUTI | *E. faecalis* OG1RF | Vehicle | 6 | 96 ± 98 | 5213 ± 6146 | 2.5 ± 1.2 | 97.5 ± 1.2 |
| | | GM-CSF | 6 | 1784 ± 2123 | 1639 ± 1191 | 37.5 ± 28.8 | 62.5 ± 28.8 |
| | | IL-4 | 5 | 738 ± 774 | 11631 ± 5094 | 4.9 ± 3.5 | 95.1 ± 3.5 |
| | | GM-CSF+IL-4 | 6 | 1380 ± 1198 | 1234 ± 717 | 41.5 ± 32.2 | 58.5 ± 32.2 |
| CAUTI | | Vehicle | 8 | 3747 ± 4509 | 3361 ± 3846 | 48.8 ± 26.3 | 51.2 ± 26.3 |
| | | GM-CSF | 6 | 10921 ± 8390 | 6351 ± 3664 | 48.4 ± 27.4 | 51.6 ± 27.4 |
| | | IL-4 | 10 | 7698 ± 5401 | 4679 ± 3693 | 61.5 ± 26.0 | 38.5 ± 26.0 |
| | | GM-CSF+IL-4 | 6 | 8333 ± 6078 | 2873 ± 1749 | 63.2 ± 29.3 | 36.8 ± 29.3 |

**Supplementary Table 10. Number and percentage of GFP+ Mφs that are M0, M1, M1/M2, or M2 in bladders of WT mice treated with inducers before catheterization and infection with either *E. coli* UTI89 or *E. faecalis* OG1RF.** Values in mean ± SD. Number of Mφs means and SD are rounded to nearest whole number.

| | | | | # of GFP+ Mφs | | | | % Out of GFP+ Mφs | | | |
|---|---|---|---|---|---|---|---|---|---|---|---|
| **Condition** | **Infection** | **Inducer** | N | **M0** | **M1** | **M1/M2** | **M2** | **M0 (%)** | **M1 (%)** | **M1/M2 (%)** | **M2 (%)** |
| uUTI | *E. coli* UTI89 | Vehicle | 6 | 11±8 | 54±33 | 39±37 | 6±3 | 8.3±4.8 | 50.3±11.7 | 29.9±17.9 | 11.5±16.8 |
| | | GM-CSF | 8 | 50±77 | 239±169 | 109±139 | 25±19 | 21.8±26.6 | 52.2±16.0 | 16.2±19.1 | 9.8±11.5 |
| | | IL-4 | 8 | 4±6 | 92±130 | 609±1122 | 387±992 | 3.5±7.8 | 19.8±21.9 | 63.2±24.4 | 13.5±14.8 |
| | | GM-CSF+IL-4 | 6 | 26±62 | 657±776 | 163±146 | 103±140 | 9.8±17.9 | 58.2±22.5 | 16.5±16.4 | 15.5±9.2 |
| CAUTI | | Vehicle | 8 | 543±767 | 643±836 | 2907±3242 | 3631±4101 | 5.4±5.1 | 24.2±30.4 | 42.8±27.2 | 27.6±21.3 |
| | | GM-CSF | 8 | 49±61 | 2419±4120 | 651±1106 | 2051±3021 | 13.3±18.7 | 52.1±24.0 | 10.0±17.8 | 24.7±25.9 |
| | | IL-4 | 9 | 607±639 | 602±675 | 2678±3172 | 4176±3735 | 8.2±10.0 | 12.7±13.8 | 34.3±18.3 | 44.7±19.7 |
| | | GM-CSF+IL-4 | 6 | 37±56 | 1771±782 | 651±688 | 3432±6157 | 2.4±3.6 | 49.1±22.6 | 10.1±9.6 | 38.4±23.6 |
| uUTI | *E. faecalis* OG1RF | Vehicle | 6 | 10±9 | 9±11 | 57±61 | 21±26 | 9.9±7.9 | 9.2±9.2 | 57.1±13.2 | 23.9±12.9 |
| | | GM-CSF | 6 | 1±1 | 1406±1849 | 98±135 | 279±247 | 6.3±15.3 | 54.4±32.0 | 14.1±18.2 | 25.2±23.9 |
| | | IL-4 | 5 | 14±8 | 2±1 | 158±120 | 564±647 | 3.1±1.6 | 0.4±0.3 | 33.9±20.5 | 62.6±21.9 |
| | | GM-CSF+IL-4 | 6 | 1±1 | 1137±983 | 156±205 | 86±76 | 6.6±10.7 | 62.2±33.9 | 11.6±11.3 | 19.6±20.9 |
| CAUTI | | Vehicle | 8 | 268±335 | 216±278 | 1153±2101 | 2110±2637 | 12.4±10.5 | 16.4±24.8 | 25.6±30.6 | 45.6±24.3 |
| | | GM-CSF | 6 | 32±45 | 3993±3002 | 5254±4419 | 1643±1538 | 5.9±11.4 | 49.4±22.3 | 32.5±24.5 | 12.2±6.5 |
| | | IL-4 | 10 | 699±927 | 308±354 | 1550±1284 | 5140±4807 | 6.4±5.9 | 3.3±3.8 | 30.9±26.5 | 59.4±22.3 |
| | | GM-CSF+IL-4 | 6 | 59±84 | 4653±3920 | 2164±1684 | 1458±982 | 2.9±5.7 | 53.3±22.4 | 24.3±11.8 | 19.5±16.2 |

**Supplementary Table 11. Number and percentage of GFP- Mφs that are M0, M1, M1/M2, or M2 in bladders of WT mice treated with inducers before catheterization and infection with either *E. coli* UTI89 or *E. faecalis* OG1RF.** Values in mean ± SD. Number of Mφs means and SD are rounded to nearest whole number.

| | | | | # of GFP- Mφs | | | | % Out of GFP- Mφs | | | |
|---|---|---|---|---|---|---|---|---|---|---|---|
| **Condition** | **Infection** | **Inducer** | N | **M0** | **M1** | **M1/M2** | **M2** | **M0 (%)** | **M1 (%)** | **M1/M2 (%)** | **M2 (%)** |
| uUTI | *E. coli* UTI89 | Vehicle | 6 | 3257±2500 | 440±184 | 15±12 | 3±3 | 70.9±35.6 | 12.5±9.2 | 0.4±0.2 | 0.5±1.0 |
| | | GM-CSF | 8 | 231±537 | 559±431 | 1±1 | 184±205 | 25.9±33.6 | 59.1±23.3 | 0.1±0.1 | 14.9±15.6 |
| | | IL-4 | 8 | 849±1029 | 326±236 | 340±410 | 763±686 | 29.9±34.5 | 20.9±22.9 | 12.5±15.9 | 36.6±29.2 |
| | | GM-CSF+IL-4 | 6 | 231±322 | 1421±1469 | 6±14 | 808±1176 | 23.0±33.5 | 57.3±26.0 | 0.0±0.0 | 19.7±20.3 |
| CAUTI | | Vehicle | 8 | 1636±2152 | 759±302 | 256±353 | 140±302 | 38.6±31.0 | 46.6±26.7 | 12.0±15.2 | 2.8±4.4 |
| | | GM-CSF | 8 | 211±251 | 1680±1386 | 4±5 | 1193±1717 | 13.5±13.4 | 62.4±24.7 | 0.1±0.2 | 24.0±33.7 |
| | | IL-4 | 9 | 708±823 | 703±699 | 952±1386 | 905±1710 | 32.8±30.5 | 31.6±29.8 | 10.4±13.9 | 25.2±26.9 |
| | | GM-CSF+IL-4 | 6 | 20±36 | 1403±1029 | 23±45 | 1181±1173 | 2.6±4.4 | 58.6±19.9 | 0.1±0.1 | 38.7±23.3 |
| uUTI | *E. faecalis* OG1RF | Vehicle | 6 | 3838±4401 | 1156±1519 | 115±152 | 105±157 | 63.9±32.2 | 31.6±29.8 | 2.8±2.0 | 1.6±1.1 |
| | | GM-CSF | 6 | 76±43 | 1291±1016 | 2±3 | 269±219 | 5.7±4.3 | 76.1±20.3 | 0.2±0.4 | 17.9±16.5 |
| | | IL-4 | 5 | 9367±3184 | 117±242 | 253±532 | 1894±1742 | 83.9±13.3 | 0.6±1.2 | 1.4±2.7 | 14.1±11.5 |
| | | GM-CSF+IL-4 | 6 | 87±123 | 789±105 | 17±29 | 341±646 | 5.2±4.6 | 75.9±25.1 | 1.7±2.7 | 17.2±23.1 |
| CAUTI | | Vehicle | 8 | 1239±878 | 315±505 | 363±658 | 1444±2716 | 53.9±37.9 | 15.0±30.6 | 5.7±7.7 | 25.4±29.9 |
| | | GM-CSF | 6 | 567±353 | 3916±2008 | 15±12 | 1853±1493 | 8.6±2.0 | 68.2±17.5 | 0.5±0.6 | 22.8±17.3 |
| | | IL-4 | 10 | 2170±2487 | 256±320 | 510±625 | 1743±1949 | 59.0±40.3 | 4.0±3.4 | 9.1±11.8 | 27.8±30.5 |
| | | GM-CSF+IL-4 | 6 | 264±244 | 1628±1074 | 234±562 | 747±766 | 8.1±4.4 | 58.5±24.6 | 12.9±31.0 | 20.5±14.6 |

**Supplementary Table 12.** Table of materials, reagents, antibodies, and dyes used in this study.

| Material/Reagent Type | Materials | Manufacturer/Source | Catalog Number |
|---|---|---|---|
| Bacterial strain | *Escherichia coli* UTI89 HK::GFP | [17]Andersen et al., 2022<br>[75]Guiton et al., 2012<br>[76]Bi et al., 2009 | N/A |
| Bacterial strain | *Escherichia coli* UTI89 | [77]Mulvey et al., 2001 | N/A |
| Bacterial strain | *Enterococcus faecalis* OG1RF | [78]Murray et al.,1993<br>[79]Xu et al., 2017 | N/A |
| Bacterial strain | *Enterococcus faecalis* OG1RF pAOJ20 (GFP) | [80]Lizier et al., 2010<br>[81]Monk et al., 2012<br>[82]Gaston et al, 2020 | N/A |
| Animals | C57BL/6 mice | Jackson Laboratory | 000664 |
| Animals | $FG^{AEK}$ | [83]Prasad et al., 2015 | N/A |
| Animals | $PG^{-/-}$ | [45]Ploplis et al., 2005 | N/A |
| Animals | $uPA^{-/-}$ | [84]Carmeliet et al., 1995 | N/A |
| Animals | $PAI^{-/-}$ | [46]Carmeliet et al., 1993 | N/A |
| Cell Lines | RAW 264.7 | ATCC | TIB-71 |
| Cell Lines | Bone marrow-derived macrophages | Isolated from femurs of C57BL/6 mice | N/A |
| Dye | LIVE/DEAD Fixable Far Red Dead Cell Stain Kit, for 633 or 635 nm exication | Invitrogen | L34973 |
| Dye | Hoechyst Dye | Thermo Scientific | 622249 |
| Antibodies | PE-Cy5 Rat Anti-mouse CD45 Monoclonal Antibody, Clone: 30-F11 | BD Pharmingen | 553082 |
| Antibodies | Brilliant Violet 650 Rat Anti-mouse CD11b Monoclonal Antibody, Clone: M1/70 | BioLegend | 101259 |
| Antibodies | Super Bright 780 Rat Anti-mouse F4/80 Monoclonal Antibody, Clone: BM8 | Invitrogen | 78-4801-82 |
| Antibodies | PE Rat Anti-mouse iNOS Monoclonal Antibody, Clone: CXNFT | Invitrogen | 12-5920-82 |
| Antibodies | eFluor 450 Rat Anti-mouse Arginase-1 Monoclonal Antibody, Clone: A1exF5 | Invitrogen | 48-3697-82 |

| | | | |
|---|---|---|---|
| Antibodies | F4/80 Monoclonal Unconjugated Antibody, Clone: BM8 | Invitrogen | 14-4801-82 |
| Antibodies | Purified Rat Anti-mouse CD16/32 Monoclonal Unconjugated Antibody, Clone: 93 | BioLegend | 101302 |
| Antibodies | iNOS Rabbit Anti-mouse Polyclonal Unconjugated Antibody | Invitrogen | PA1-036 |
| Antibodies | Arginase-1 Goat Anti-Mouse Polyclonal Unconjugated Antibody | Invitrogen | PA5-18684 |
| Antibodies | Goat Anti-Fibrinogen Polyclonal Unconjugated Antibody | Sigma-Aldrich | F8512-2ML |
| Antibodies | Rabbit Anti-beta actin Antibody | ThermoFisher Scientific | PA5-85271 |
| Antibodies | Goat Anti-beta actin Antibody | OriGene Technologies | AB0145-200 |
| Antibodies | Rabbit Anti-*E. coli* serotype O/K Polyclonal Antibody | Invitrogen | PA1-25636 |
| Antibodies | Rabbit Anti-Group D Streptococcus Polyclonal Antibody | Invitrogen | PA1-73120 |
| Antibodies | DyLight 488 Donkey anti-goat IgG (H+L) Cross-Adsorbed Secondary Antibody | ThermoFisher Scientific | SA5-10086 |
| Antibodies | DyLight 488 Donkey anti-rabbit IgG (H+L) Cross-Adsorbed Secondary Antibody | ThermoFisher Scientific | SA5-10038 |
| Antibodies | DyLight 550 Donkey anti-rat IgG (H+L) Cross-Adsorbed Secondary Antibody | ThermoFisher Scientific | SA5-10027 |
| Antibodies | DyLight 550 Donkey anti-rabbit IgG (H+L) Cross-Adsorbed Secondary Antibody | ThermoFisher Scientific | SA5-10039 |
| Antibodies | DyLight 650 Donkey anti-rabbit IgG (H+L) Cross-Adsorbed Secondary Antibody | ThermoFisher Scientific | SA5-10041 |

| | | | |
|---|---|---|---|
| Antibodies | DyLight 650 Donkey anti-goat IgG (H+L) Cross-Adsorbed Secondary Antibody | ThermoFisher Scientific | SA5-10089 |
| Antibodies | DyLight 650 Donkey anti-rat IgG (H+L) Cross-Adsorbed Secondary Antibody | ThermoFisher Scientific | SA5-10041 |
| Antibodies | IRDye680RD donkey anti-rabbit secondary antibody | LI-COR Bioscience | 926-68073 |
| Antibodies | IRDye680RD anti-goat secondary antibody | LI-COR Bioscience | 926-68074 |
| Antibodies | IRDye800CW donkey anti-goat secondary antibody | LI-COR Bioscience | 926-32214 |
| Antibodies | IRDye800CW donkey anti-rabbit secondary antibody | LI-COR Bioscience | 926-32213 |
| Recombinant Protein | Mouse Macrophage-colony Stimulating Factor (M-CSF) | R&D Systems | 416-ML-010 |
| Recombinant Protein | Mouse Granulocyte-macrophage colony-stimulating Factor (GM-CSF) | IrvineScientific | 200-15 |
| Recombinant Protein | Animal-Free murine interleukin-4 (IL-4) | PeproTech | AF-214-14 |
| Recombinant Protein | Lipopolysaccharides from *Escherichia coli* O111:B4, purified | Sigma-Aldrich | L3012-5MG |
| Recombinant Protein | Recombinant mouse IFN-gamma | R&D Systems | 485-MI-100 |
| Reagents | UltraComp eBeads Compensation Beads | Invitrogen | 01-2222-42 |
| Reagents | Liberase™ | Roche | 05401127001 |
| Reagents | DNaseI | ThermoFisher Scientific | EN0521 |
| Reagents | Human Fibrinogen | Enzyme Research Laboratories | FIB 3 |
| Reagents | Fibrinogen From Human Plasma, Alexa Fluor™ 647 Conjugate | Invitrogen | F35200 |
| Reagents | Human Thrombin | Sigma-Aldrich | T6884-250UN |
| Reagents | Phosphate Buffer Saline (PBS) | Sigma-Aldrich | P3813-10PAK |
| Reagents | Dulbecco's Phosphate Buffer Saline (DPBS) | Corning | 21-030-CM |
| Reagents | 10% Neutral Buffered Formalin | Leica | 3800600 |

| | | | |
|---|---|---|---|
| Reagents | Tween-20 | VWR | M147-1L |
| Reagents | Bovine Serum Albumin (BSA), for biochemistry, Nuclease-free, Protease-free | VWR | 97061-418 |
| Reagents | HEPES free acid | VWR | 97061-826 |
| Reagents | Roswell Park Memorial Institute (RPMI) 1640 with L-glutamine | Corning | 10-040-CM |
| Reagents | Dulbecco's Modified Eagle Medium (DMEM) with 4.5 g/L glucose and sodium pyruvate | Corning | 10-013-CV |
| Reagents | Luria Broth (LB) | MP Biomedicals | 3002-042 |
| Reagents | Brain Heart Infusion (BHI) | Hardy Diagnostics | C5143 |
| Reagents | Fusidic Acid | Chem-Impex | 26504 |
| Reagents | Rifampicin | VWR | TCR0079-25G |
| Reagents | Chloramphenicol | Sigma Life Science | C0378-25G |
| Reagents | Kanamycin | Santa Cruz Biotechnologies | sc-257635A |
| Reagents | Sodium Pyruvate (100 mM Solution) | Corning | 25-000-CI |
| Reagents | Fetal Bovine Serum (FBS) | ThermoFisher Scientific | 10-437-028 |
| Reagents | L-glutamine | VWR | 45000-404 |
| Reagents | Histopaque 1077 | Sigma-Aldrich | 10771-100 |
| Reagents | Histopaque 1119 | Sigma-Aldrich | 11191-100 |
| Reagents | Xylene | Sigma-Aldrich | 247642-4L |
| Reagents | Isopropanol | VWR | BDH1133-4LP |
| Reagents | Sodium Citrate | VWR | 0101-500G |
| Reagents | Triton X-100 | Acros Organics | 21568-2500 |
| Reagents | ProtoGel 30% (w/v) Acrylamide:0:8% (w/v) Bis-Acrylamide Stock Solution (37 .5:1) Protein and Sequencing Electrophoresis Grade | National Diagnostics | EC-890 |
| Reagents | Penicillin : Streptomycin solution, Corning® 100X | Corning | **30-002-CI** |
| Consumables | 30G ½" Needles | BD | 305106 |
| Consumables | Round-bottom 96-well plates | Corning | 3788 |
| Consumables | Flat-bottom 96-well plates | Greiner Bio-One | 655180 |
| Consumables | Round-bottom 96-well plates | Corning | 3788 |
| Consumables | CELLSTAR Tissue culture 25 $cm^2$, 75 $cm^2$, and 175 $cm^2$ flasks | VWR | 82050 |

| | | | |
|---|---|---|---|
| Consumables | BD Intramedic™ Polyethylene Tubing | BD Intramedic | 427401 |
| Consumables | RenaSil Silicone Rubber Tubing | Braintree Scientific | SIL 025 |
| Consumables | Polyvinylidene difluoride membrane (PVDF) | Millipore Sigma | IPFL00005 |
| Equipment | Zeiss Axio Observer | Zeiss | N/A |
| Equipment | Apotome | Zeiss | N/A |
| Equipment | Sonicator | Branson | 2800 |
| Equipment | Spectramax ABS plus | Molecular Devices | ABSPLUS |
| Equipment | BD Fortessa Flow Cytometer | BD Biosciences | N/A |
| Equipment | TransBlot Turbo Semi-dry transfer equipment | Bio-Rad | N/A |
| Software/Programs | ImageJ / Fiji | Schnieder et al. | N/A |
| Software/Programs | Prism 9 | GraphPad | N/A |
| Software/Programs | ZenPro | Zeiss | N/A |
| Software/Programs | FlowJO | FlowJo LLC | N/A |
